# Drivers of spatio-temporal variability in a marine foundation species

**DOI:** 10.1101/2024.06.25.600483

**Authors:** Anita Giraldo-Ospina, Tom Bell, Mark H. Carr, Jennifer E. Caselle

## Abstract

Marine foundation species are critical for the structure and functioning of ecosystems and constitute the pillar of trophic chains while also providing a variety of ecosystem services. In recent decades many foundation species have declined in abundance, sometimes threatening their current geographical distribution. Kelps (Laminariales) are the primary foundation species in temperate coastal systems worldwide. Kelp ecosystems are notoriously variable and identifying the key factors that control the dynamics of kelp abundance is key to predicting the fate of kelp ecosystems under climatic change and informing management and conservation decisions such as forest restoration. Here, we used *in situ* data from long-term monitoring programs across 1,350 km of coast spanning multiple biogeographic regions in the state of California (USA) to identify the major regional drivers of density of two dominant canopy-forming kelp species and to elucidate the spatial and temporal scales over which they operate. We used generalized additive models to identify the key drivers of density of two dominant kelp species (*Nereocystis luetkeana* and *Macrocystis pyrifera*) across four ecological regions of the state of California (north, central, south-west and south-east) and for the past two decades (2004-2021). Our study identified that the dominant drivers of kelp density varied between regions and species but always included some combination of nitrate availability, wave energy and exposure, density of purple sea urchins, and temperature as the most important predictors explaining 63% of the variability of bull kelp in the north and central regions, and 45% and 51.4% of the variability in giant kelp for the central/south-west and south-east regions, respectively. These large-scale analyses infer that a combination of lower nutrient availability, changes in wave energy and exposure, and increases in temperature and purple sea urchin counts have contributed to the decline of kelp observed in the last decade. Understanding the drivers of kelp dynamics can be used to identify regions and periods of significant change and historical stability, ultimately informing resource management and conservation decisions such as site selection for kelp protection and restoration.

**Open research statement:** Data (Giraldo-Ospina et al. 2023) are available in DataOne at doi:10.25494/P6/When_where_and_how_kelp_restoration_guidebook_2.

## Introduction

Of the many species whose distributions and dynamics are exhibiting dramatic changes in response to a changing global climate, few are as important and concerning as foundation species (*sensu* Dayton 1972, Ellison et al. 2005). These species support ecosystem productivity, create structural habitat, act as ecological engineers (*sensu* Jones et al. 1994), and underpin a multitude of ecosystem services. In coastal marine ecosystems, foundation species such as corals (Hughes et al. 2003, Hoegh-Guldberg et al. 2007, De’ath et al. 2012), seagrasses (Short and Neckles 1999, Koch et al. 2013, Serrano et al. 2021), mangroves (Alongi 2015, Ward et al. 2016), and oysters (Beck et al. 2011), among others, are experiencing marked changes in distribution and dynamics in response to gradual and episodic changes in water temperature (e.g., marine heatwaves) and other anthropogenic stressors. These spatial and temporal responses in such population attributes as abundance, productivity, and demographic and genetic structure are complex because of the interactions among multiple simultaneously changing environmental and ecological drivers. This complexity is compounded by the many spatial (local, regional, global) and temporal (seasonal, interannual, decadal) scales over which environmental conditions vary and interactions among scale-specific sources of variation. Consequently, one of the most challenging goals in ecology and conservation biology is to elucidate the spatial and temporal relationships between species abundance and variable environmental and ecological conditions. Such relationships are central to explaining variability in their distribution and predicting the dynamics of foundation species abundance and the ecosystems they create. In light of the accelerating effects of climate change, there is an urgent need to identify the key drivers of geographical and temporal variation in foundation species in order to make effective management decisions that protect them and the ecosystem services they provide.

Globally, kelps (Laminariales) are among the most important foundation species in temperate coastal oceans, inhabiting shallow temperate rocky reefs throughout the world (Steneck et al. 2002, Graham et al. 2007, Bolton 2010, Assis et al. 2020, Eger et al. 2023). Like many other foundation species, their primary production both fuels and creates physical habitat structures for highly species-rich ecosystems. These productive ecosystems support a multitude of culturally and economically important fisheries species, among other provisioning, regulating, supporting, and cultural services (Vásquez et al. 2014, Eger et al. 2023). With declines of kelp forests in many parts of the world associated with gradual and episodic (marine heatwaves) increases in water temperature and other anthropogenic stressors (Krumhansl et al. 2016, Wernberg et al. 2016, Beas-Luna et al. 2020, Arafeh-Dalmau et al. 2021b), there is global interest in conservation and restoration of kelps and their associated ecosystems (Morris et al. 2020, Eger et al. 2022).

From Baja California, Mexico to southeast Alaska, USA, the dominant canopy-forming kelps are the bull kelp (*Nereocystis luetkeana*) and the giant kelp (*Macrocystis pyrifera*), both of which create large, floating canopies (Graham et al. 2007, Schiel and Foster 2015, Carr and Reed 2016). Over the past decade, both bull and giant kelp forests on the West coast of North America experienced two major disturbances: the 2013 sea star wasting disease that led to local extinction of a key sea urchin predator, the sunflower star (*Pycnopodia helianthoides*), and the North East Pacific marine heatwave of 2014-2016 (NE Pacific MHW)(Michaud et al. 2022). These events were rapidly followed by extensive loss (> 90%) of bull kelp along the coast of northern California (Rogers-Bennett and Catton 2019, McPherson et al. 2021) and substantial losses of giant kelp in central California (Smith et al. 2021, 2024) and Baja California, Mexico (Arafeh-dalmau et al. 2019, Beas-Luna et al. 2020). The loss of kelp in northern California drove closures of important recreational and commercial fisheries such as red abalone and red sea urchins, and critically, almost a decade later, these lost forests have yet to recover (Rogers-Bennett and Catton 2019, McPherson et al. 2021). The detrimental consequences of the widespread loss of kelp have given rise to an urgent need to better understand the drivers of kelp dynamics in order to optimize decisions about conservation and restoration of this important marine habitat.

In California, the geographic distributions of these two kelp species extend across two well recognized biogeographic regions distinguished by persistent differences in oceanographic conditions (e.g. water temperature, wave energy, and coastal upwelling) (Briggs 1974, Horn et al. 2006, Blanchette et al. 2008, Reed et al. 2011). This environmental variability generates geographic differences in the structure and dynamics of kelp forest communities (Carr and Reed 2016), making California an ideal place to evaluate the drivers of climate-induced losses, gains, and distributional shifts.

To further advance our understanding of the environmental and ecological variables that explain the distribution and dynamics of canopy-forming kelps, including observed changes before and after the NE Pacific MHW, we ask (1) What environmental and biological variables best explain and predict the density dynamics of bull kelp and giant kelp along the coast of California? (2) How do the relative and combined impact of these variables on the spatio–temporal dynamics of kelp density vary across the bioregions of the California coast? and (3) How well can these variables explain and predict regional dynamics of kelp (canopy) abundance detected by satellite imagery?

To answer these questions, we leverage a long-term (~18 years) dataset of biological observations and remotely sensed environmental time series to construct species distribution models (SDMs) of the densities of the two canopy-forming kelp species (bull and giant kelp). We use the SDMs to extrapolate kelp distributions along the coast of California and apply these annual predictions to construct a coast-wide time series of interannual kelp density dynamics. We then compare the modeled regional kelp density dynamics with Landsat-derived spatio-temporal kelp canopy dynamics to evaluate how well our model explains patterns observed from satellite imagery. We used these results to inform resource managers about which sites are more likely to support dense forests in the face of future MHWs, and this knowledge can be used to prioritize sites for protection and restoration while considering the broad range of responses across a large geographical area (Giraldo-Ospina et al. 2023a).

## Materials and Methods

### Study system

#### Study species

Bull kelp (*Nereocystis luetkeana*) is an annual species with high interannual variation in forest density and area (McPherson et al. 2021). In California, bull kelp is distributed from the Oregon border in the north to Point Conception in the south. North of Monterey Bay, central California, it is the dominant habitat-forming kelp, whereas in central California bull kelp usually grows in mixed beds with giant kelp. Individuals are characterized by a single long stipe up to 25 m in length that extends through the water column from the rocky reef surface, buoyed by a large buoyant pneumatocyst. Long blades attached to the pneumatocyst contain spore-filled sori. Haploid zoospores released from the adult sporophytes in fall, settle to the reef and grow into microscopic gametophytes, which release gametes, fertilize, and develop into young sporophytes during winter. Young sporophytes quickly grow into adults over spring, forming a surface canopy in late summer and fall, which is typically dislodged by winter storms (Dobkowski et al. 2019).

Giant kelp (*Macrocystis pyrifera*) is a perennial species dominant in the temperate eastern Pacific and Southern Oceans. Giant kelp forests experience large temporal variation with biomass dynamics driven largely by the longevity of individual fronds rather than whole plants (Reed et al. 2008, Rodriguez et al. 2013). In California, giant kelp ranges predominantly from Pigeon Point in the north to the border with Mexico in the south (Graham et al. 2007, Schiel and Foster 2015, Carr and Reed 2016). Giant kelp abundance in California is very dynamic since individuals as well as entire forests are highly susceptible to dislodgment by ocean waves (Graham 1997, Edwards and Estes 2006). Because of this, forests are more persistent in the more protected waters of southern versus central California (Reed et al. 2011). Individual adult sporophytes are composed of numerous stipes that extend as much as 20-30 m through the water column from the rocky reef to the surface, each with many pneumatocysts and blades (Schiel and Foster 2015). Sporophylls at the base of the alga have the potential to produce zoospores throughout the year (Reed et al. 1997; Graham 1999). Like bull kelp, zoospores that settle to the reef produce gametophytes, which produce gametes that become young sporophytes upon fertilization.

#### Study region

This study focused on both bull kelp and giant kelp and encompassed the entire 1,350 km of coastal California, between the borders of Mexico to Oregon, including the offshore islands (Figure 1). The two species of kelp are distributed across two well-recognized biogeographic regions, each of which encompasses smaller “ecoregions” distinguished by persistent differences in ocean temperatures and species composition (Briggs 1974, Horn et al. 2006, Blanchette et al. 2008). In addition to persistent differences in oceanographic conditions, the coastal geomorphology varies among provinces and ecoregions. These geomorphological differences include the width of the continental shelf (and coastal upwelling), the exposure of the shoreline to ocean swell, the steepness of subtidal rocky reefs, turbidity, and the composition, vertical relief and rugosity of the rocky reef substratum (Hamilton et al. 2010; Blanchette et al. 2008). Separately and in combination, these oceanographic and geomorphological features generate persistent geographic patterns of the community structure and dynamics of kelp forest ecosystems (Carr and Reed 2016, Beas-Luna et al. 2020). For example, some subregions, such as the North-western Channel Islands, show evident separation from their ecoregion and such breaks coincide with known differences in oceanographic patterns (Blanchette et al. 2008, Hamilton et al. 2010). Based on these marked ecological and environmental differences, we divide the coastline into four regions to evaluate the determinants of kelp dynamics: 1) North, from the border with Oregon to Pigeon Point; 2) Central, from Pigeon Point to Point Conception; 3) South-west, from Point Conception to Naples beach including San Miguel and Santa Rosa Islands; and 4) South-east, from Naples beach to the border with Mexico (Figure 1).

**FIGURE 1.**
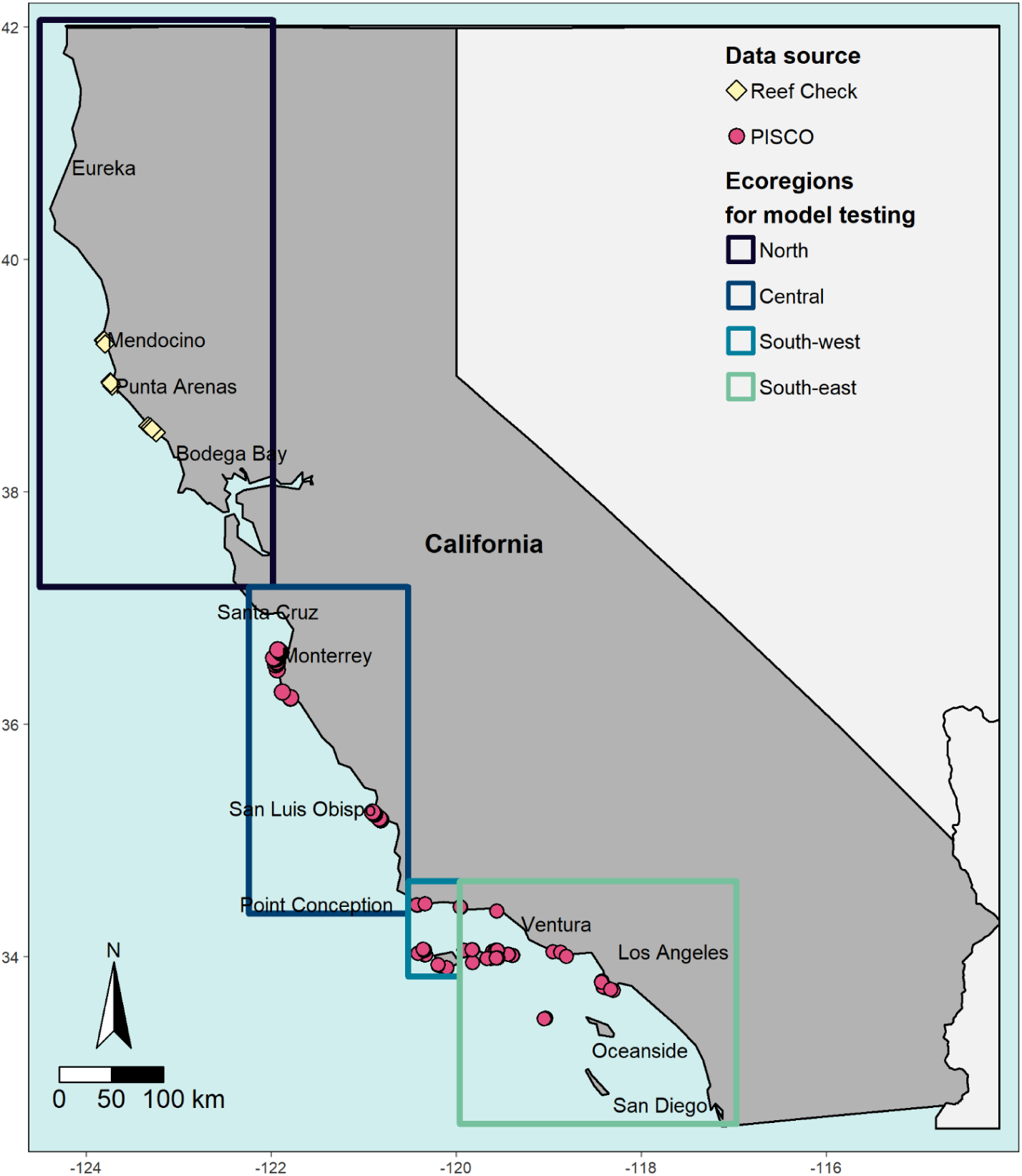
Study area along the coast of California and the four ecoregions that were used for modelling density of bull kelp (Northern and Central), and giant kelp (central, south-west and south-east). Symbols indicate the *in situ* survey sites used for modelling and their source (Reef Check vs. PISCO).

### Biological data from in situ surveys

Statewide data on kelp and grazer densities were obtained from kelp forest monitoring programs initiated by the Partnership for Interdisciplinary Studies of Coastal Oceans (PISCO; Carr et al. 2020, Malone et al. 2022) and Reef Check (https://www.reefcheck.org/country/usa-california/) and more recently conducted by a consortium of institutions that survey kelp forest ecosystems in California’s statewide network of marine protected areas (MPAs): UC Santa Barbara, UC Santa Cruz, California State Polytechnic University, Humboldt, and the Vantuna Research Group at Occidental College (VRG). Reef Check California conducts kelp monitoring surveys by training and leading citizen scientists across the state using methods comparable to PISCO. Densities of the giant kelp, purple sea urchin (*Strongylocentrotus purpuratus*) for the Central region (PISCO-UCSC) and two Southern regions (PISCO-UCSB and VRG) of California were obtained by visual surveys using scuba conducted at 95 sampling sites (Figure 1). Surveys were conducted from 1999 to 2021 using two (Central region) and three (Southern regions) transects of 30 m in length and 2 m width (60 m^2^) in three depth zones: inner (~ 5 m), mid (~12 m) and outer (~20 m) at each site, for a total of six (Central region) and nine (Southern regions) transects per site. Because the resolution of most environmental variables was much coarser than the scale of a transect or site, we used a central coordinate in the center of each site to extract the environmental data for each site. For the spatio-temporal modeling described below, only sites that contained three years of survey data before the NE Pacific MHW were used (N= 95 sites).

We used Reef Check data for the Northern region models as PISCO data were limited before the NE Pacific MHW. Reef Check California surveyed 25 sites in Sonoma and Mendocino Counties between 2006 and 2021, ranging between one and 21 sites annually. Two depths were surveyed at each site (inner: 0-10 m, and outer: 10-20 m) using three consecutive transects, similar to PISCO, for a total of six transects per surveyed site, per year. For the spatio-temporal modeling described below, only sites that contained three years of survey data prior to NE Pacific MHW were used (N= 10 sites).

### Environmental datasets of predictor variables

#### Satellite derived environmental variables

We obtained data for a comprehensive suite of environmental variables previously identified as correlates with bull and giant kelp distribution and dynamics (Bell et al. 2015). All temperature data and derived metrics were produced from the Daily Global 5 km Satellite Sea Surface Temperature dataset, available from NOAA Coral Reef Watch (https://coralreefwatch.noaa.gov/product/5km/index_5km_sst.php). The CoralTemp SST product combines a series of datasets to produce a timeseries from 1985 to present of daily global night-only SST at 5 km resolution. Daily SST data were used to calculate 44 annual and seasonal (summer vs. upwelling) metrics of SST that included general statistics (mean, max, min SST) and marine heatwave estimates (degree days and anomalies; see Appendix S1: Table S1 for full list of metrics). Surface nitrate concentrations were estimated using the spatial SST datasets. Surface nitrate concentrations for the Central and two Southern regions (< 37.75 N) were calculated using the observed relationship between ocean temperature and nitrate concentration as per (Snyder et al. 2020). The observed relationship between nitrate concentration and SST is different north of 37.75 N, so to estimate nitrate concentrations from SST for the Northern region, we used CalCOFI data (https://calcofi.org/data/oceanographic-data/) to develop a novel relationship using a generalized additive model. The resulting relationship was similar to the one presented in (García-Reyes et al. 2014). Using the daily spatial SST dataset, a suite of 41 metrics of nitrate concentrations similar to the SST metrics were determined at 5 km resolution (See Appendix S1: Table S1 for full list of metrics).

Wave height observations were obtained from the Coastal Data Information Program (CDIP; http://cdip.ucsd.edu/) MOPv1.1 wave model on an hourly timescale at 1 km coastline segments for the entire coast of California. All wave data prior to 2004 were hindcasted by developing a non-linear statistical model (GAM) between CDIP data and data from one of 18 offshore US Army Corp Wave Information Study (WIS) model sites. The site that produced the best model estimating CDIP wave height from WIS wave height, period, and direction was used to model the daily maximum significant wave height. Data on net primary production (NPP) were acquired from the Ocean Productivity online facility of Oregon State University (http://sites.science.oregonstate.edu/ocean.productivity/index.php). The Standard Vertical Generalized Production Model (VGPM) dataset was used. This is a “chlorophyll-based” model where NPP is a function of chlorophyll, available light, and photosynthetic efficiency. The NPP monthly aggregation was obtained and processed to make spatial datasets at the same spatial domain and pixels as the SST and nitrate concentration datasets. Monthly NPP data was also processed to produce seasonal and annual metrics of mean, maximum and minimum NPP (see Appendix S1: Table S1 for full list).

#### Seafloor terrain variables

We obtained spatial files of bathymetry data collected using multibeam sonar by the California Seafloor Mapping Project (CSMP; https://www.usgs.gov/centers/pcmsc/science/california-seafloor-mapping-program). To include information on substrate type (hard vs. soft) in the production of our models we also obtained binary files (rock vs sediment) available at 2 x 2 m resolution (4 m^2^). These datasets are publicly available for most of California with the exception of the northern Channel Islands (San Miguel, Santa Rosa, Santa Cruz and Anacapa). We obtained the fine scale bathymetry for the northern Channel Islands directly from NOAA. The mapped fine scale bathymetry of California is missing or limited in shallow waters (referred to as the ‘white zone’; 50 m to 500m offshore) where navigation hazards impede the access of seafloor mapping vessels and where multibeam sonar is generally infeasible due to underwater hazards and/or dense kelp canopies. To extend the bathymetry into the ‘white zone’ we interpolated the area between the mapped bathymetry and the shoreline using a natural neighbor algorithm in ArcGIS 10.2 (ArcGIS 10.2, Esri Industries, Redlands CA) (See Appendix S2 for interpolation details). To incorporate other characteristics of substrate in our models, we used the bathymetry aggregated to a 900 m^2^ grid (30 m x 30 m) to calculate slope (using the ‘raster’ package) and vector roughness measure (VRM; spatialEco) using R software (R Core Team 2022). Finally, we tested all bathymetry derivatives (depth, probability of rock, slope and VRM) for spatial autocorrelation and found that bathymetry was autocorrelated up to ~ 300 m, thus, all bathymetry files were aggregated to 300 m x 300 m resolution (90,000 m^2^) by calculating the mean using the raster package in R (R Core Team 2022).

#### Zoospore availability and density

We created spatial layers that estimated annual giant kelp zoospore availability at any given 900 m^2^ grid as a function of the maximum kelp canopy biomass each year and an empirical zoospore dispersal function. These layers were only estimated for giant kelp, since an equivalent dispersal function for bull kelp does not yet exist. The framework to create annual spatial files of maximum zoospore density per year required four steps: 1) Estimate giant kelp fecundity (cm^2^ of sorus area per m^2^ of kelp biomass) from Landsat-derived kelp biomass using a nonlinear relationship identified in previous work by Castorani et al. (2017); 2) estimate number of spores released per fertile unit (cm^2^ of sorus area per m^2^ of kelp biomass); 3) estimate the dispersal distances of spores for each pixel using a dispersal curve for giant kelp spores by Gaylord et al. (2006), and construct a map of zoospore dispersal densities; and 4) filtering the maps of zoospore density to areas with adequate substrate (rock). We repeated this procedure for every year we had Landsat-derived canopy biomass data, resulting in a series of maps of number of zoospores per 900 m^2^. We then aggregated these data to a 90,000 m^2^ grid (300 m x 300 m resolution) using the sum of zoospore abundance (See Appendix S3 for detailed methods). Zoospore density data was included in the giant kelp models as a lag effect variable, so that the zoospore density calculated for year 1, was used as a predictor for kelp density in year 2.

### Satellite-derived kelp abundance data

We used a remotely sensed time series of kelp canopy coverage (for bull kelp) and kelp canopy biomass (for giant kelp) to compare with the regional kelp abundance dynamics reconstructed from our kelp density models. Quarterly kelp canopy and biomass data for California from 1984 to 2022 were obtained from the Santa Barbara Coastal Long Term Ecological Research Program (SBC LTER) (Bell et al. 2023a). From this dataset we extracted the maximum area (bull kelp; North and Central regions) and biomass (giant kelp; Central and the two South regions) observed in each year, which usually occurred in quarter two (April-June) or quarter three (July-September) to obtain the maximum area or biomass for each cell per year. The data is available at 900 m^2^ resolution which we then aggregated into 90,000 m^2^ by summing the total maximum canopy area or biomass and then we calculated the mean canopy among all pixels per region, per year, to obtain a mean canopy estimate for each year, and region that we could compare to the projections obtained from our regional kelp density models.

### Spatio-temporal modeling of kelp density

#### Modelling framework

We modeled the density of each kelp species separately. The response variable for each model was the mean density of kelp estimated from *in situ* SCUBA surveys. Independent variables consisted of the environmental (e.g. temperature, nitrate, wave height, orbital velocity, NPP) and ecological variables (sea urchin and zoospore density). We first tested the models for each species in each of the regions to select the model with best predictive capacity. For bull kelp, we tested the models separately for the Northern and Central regions and then tested a model for both regions combined. For giant kelp we tested the models for the three regions containing giant kelp separately (Central, South-west and South-east) and then tested a model combining Central and South-west as previous studies indicate that these locations have similar species composition (Hamilton et al. 2010, Claisse et al. 2018, Carr et al. 2021). We used generalized additive mixed models (GAMs) (Wood 2006) to determine the best predictive relationship (i.e. deviance explained) between giant and bull kelp density and the independent ecological and environmental variables, and to identify the relative contribution of each independent variable in contributing to the model. Models were run in the ‘mgcv’ package in R (Wood 2011) with a tweedie distribution for the density of both species. The level of smoothing (number of basis ‘k’ in smoothing functions) for each predictor variable was restricted to 4 to avoid overfitting. We first assessed correlations between environmental variables using the training dataset and the corrplot package (Wei and Simko 2021), and removed all predictors with correlations higher than 0.7 to reduce intensifying effects of correlated variables. Due to the large number of predictor variables available, variable selection was first done by selecting a subset of metrics calculated from each type of variable (i.e. SST, nutrients, NPP, waves, substrate) selecting the ones with highest correlation with the dependent variable. Further variable selection was selection was completed conducting a full-subset approach implemented in the package FSSgam (Fisher 2022) which selected the best models (subset of predictors) based on performance and AIC. Prior to running the models, data were split into training and evaluation sets (70% and 30% of the dataset respectively). Final model selection was conducted with a 15-fold cross-validation approach and the best model selected based on R^2^ and deviance explained.

#### Spatial predictions of grazers

Density of purple sea urchins was the only biotic variable that was consistently selected in the models of kelp density for all regions and for both kelp species. Since spatial data on urchin densities were needed to recreate spatial maps of kelp density, we modeled the density of sea urchins using the same framework and variables as described above for kelp. Urchin models were tested separately in all four regions. Initial variable selection was done using the FSSgam package and model selection by conducting a 15-fold cross-validation. Historical maps of urchin densities were constructed by using the best models to predict in each region and for the years we had environmental data for (2004-2021) (Appendix S4: Table S1 and Figures S1-S18). Urchin density maps were created at the same 90,000 m^2^ resolution as the kelp maps.

### Reconstruction of historical kelp densities and their correlation with satellite-derived data

We used the predict function within the ‘mgcv’ package in R software (Wood 2011) and the historical spatial data of predictors selected in the best models for each region to create historical maps of kelp density for each species and each year for which we had environmental data (2004-2021) by projecting the density predictions over the study region (separately in each region). All predictor variables were converted to 90,000 m^2^ resolution (300 m × 300 m) to produce historical maps. This resolution was chosen based on the mean length of PISCO and Reef Check survey sites used for this study and based on spatial autocorrelation of the bathymetry data (depth and derivatives). Substrate and orbital velocity data that were at 30 m resolution, were aggregated to 300 m by calculating the mean, while other environmental variables which varied in spatial resolution from 1 to 5 km, were disaggregated using the ‘terra’ package in R (Hijmans 2022). The maps of kelp density produced with the best models were processed by limiting the maximum kelp density to the maximum kelp observed for each species from *in situ* surveys over the entire data series, and multiplied by the presence of rock in that area so that if the probability of rock in a given pixel was zero, this would result in zero rock in the produced maps.

## Results

### Predictors of spatial and temporal variation in bull kelp density

The density of bull kelp was best described by one model for both the North and Central regions combined (Table 1, Figure 2a-2g) with a deviance explained of 63%, R^2^ of 0.58 and the lowest AIC (AIC = 5098.27, Appendix 5: Table S1). The final model selected mean wave height, maximum orbital velocity, mean temperature during the upwelling season, mean density of purple sea urchins, depth, mean annual NPP, and minimum annual nitrate concentrations to best predict the density of bull kelp in California (Table 1, Figure 2a-2g).

**FIGURE 2.**
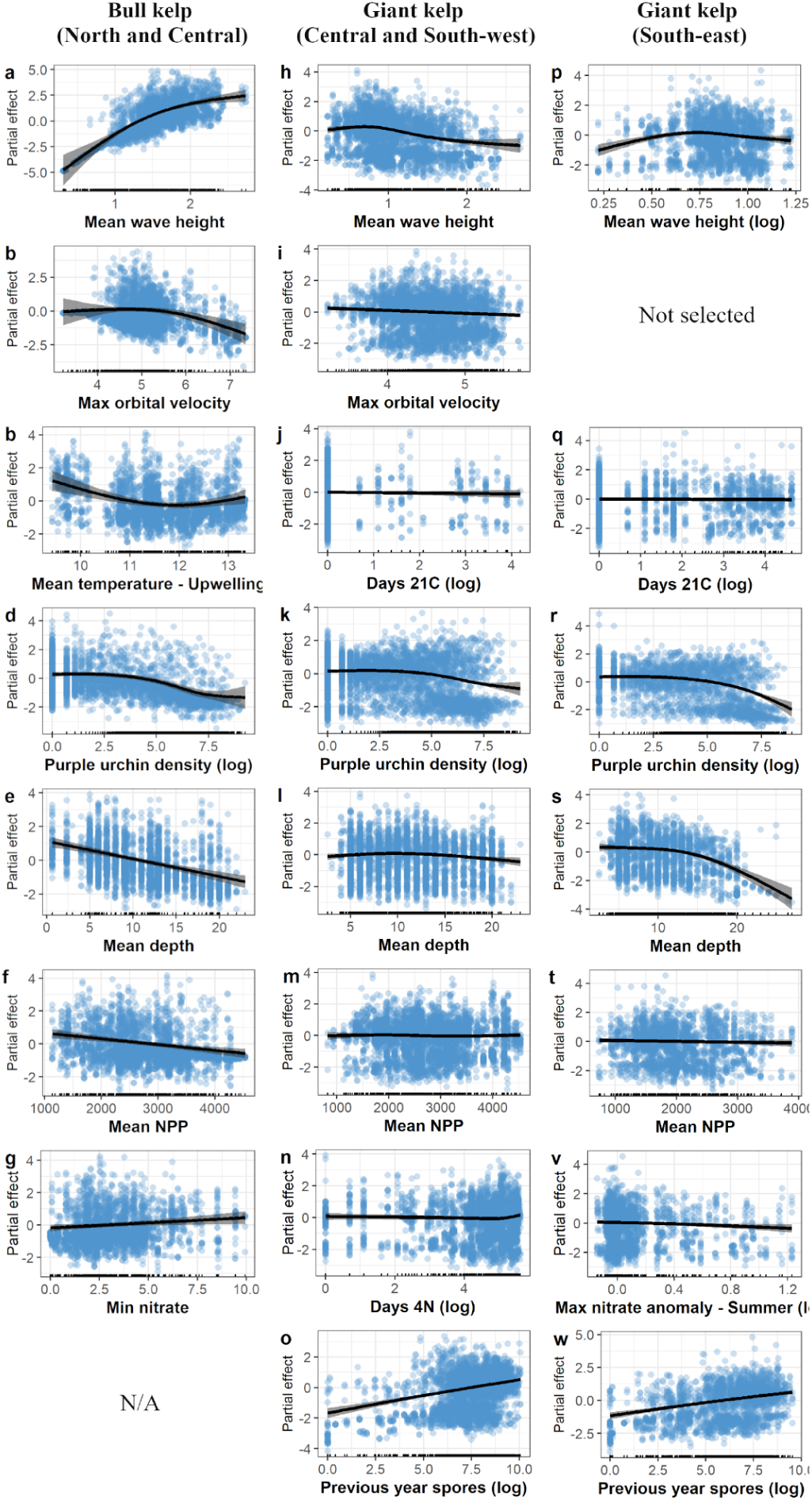
Drivers and trends of bull kelp density in north and central California (a-g), giant kelp in central/south-west region (h-o), and giant kelp in south-east region (p-w). Direction and magnitude of the predictors of best generalized additive models on kelp density in California. Shaded area indicates 95% confidence interval.

**TABLE 1.** Summary statistics of predictive performance of the best GAMs describing drivers of kelp density dynamics in California. Higuer r^2^ and deviance explained indicate improved model fit.

| Species | Region | Model | $r^2$ | Deviance explained |
| --- | --- | --- | --- | --- |
| *N. luetkeana* | Northern and Central | Minimum nitrate +<br>mean temperature upwelling +<br>mean wave height +<br>max orbital velocity (log) +<br>mean NPP +<br>depth +<br>density of purple sea urchins (log) | 0.58 | 63.30% |
| *M. pyrifera* | Central /South-west | Days 4N (log) +<br>maximum temperature anomaly +<br>mean wave height +<br>max orbital velocity (log) +<br>mean NPP +<br>depth +<br>density of purple sea urchins (log) +<br>Previous year spores (log) | 0.59 | 45.03% |
| *M. pyrifera* | South-east | Maximum nitrate summer anomaly (log) +<br>days 21C (log) +<br>mean wave height (log) +<br>mean NPP +<br>depth +<br>density of purple sea urchins (log) +<br>Previous year spores (log) | 0.62 | 51.44% |

The relationship between bull kelp density and mean wave height was positive asymptotic and was the strongest contributor to the model (Figure 2a). Wave heights > 1.2 m were positively correlated with bull kelp densities (relative importance = 1, *P*-value < 0.001, Figure 2a). The relationship with orbital velocity was non-linear and negative when maximum orbital velocity was higher than 5 m/s (relative importance = 1, *P*-value = 0.009, Figure 2b). Bull kelp density was inversely related to the mean temperature during the upwelling season and the decline was strongest across lower temperatures (relative importance = 1, *P*-value < 0.001, Figure 2c). The relationship with purple sea urchin density was negative, but only at urchin densities greater than 2.5 (log individuals/60 m^2^, ~ 0.18 individuals/m^2^) (relative importance = 1, *P*-value < 0.001, Figure 2d). Mean depth had a negative linear relationship with bull kelp density, with a negative effect at depths greater than 10 m (relative importance = 0.938, *P*-value < 0.001, Figure 2e). Mean NPP also displayed a negative linear relationship with bull kelp, with a negative effect at NPP higher than ~2,500 mg C m-2 d-1 (relative importance = 0.991, *P*-value < 0.001, Figure 2f). Finally, minimum nitrate had a positive linear relationship in the density of bull kelp, with the smallest magnitude of effect, with minimum nitrate values > 2.5 μmol L-1 showing a positive effect on bull kelp density (relative importance = 0.021, *P*-value = 0.07, Figure 2g).

### Predictors of spatial and temporal variation in giant kelp density

The density of giant kelp was best predicted by two models with different geographic regions combined. The first one, produced the best performance by combining the central and south-west regions into one model (from here referred to as central/south-west, deviance explained = 45%, R^2^ = 0.59; Appendix 5: Table S2). The second model produced the best results for the south-east region (deviance explained = 51%, R^2^ = 0.62). One single model for both central, south-west and south-east regions combined did not improve overall performance, and would have decreased the deviance explained and R^2^, especially for the south-east coast (Appendix 5, Table S2).

The best models identified for giant kelp in the central/south-west and south-east regions shared very similar predictors to each other with a few exceptions (Figure 2h-2w). Mean wave height emerged as a more important predictor of giant kelp in the central/south-west region (relative importance = 1, *P*-value < 0.001) than in the south-east coast (relative importance = 0.561, *P*-value < 0.07) (Figure 2h and 2p). In the central/south-west region, there was a negative effect of wave heights greater than 1 m (Figure 2h), while in the south-east coast mean wave heights did not reach that magnitude and a small negative effect was displayed when wave height was greater than approximately 0.65 m (Figure 2p). Maximum orbital velocity was important for giant kelp density in the central/south-west region, showing a negative effect at values greater than 4 m/s (relative importance = 0.689, *P*-value < 0.01, Figure 2i). The addition of maximum orbital velocity to the south-east coast model did not improve the AIC and it was not included in the final model for that region. Days above 21°C (log) for temperature was selected in both models, however its importance was low compared to other predictors (relative importance of 0.081 and 0.225 and *P*-value of 0.06 and 0.18 for the central/south-west region and south-east region models respectively), with a negative effect above ~ 2.5 log days above 21°C (Figure 2j and 2q).

Density of purple sea urchins showed a non-linear relationship and one of the largest effects on giant kelp density in both regions, with a negative relationship at urchin densities higher than ~5 (log urchins/60 m^2^, ~ 2.5 individuals/m^2^) (relative importance = 1, *P*-value < 0.001, in both regional models Figure 2k and 2r). Depth was also a significant predictor of giant kelp in both models, showing a negative relationship at depths greater than 15 m however, the relationship was stronger (greater slope) in the south-east (relative importance = 1, *P*-value < 0.001) than in the central/south-west region (relative importance = 0.996, *P*-value < 0.001) (Figure 2l and 2s). Mean NPP was another predictor selected in both regional models, however, its importance in the central/south-west region (relative importance = 0.026, *P*-value = 0.71) was much lower than in the south-east coast (relative importance = 0.654, *P*-value = 0.04) (Figure 2m and 2t), and displayed a different relationship with giant kelp in both regions. In the central/south west region, mean NPP showed a slight positive effect at NPP higher than 4000 mg C m^−2^ d^−1^ (Figure 2m) while in the south-east coast, it showed a negative linear relationship with kelp density with negative effects at NPP higher than 2500 mg C m^−2^ d^−1^ (Figure 2t). Different variables were selected to describe the relationship between nutrients and giant kelp density for the central/south-west region compared to the south-east region (Figure 2n and 2v). The central/south-west model selected for days above 4 μmol L^−1^ as the best nitrate predictor (relative importance = 0.995, *P*-value < 0.001) and a non-linear relationship with kelp density, with a positive effect when concentration of nitrate where above 4 μmol L^−1^ for more than 5 log days (Figure 2n). On the other hand, the best model for the south-east region selected for the maximum anomaly of nitrate concentration during the summer season, showing a negative linear relationship and a negative effect after anomalies higher than log 0.25 μmol L^−1^ (relative importance = 0.722, *P*-value < 0.001, Figure 2v). Finally, abundance of zoospores in the previous year also had a large effect on giant kelp density in both regions, displaying a positive linear relationship with kelp density at around 7.5 log spores/m^2^ (relative importance = 1, *P*-value < 0.001, in both regional models Figure 2o and 2w).

### Reconstructed spatial dynamics of kelp density

Using the selected variables in each of the three best models described above, the density of kelp was reconstructed for each year from 2004 to 2021. The effect of the NE Pacific MHW is well captured by the models, which show the collapse in bull kelp populations from 2013 to 2014 (Appendix S5: Figure S3). After 2014, the reconstructed maps show the depletion of bull kelp populations across the north region (Appendix S5: Figures S4-S5). The reconstructed maps also show an increase in kelp densities in 2018 and from 2020 to 2021 (Appendix S5: Figures S4-S5), where a large and widespread recovery of bull kelp populations was predicted but was not observed by divers or was detected by remote sensing, indicating a mismatch between the drivers of kelp density before and after the NE Pacific MHW. Yet, the estimated mean kelp densities before the NE Pacific MHW (2004-2013) are visibly higher for the entire north coast region than after the NE Pacific MHW (Figure 3).

**FIGURE 3.**
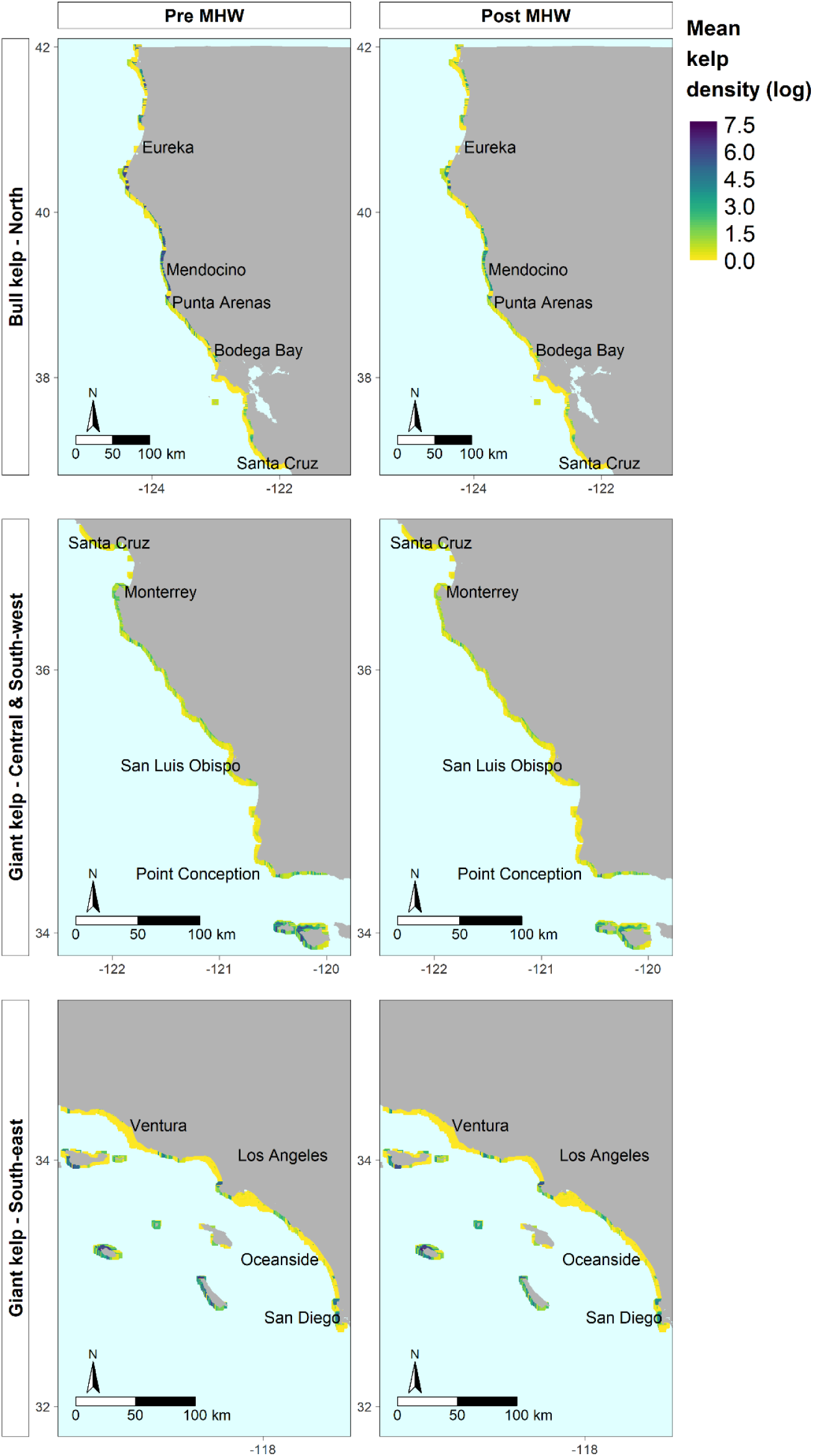
Model predicted kelp density distribution between 0 and 30 m of depth prior to (“Pre-MHW”, 2004 to 2013) and after (“Post-MHW”, 2014 to 2021) the NE Pacific MHW. Cells are color coded corresponding to the mean predicted kelp density (log scale) across the years of each period obtained from the best model for bull kelp and the two models for giant kelp (Appendix 5: Figures S1 and S15). North region is depicting bull kelp density only since bull kelp is dominant in this region (top row), central/south-west region is depicting density of giant kelp since kelp beds in this region are predominantly dominated by this species (middle row), and the south-east region is depicting only predicted mean densities of giant kelp (bottom row).

The reconstructed maps of predicted giant kelp density in the central/south-west regions show a general decline in kelp density from 2013 to 2014 (Figure 3), particularly at the mainland sites, while the decline in kelp density at the islands in that region becomes apparent only after 2016 with no significant recovery up to 2021 (Appendix 5: Figures S9-S10). A general decline in giant kelp densities was projected after the NE Pacific MHW compared to mean kelp densities before the NE Pacific MHW (2004-2013) (Figure 3).

The model for the south-east region showed a patchy distribution with fluctuations in kelp densities between 2004 and 2015 (Appendix 5: Figures S11-S13) with no significant decline in kelp densities observed in the region from 2013 to 2014 as was observed for bull kelp in the north coast and giant kelp in the central/south-west coast (Figure 3).

A visible decline in kelp densities over the entire region was observed in 2016 potentially in response to the NE Pacific MHW, however, in subsequent years an increase in kelp was observed with similar fluctuations as observed previous to the NE Pacific MHW (Appendix 5: Figures S13-S15). Mean kelp densities were slightly higher before than after the NE Pacific MHW, with more evident declines observed north of San Clemente, in San Clemente Island and in the San Diego region (Figure 3).

### Predicted kelp density vs satellite-derived kelp canopy abundance

Correlations between the mean annual kelp densities predicted from the best models with the mean annual kelp canopy abundance (area for the bull kelp model or biomass for the giant kelp models) derived from Landsat imagery were all positive and significant for each species and region indicating that the interannual dynamics of kelp canopy were generally well captured by the models (Figure 4). Mean annual bull kelp density in the north coast predicted by the models generally followed the fluctuations of kelp area observed from Landsat imagery (r_(16)_ = 0.66, p = 0.0026) (Figure 4a-4c). A large discrepancy between predicted bull kelp density and observed kelp area in the north coast is evident in 2021 (Appendix 5: Figures S5), when predicted kelp density is higher than the observed kelp area (Figure 4c) and in fact, when analyzed separately, the high and significant correlation between predicted kelp density and observed kelp canopy from Landsat pre-MHW (r_(8)_ = 0.82, p = 0.0036, Figure 5a) becomes low and not significant after the NE Pacific MHW (r_(6)_ = 0.23, p = 0.58, Figure 5b).

**FIGURE 4.**
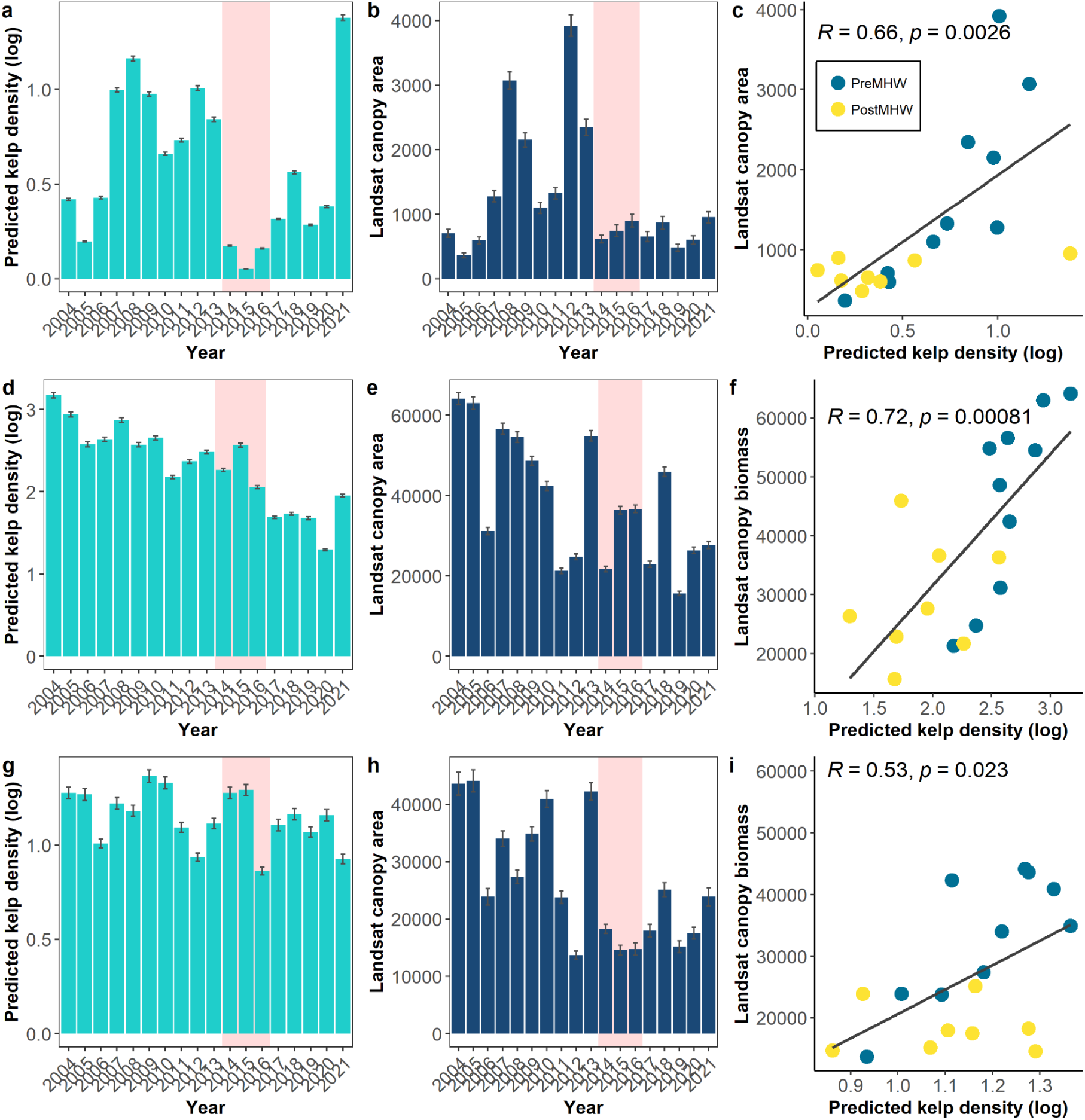
Mean annual predicted kelp density (log) (a,d,g), mean annual kelp canopy abundance (area or biomass) detected from Landsat (b, e, h) and correlation between these two (c, f, i). For bull kelp in the north coast (kelp canopy area in m^2^ per 90,000 m^2^, top row), for giant kelp in the central/south-west region (kelp canopy biomass in kg per 90,000 m^2^, middle row) and giant kelp in the south-east region (kelp canopy biomass in kg per 90,000 m^2^, bottom row). Bars and points are the mean annual kelp density, area or biomass over 90,000 m^2^ pixel per year for each region (± standard error, for bars).

**FIGURE 5.**
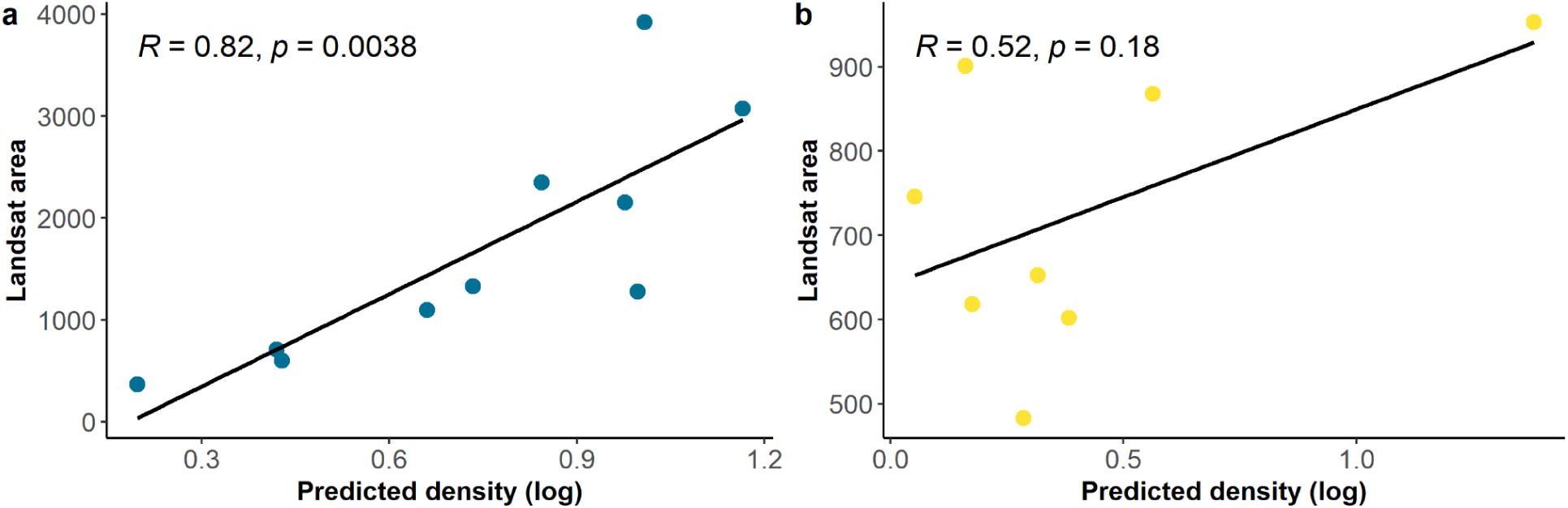
Correlation between mean annual predicted kelp density and mean annual kelp canopy area (in m^2^ per 90,000 m^2^) detected from Landsat for bull kelp in the north coast before (a) and after (b) the NE Pacific MHW. Bars and points are the mean annual kelp density, area or biomass for each region per 90,000 m^2^ pixel per year (± standard error, for bars).

Mean annual predicted giant kelp densities in the central/south-west region showed a general trend of decline that began before the NE Pacific MHW and which is also observed in the biomass derived from Landsat (Figure 4d-4f). The correlation between predicted kelp densities and kelp area from Landsat was strongest for the giant kelp model in the central/south-west region (r_(16)_ = 0.72, p = 0.0081) (Figure 4f).

Finally, mean annual predicted giant kelp densities in the south-east region, showed fluctuations in kelp density from year to year, with a slight trend of decline that was not associated with the NE Pacific MHW conditions, as the years 2014 and 2015 were predicted to have had high kelp densities (Figure 4g-4i).

This is a discrepancy with the kelp biomass observed from Landsat where the years 2014 and 2015 showed some of the lowest biomass observed between 2004 and 2021 (Figure 4g-4h). The mean annual kelp biomass in the south coast derived from Landsat shows large fluctuations from year to year from 2004 to 2013 and lower fluctuations from 2014 to 2021 (Figure 4h). Lower fluctuations in mean annual kelp densities were predicted from the model for the entire time series, which resulted in this region having the weakest correlation between predicted kelp density and kelp biomass (r_(16)_ = 0.53, p = 0.023) (Figure 4i).

## Discussion

Kelp forests are dynamic ecosystems that naturally experience great variability across space and time, yet, globally and in California, they have become increasingly threatened by multiple stressors that are exacerbated by climate change (Krumhansl et al. 2016, Wernberg et al. 2016, Arafeh-Dalmau et al. 2021a). Indeed, many regions have lost all or much of their kelp forests and are struggling with management and restoration decisions to stem further losses and/or rebuild populations (Rogers-Bennett and Catton 2019, Butler et al. 2020, Hynes et al. 2021, Miller et al. 2023). The first step to developing management actions to ensure the persistence of kelp forest ecosystem functioning and services under a changing climate is to understand the key environmental and ecological factors that shape both temporal and spatial patterns of abundance. Kelps are unlike many terrestrial forest systems, being highly variable in time and space. Here we have identified the key environmental and biological drivers of two dominant surface canopy-forming kelp species (bull kelp and giant kelp) across the 1,350 km of coast of California. The results from our models are robust as the drivers identified in this approach were useful in reconstructing regional patterns of historical kelp density from remotely-sensed canopy cover. However, we also identified important mismatches in the predicted dynamics resulting from the two data sources, that arise from ecological processes (e.g. grazing by sea urchins) which are not captured by remote sensing yet are critical to kelp population dynamics. The results from this study highlight the importance of long-term *in situ* monitoring surveys of complex ecosystems such as kelp forests (Magurran et al. 2010, Hughes et al. 2017), as well as the role for remote sensing to fully understand population dynamics (Cavanaugh et al. 2021) which are key for effective management of these species and the ecosystems they sustain.

### Drivers of bull kelp distribution and dynamics

The best model we identified for bull kelp covered both the north and central coasts combined, indicating that the drivers of bull kelp density were the same for the entire extent of its distribution in the state of California. Increasing wave height had the largest effect on bull kelp density, as expected for a species that dominates exposed coastlines such as found in Northern California (Springer et al. 2010, Carr and Reed 2016). In central California, bull kelp commonly occurs in mixed beds with giant kelp where it thrives in the shallower areas that experience breaking waves as evidenced by its higher density in areas with increased wave heights compared to giant kelp (Springer et al. 2010). Higher water motion, brought on by waves may aid in the uptake of nutrients and carbon (Koehl and Alberte 1988, Hurd 2000), increase irradiance by pushing the fronds in different directions (Koehl and Alberte 1988), and enhance blade production (Breitkreutz et al. 2022). However, an upper threshold of water motion for bull kelp appeared to be reached at maximum orbital velocities higher than ~ 5 m/s, beyond which density was reduced, likely the result of the ripping of kelp stipes from increased drag in high flow speeds (Johnson and Koehl 1994).

Bull kelp inhabits areas of significant coastal upwelling, which delivers cold and nutrient-rich water to coastal areas (Springer et al. 2010). The optimal growing temperature for adult and early life stages of bull kelp in Canada has shown to be ~ 11.9°C with an upper thermal limit between 18°C and 20°C (Supratya et al. 2020). For California we found, similarly, that bull kelp density increased at mean upwelling season temperatures lower than 12°C. Sea surface temperature during the upwelling season is negatively associated with nutrient availability, indicating that nutrient limitation during this time is usually what drives the interannual dynamics of bull kelp (McPherson et al. 2021, García-Reyes et al. 2022). Further evidence for the importance of nutrient limitation to bull kelp dynamics was the positive relationship between bull kelp density and minimum annual nitrate which indicated that the higher the nitrate concentration from the physiological thresholds of bull kelp, the better it does in general.

Increasing SST and lower nutrient levels combined with an increase in herbivory levels have been shown to be important drivers in the collapse of bull kelp populations after the NE Pacific MHW (McPherson et al. 2021). Densities of purple sea urchins spiked after the NE Pacific MHW, potentially due to the combination of a lack of top-down control of sunflower sea stars on herbivorous sea urchins (McPherson et al. 2021) and anomalously high settlement of sea urchins around the Fort Bragg region between 2013 and 2015 resulting in high recruitment (Okamoto et al. 2020). Densities of bull kelp did not show a response at low urchin densities, but declined dramatically at urchin densities greater than ~ 4.5 log urchins per 60 m^2^ transect (equivalent to about 1.5 urchins/m^2^). This parallels previous findings that show urchin densities of about 0.1 to 1.7 urchins/m^2^ before 2013, which increased up to 60 fold during the NE Pacific MHW and persisted through the collapse of bull kelp coverage in northern California (Rogers-Bennett and Catton 2019). Importantly, after the NE Pacific MHW, our model predicted a recovery of bull kelp in 2018 and 2021, when our models predicted a decline in urchin populations and favorable abiotic conditions for kelp had returned. However, this predicted recovery of bull kelp abundances was not evident in satellite imagery or *in situ* diver surveys in the region (Cavanaugh et al. 2023). This result may be evidence of hysteresis in the system driven by sea urchin overgrazing (Filbee-Dexter and Scheibling 2014).

Net primary productivity was another factor associated with bull kelp density dynamics. High values of NPP may indicate that kelp is competing for nutrients and light with phytoplankton in the water column (Kavanaugh et al. 2009), explaining why NPP values higher than 2,500 mg C m^−2^ d^−1^ resulted in a decline in bull kelp. Evidence of competition for nutrients at large spatial scales has been shown in other kelps, especially during high-nutrient conditions driven by interannual climatic oscillations (Dayton et al. 1999). Densities of bull kelp declined linearly with depth likely due to light attenuation (Goldberg and Kendrick 2004). In bull kelp, light is the most important factor in the development of gametophytes and sporophytes, and allows sporophytes to reach sexual maturity (Vadas 1972). We also found that grazers like sea urchins were less abundant in shallower areas, potentially due to higher wave energy in shallow areas (Duggins et al. 2001). The combination of higher light levels and lower grazer density potentially creates an optimum environment in the shallows for bull kelp to thrive.

### Drivers of giant kelp distribution and dynamics

Unlike bull kelp, two models were required to best describe the distribution of giant kelp in California and were generally consistent with different oceanographic conditions between the central and southern coasts. These domains are not defined by the typical biogeographic limit of Point Conception (Blanchette et al. 2008, Hamilton et al. 2010, Claisse et al. 2018), but rather by an approximate midpoint in Santa Barbara Channel, which has been shown to be the limit between a system with more nutrient availability and lower sea surface temperatures driven by upwelling conditions to the west of the Santa Barbara Channel, including Santa Rosa and San Miguel islands (Broitman et al. 2005, Hamilton et al. 2010, Gosnell et al. 2014, Claisse et al. 2018).

Large waves are one of the most important causes of giant kelp mortality in California (Dayton et al. 1984, Reed and Foster 1984) and wave height was an important driver of kelp density in our models, but the effect was different in the two regions. In the central/southwest coast, which naturally experiences larger waves than the south-east coast (Reed et al. 2011), kelp density was resilient to mean annual wave heights up to approximately 1 m, while in the south coast, kelp density was the highest at lower wave heights of approximately 0.75 m. This regional difference in wave disturbance has been shown to determine giant kelp net primary productivity in California (Reed et al. 2011, Castorani et al. 2022). Stronger and more frequent wave events in the central/southwest coast may explain why orbital velocity was a significant factor negatively correlated with kelp density in that region but not in the calmer, south coast. In shallow waters, the horizontal motion of a wave may be up to 5 times greater than the wave height, creating drag forces which, when strong enough, break stipes or remove holdfasts (Seymour et al. 1989). Previous studies in central California have found wave orbital velocity to be the most frequent disturbance responsible for tearing out plants (Graham 1997) and giant kelp in the region can occur across a range of wave orbital velocities but is most abundant in a moderate wave environment (~ 0.86 m/s) (Young et al. 2016).

The nutrient environment varies widely along the distribution of giant kelp in California, which covers approximately 10 degrees of latitude. Central California generally has a more stable and consistent nutrient supply through upwelling due to the exposed coastline and narrow continental shelf than the south coast (Huyer 1983, Zimmerman and Kremer 1984). Here, we add empirical evidence suggesting that kelp populations are adapted to local nutrient conditions, a process which has been observed in a number of marine species (Sanford and Kelly 2011, Howells et al. 2011, Bennett et al. 2015). Interestingly, in central/south-west California, kelp density was positively related to the number of days above 4 μmol L^−1^ while in the south coast, the maximum nitrate anomaly during the summer season was the more important predictor, and was negatively associated with kelp growth. Positive growth of giant kelp in southern populations has been shown to occur at extremely low nitrate concentrations (less than 1 μmol L^−1^), while in central California a positive effect on giant kelp biomass is only visible past the 4 μmol L^−1^ (Kopczak et al. 1991). During the summer, declines in giant kelp are usually related to the reduction in nutrient availability (Jackson 1977, Gerard 1982, North and Zimmerman 1984), with the principal nutrient source during summer and fall through internal wave propagation (Zimmerman and Kremer 1984). The negative relationship between maximum nitrate anomaly in the summer months and kelp density that we found in the south coast is hence, unexpected. This may be explained by low rates of frond production during summer, which may not be able to keep up with natural frond loss (Zimmerman and Robertson 1985). Previous work has found that frond dynamics are better explained by intrinsic biological processes, such as frond age, rather than external environmental conditions (Rodriguez et al. 2013), indicating that kelp senescence, which in southern California occurs in summer, can overpower external environmental conditions (Bell and Siegel 2021).

As with nutrient availability, there is also evidence of adaptation to thermal stress in the microscopic reproductive stages of giant kelp (Ladah and Zertuche-González 2007). A strong inverse relationship between SST and nitrate availability is known to exist in California (Zimmerman and Robertson 1985) with warm sea temperatures typically associated with reduced upwelling and nutrient limited conditions. Although the number of days above 21° C was selected as an important driver for kelp density in central and southern California, we found it had a small effect in both regions. Previous studies have also found SST metrics not to be significantly related to the resilience of giant kelp to warming events (Cavanaugh et al. 2019), indicating that nutrients and other local processes are more determinant at driving kelp populations than temperature, both in central and southern California. We also found NPP to be an important factor explaining kelp density, with a small negative effect on kelp density overall. Net primary productivity (NPP) can indicate the levels of plankton abundance and extensive blooms of phytoplankton can reduce light intensity (Kavanaugh et al. 2009) especially during upwelling periods (Strub and Powell 1987) when kelp is uptaking nutrients and growing. This could result in competition for both light and nutrients between phytoplankton and the kelp recruits, explaining the negative effect of increasing NPP. Competition for nutrients between giant kelp and other species has been has been shown to be more noticeable in shallow depths and during large-scale low frequency events that drive nutrient-rich conditions (such as La Niña), as these drive surface nutrient availability and have long-term influence on the surface canopy of giant kelp (Dayton et al. 1999).

Both the density and foraging behavior of grazers have been shown to be an important factor determining the distribution and biomass density of kelp beds (Johnson et al. 2011, Filbee-Dexter and Scheibling 2014, Bell et al. 2015, Young et al. 2023). Similar to studies in other regions (Filbee-Dexter and Scheibling 2014, Balemi and Shears 2023, Ling and Keane 2024) we found that giant kelp declined in both central/south-west and south-east regions of California when densities of sea urchins exceeded approximately 2.5 urchins/m^2^ (5 log urchins per 60 m^2^ transect). However, a previous study of giant kelp abundance in the central coast of California identified some sites with a positive relationship between urchins and kelp (Bell et al. 2015).

Indeed, under some circumstances, high density of giant kelp can co-exist with high densities of grazers (Karatayev et al. 2021, Randell et al. 2022, Rennick et al. 2022). However, this was not the case for giant kelp in central and northern California following the NE Pacific MHW, where urchins increased dramatically to previously undocumented levels (Rogers-Bennett and Catton 2019). This increase in grazing combined with increased physiological stress in the kelp caused by high SST and low nutrient levels may explain the post-MHW decline in giant kelp, which although not as dramatic as for bull kelp in the north coast, was more evident in the central/south-west, than the south-east region.

Demographic connectivity is key to the persistence of kelp metapopulations, however, it is highly variable due to spore supply and dispersal (Hanski 1998, Castorani et al. 2015). Previous findings in southern California found that temporal variation in fecundity (i.e. spore supply) had a larger effect on the persistence and recovery of giant kelp beds than variation in the physical transport of spores (Castorani et al. 2017). Here, we used a spore supply metric that accounted for annual variation based on the maximum kelp biomass from the previous year (a proxy for spore production) and found that it was one of the most important drivers of giant kelp density in both regions. While this might also be the case for bull kelp, we currently do not have a relationship between adult biomass and spore production, or dispersal information for that species.

Depth is known to be one of the most important factors affecting kelp growth and recruitment as it is correlated with light availability (Gerard 1984). Our models found that giant kelp densities declined below depths of 15 m in both regions, with a stronger negative effect in the south-east coast. In addition to depth, light attenuation beneath the kelp canopy can be reduced up to 99% at 20 m of depth (Dean 1985). Such low irradiances may reduce kelp density by limiting recruitment, as microscopic stages of kelp have high light requirements and are vulnerable to intraspecific competition through shading (Stewart et al. 2009).

### Comparison with Landsat observations

Our comparison of the regional model predictions of *in situ* kelp density with canopy cover estimated from Landsat indicated that dynamics from the two sources match quite well for both kelp species. Thus, the kelp dynamics model performed well in predicting the dynamics of forest density of both canopy-forming species and provided confidence in spatially projecting forest stability of both kelps throughout their California ranges. But the comparison also revealed a critically important limitation of the model. The bull kelp density dynamics model predicted the recovery of forests in the years following the NE Pacific MHW whereas Landsat did not detect a recovery. This mismatch is likely due to a hysteresis effect that the modeling approach did not capture, where abundances of kelp did not track abundances of urchins after the MHW as they did before the MHW. Persistent overgrazing of macroalgae by urchins has been known to shift algal ecosystems into an alternative stable state from which it is challenging to recover because although high urchin densities are required to overgraze kelp, much lower densities are required to maintain the deforested, urchin barren state (Filbee-Dexter and Scheibling 2014, Ling et al. 2019). One study found that the biomass of urchins required to tip a kelp forest into a barren state is one order of magnitude higher than that required to maintain the urchin barrens state (Ling et al. 2015). As such, the ecosystem shift into this alternative state can be difficult to reverse, even when abiotic conditions are conducive for forest recovery, and such change in ecological relationships is not adequately captured by the model. In the case of northern California, further analyses of urchin densities from *in situ* surveys are required to confirm such change in the kelp-urchin relationship, once more reinforcing the necessity to couple *in situ* and remote sensing monitoring data to inform the interpretation of ecological processes in modeling results.

### Implication for understanding future disturbances and management of kelp forests

Identifying the biotic and abiotic factors that drive spatial and temporal variation in abundance and condition of foundation species is key for successful management of biodiversity under a changing climate. Our kelp density models quantified the functional relationships between various environmental and ecological drivers and kelp. These models explained and predicted spatial and temporal variation in the two kelp species quite well across markedly different geographic regions. However, the bull kelp model failed to predict the persistence of a deforested state as abiotic conditions were favorable for kelp. In this case, ecological interactions such as overgrazing by sea urchins, superseded the ability of kelps to naturally recover. This result reinforces the importance of coupling multiple sources of data, including *in situ* and remotely sensed, to predict future dynamics. It also highlights the need to develop models that incorporate shifts in ecological interactions, such as that between urchins and kelp (e.g., (Karatayev et al. 2021, Arroyo-Esquivel et al. 2023). While we expect that the environmental drivers identified in this study will vary in strength and spatial distribution with future climate change, we also assume that the functional relationships between kelp growth or loss and those drivers will remain constant in the near future (years to decades). If the assumption holds, then the models created here, when coupled with knowledge or predictions of urchin densities and projections of environmental variables, should accurately reproduce kelp dynamics across the state for the upcoming years, and can be used to predict the dynamics of kelp into the future. However, if the fundamental nature of the relationships changes with climate change, then those relationships will need to be explored with experiments, and new models will need to be developed. How might the relationship between the drivers of kelp dynamics as we know them now, change in the future? Nutrients and temperature are important to primary producers, all of which possess specific photosynthetic temperature response curves that define an optimal temperature for photosynthesis and critical temperature thresholds. These are known for California kelps (Zimmerman and Kremer 1984, Bell et al. 2015, Cavanaugh et al. 2019) but as sea temperature increases local adaptation could change the response curves and the thresholds. Changes in the spatial distribution of various ecotypes of kelp through movement (passive or assisted) could also change these relationships. Similarly, interventions to assist the recovery of kelp forests, may accelerate local adaptation and alter the response to environmental variables found here, so as to contribute towards more resilient or resistant populations that can withstand contemporary and future environmental conditions.

As we and others have demonstrated (Bell et al. 2015, Cavanaugh et al. 2019, McPherson et al. 2021), kelp forests are dynamic systems that can respond rapidly to fluctuating environmental drivers. Much of our understanding of the dynamic relationships between these oscillations and kelp dynamics is due to the availability of multidecadal time series of kelp canopy from remotely sensed imagery (Bell et al. 2023b) and increasingly from i*n situ* long-term monitoring of kelp forests. The *in situ* diver surveys from long-term monitoring programs in California provided key data that enabled the model construction of kelp and urchin densities, and identification of ecological processes (i.e. grazing) that contribute to their temporal and spatial patterns of abundance in the study region. The co-located and simultaneous sampling of kelp and urchin densities was key to capture the covariation of urchin and kelp densities and incorporated that covariance into the species distribution models (GAMs). These models revealed the important inverse relationships between purple urchin density with either kelp species. Furthermore, our analyses allowed for an independent assessment of the drivers of kelp dynamics, and the important drivers were similar to those identified using Landsat data from previous studies (Bell et al. 2015), providing added confidence in both methods.

## Conclusion

Understanding how biotic and abiotic factors contribute to the temporal variation in the abundance of foundation species is a major focus of ecology and biogeography and is key to understanding biodiversity in a changing climate. Our results demonstrate the benefits of combining long-term *in situ* monitoring data that provides information about species interactions (not yet obtainable through remote sensing) with remote sensing datasets to interpret population modelling results. Using both *in situ* and remote sensing data to understand the dynamics of surface canopy kelps allowed for the recognition of a tipping point in the system and validate the robustness of the model outputs. The maps produced from these robust models provide valuable information for managers and stakeholders about the locations that are more likely to support healthy kelp ecosystems and the functional relationships identified, form a basis for future focused on the future spatio-temporal dynamics of kelp forests under a changing climate (Giraldo-Ospina et al. 2023a).

## Acknowledgements

We thank the numerous individuals who contributed to data collection as part of the long-term kelp forest monitoring programs of Partnership for Interdisciplinary Studies of Coastal Oceans (PISCO) and Reef Check California. We also thank D. Malone, A. Parsons-Field for assistance with data curation and J. Freiwald for assistance with Reef Check data acquisition. This research was funded by California Sea Grant (R/HCEOPC-18) as part of the California state-wide Kelp Recovery Research Program in collaboration with the California Ocean Protection Council. We also wish to thank M. Yeager, the Caselle lab members and Passionate Women in Science (PWIS) team for support, advice and fruitful discussions.

## Author Contributions

JC, MC and TB conceived the project ideas and obtained funding. JC, MC and AGO contributed with biological data collection. TB contributed with abiotic data collection and Landsat data collection. JC, TB, AGO and MC conceived methodology. AGO led the conceptualization and writing of the manuscript and performed all analyses. JC, MC and TB contributed to the conceptual design of the manuscript and manuscript writing. All authors gave final approval for publication.

## References

Alongi, D. M. 2015. The Impact of Climate Change on Mangrove Forests. Current Climate Change Reports 1:30–39.

Arafeh-Dalmau, N., I. Brito-Morales, D. S. Schoeman, H. P. Possingham, C. J. Klein, and A. J. Richardson. 2021a. Incorporating climate velocity into the design of climate-smart networks of marine protected areas. Methods in ecology and evolution / British Ecological Society 12:1969–1983.

Arafeh-Dalmau, N., K. C. Cavanaugh, H. P. Possingham, A. Munguia-Vega, G. Montaño-Moctezuma, T. W. Bell, K. Cavanaugh, and F. Micheli. 2021b. Southward decrease in the protection of persistent giant kelp forests in the northeast Pacific. Communications Earth & Environment 2:1–7.

Arafeh-dalmau, N., G. Montaño-moctezuma, J. A. Martínez, and D. A. Smale. 2019. Extreme marine heatwaves alter kelp forest community near its equatorward distribution limit. Frontiers in Marine Science 6:499.

Arroyo-Esquivel, J., M. L. Baskett, M. McPherson, and A. Hastings. 2023. How far to build it before they come? Analyzing the use of the Field of Dreams hypothesis in bull kelp restoration. Ecological applications: a publication of the Ecological Society of America 33:e2850.

Assis, J., E. Fragkopoulou, D. Frade, J. Neiva, A. Oliveira, D. Abecasis, S. Faugeron, and E. A. Serrão. 2020. A fine-tuned global distribution dataset of marine forests. Scientific data 7:119.

Balemi, C. A., and N. T. Shears. 2023. Emergence of the subtropical sea urchin Centrostephanus rodgersii as a threat to kelp forest ecosystems in northern New Zealand. Frontiers in Marine Science 10.

Beas-Luna, R., F. Micheli, C. B. Woodson, M. Carr, D. Malone, J. Torre, C. Boch, J. E. Caselle, M. Edwards, J. Freiwald, S. L. Hamilton, A. Hernandez, B. Konar, K. J. Kroeker, J. Lorda, G. Montaño-Moctezuma, and G. Torres-Moye. 2020. Geographic variation in responses of kelp forest communities of the California Current to recent climatic changes. Global change biology 26:6457–6473.

Beck, M. W., R. D. Brumbaugh, L. Airoldi, A. Carranza, L. D. Coen, C. Crawford, O. Defeo, G. J. Edgar, B. Hancock, M. C. Kay, H. S. Lenihan, M. W. Luckenbach, C. L. Toropova, G. Zhang, and X. Guo. 2011. Oyster Reefs at Risk and Recommendations for Conservation, Restoration, and Management. Bioscience 61:107–116.

Bell, T., K. Cavanaugh, and D. Siegel. 2023a, January 1. SBC LTER: Time series of quarterly NetCDF files of kelp biomass in the canopy from Landsat 5, 7 and 8, since 1984 (ongoing).

Bell, T. W., K. C. Cavanaugh, D. C. Reed, and D. A. Siegel. 2015. Geographical variability in the controls of giant kelp biomass dynamics. Journal of biogeography 42:2010–2021.

Bell, T. W., K. C. Cavanaugh, V. R. Saccomanno, K. C. Cavanaugh, H. F. Houskeeper, N. Eddy, F. Schuetzenmeister, N. Rindlaub, and M. Gleason. 2023b. Kelpwatch: A new visualization and analysis tool to explore kelp canopy dynamics reveals variable response to and recovery from marine heatwaves. PloS one 18:e0271477.

Bell, T. W., and D. A. Siegel. 2021. Nutrient availability and senescence spatially structure the dynamics of a foundation species. Proceedings of the National Academy of Sciences of the United States of America 119:e2105135118.

Bennett, S., T. Wernberg, B. Arackal Joy, T. de Bettignies, and A. H. Campbell. 2015. Central and rear-edge populations can be equally vulnerable to warming. Nature communications 6:10280.

Blanchette, C. A., C. Melissa Miner, P. T. Raimondi, D. Lohse, K. E. K. Heady, and B. R. Broitman. 2008. Biogeographical patterns of rocky intertidal communities along the Pacific coast of North America. Journal of biogeography 35:1593–1607.

Bolton, J. J. 2010. The biogeography of kelps (Laminariales, Phaeophyceae): a global analysis with new insights from recent advances in molecular phylogenetics. Helgoland marine research 64:263–279.

Breitkreutz, A., L. J. M. Coleman, and P. T. Martone. 2022. Less Than the Sum of Its Parts: Blade Clustering Reduces Drag in the Bull Kelp, Nereocystis luetkeana (Phaeophyceae). Journal of phycology 58:603–611.

Briggs, J. C. 1974. Marine Zoogeography. McGRaw-Hill Book Company, New York.

Broitman, B. R., C. A. Blanchette, and S. D. Gaines. 2005. Recruitment of intertidal invertebrates and oceanographic variability at Santa Cruz Island, California. Limnology and oceanography 50:1473–1479.

Butler, C. L., V. L. Lucieer, S. J. Wotherspoon, and C. R. Johnson. 2020. Multi-decadal decline in cover of giant kelp Macrocystis pyrifera at the southern limit of its Australian range. Marine ecology progress series 653:1–18.

Carr, M., J. Caselle, K. Cavanaugh, J. Fiewald, K. Kroeker, D. Pondella, Tissot, B, Malone, D, A. Parsons-Field, and B. Spiecker. 2021. Monitoring and evaluation of kelp forest ecosystems in the MLPA marine protected area network. Technical report submitted to: California Seagrant, Ocean Protection Council Marine Protected Areas (MPA) Monitoring Program, California DEpartment of Fish and Wildlife Marine Resources Division.

Carr, M. H., J. E. Caselle, K. D. Koehn, and D. P. Malone. 2020. PISCO Kelp Forest Community Surveys.

Carr, M. H., and D. C. Reed. 2016. Chapter 17: Shallow rocky reefs and kelp forests. Pages 311–336 in H. Mooney and E. Zavaleta, editors. Ecosystems of California. University of California Press, Berkeley.

Castorani, M. C. N., T. W. Bell, J. A. Walter, D. C. Reuman, K. C. Cavanaugh, and L. W. Sheppard. 2022. Disturbance and nutrients synchronise kelp forests across scales through interacting Moran effects. Ecology letters 25:1854–1868.

Castorani, M. C. N., D. C. Reed, F. Alberto, T. W. Bell, R. D. Simons, K. C. Cavanaugh, D. A. Siegel, and P. T. Raimondi. 2015. Connectivity structures local population dynamics: a long-term empirical test in a large metapopulation system. Ecology 96:3141–3152.

Castorani, M. C. N., D. C. Reed, P. T. Raimondi, F. Alberto, T. W. Bell, K. C. Cavanaugh, D. A. Siegel, and R. D. Simons. 2017. Fluctuations in population fecundity drive variation in demographic connectivity and metapopulation dynamics. Proceedings. Biological sciences / The Royal Society 284:20162086.

Cavanaugh, K. C., T. Bell, M. Costa, N. E. Eddy, L. Gendall, M. G. Gleason, M. Hessing-Lewis, R. Martone, M. McPherson, O. Pontier, L. Reshitnyk, R. Beas-Luna, M. Carr, J. E. Caselle, K. C. Cavanaugh, R. Flores Miller, S. Hamilton, W. N. Heady, H. K. Hirsh, R. Hohman, L. C. Lee, J. Lorda, J. Ray, D. C. Reed, V. R. Saccomanno, and S. B. Schroeder. 2021. A Review of the Opportunities and Challenges for Using Remote Sensing for Management of Surface-Canopy Forming Kelps. Frontiers in Marine Science 8:1536.

Cavanaugh, K. C., K. C. Cavanaugh, C. C. Pawlak, T. W. Bell, and V. R. Saccomanno. 2023. CubeSats show persistence of bull kelp refugia amidst a regional collapse in California. Remote sensing of environment 290:113521.

Cavanaugh, K. C., D. C. Reed, T. W. Bell, M. C. N. Castorani, R. Beas-luna, and N. S. Barrett. 2019. Spatial variability in the resistance and resilience of giant kelp in southern and Baja California to a multiyear heatwave. Frontiers in Marine Science 6:1–14.

Claisse, J. T., C. A. Blanchette, J. E. Dugan, J. P. Williams, J. Freiwald, D. J. Pondella II, N. K. Schooler, D. M. Hubbard, K. Davis, L. A. Zahn, C. M. Williams, and J. E. Caselle. 2018. Biogeographic patterns of communities across diverse marine ecosystems in southern California. Marine ecology 39:e12453.

Dayton, P. K. 1972. Toward an understanding of community resilience and the potential effects of enrichments to the benthos at McMurdo Sound, Antarctica. Pages 81–96 Proceedings of the colloquium on conservation problems in Antarctica. Allen Press, Lawrence, Kansas.

Dayton, P. K., V. Currie, T. Gerrodette, B. D. Keller, R. Rosenthal, and D. V. Tresca. 1984. Patch Dynamics and Stability of Some California Kelp Communities. Ecological monographs 54:254–289.

Dayton, P. K., M. J. Tegner, P. B. Edwards, and K. L. Riser. 1999. Temporal and spatial scales of kelp demography: The role of oceanographic climate. Ecological monographs 69:219–250.

Dean, T. A. 1985. The temporal and spatial distribution of underwater quantum irradiation in a southern California kelp forest. Estuarine, coastal and shelf science 21:835–844.

De’ath, G., K. E. Fabricius, H. Sweatman, and M. Puotinen. 2012. The 27-year decline of coral cover on the Great Barrier Reef and its causes. Proceedings of the National Academy of Sciences of the United States of America 109:17995–17999.

Dobkowski, K. A., K. D. Flanagan, and J. R. Nordstrom. 2019. Factors influencing recruitment and appearance of bull kelp, Nereocystis luetkeana (phylum Ochrophyta). Journal of phycology 55:236–244.

Duggins, D., J. E. Eckman, C. E. Siddon, and T. Klinger. 2001. Interactive roles of mesograzers and current flow in survival of kelps. Marine ecology progress series 223:143–155.

Edwards, M. S., and J. A. Estes. 2006. Catastrophe, recovery and range limitation in NE Pacific kelp forests: a large-scale perspective. Marine ecology progress series 320:79–87.

Eger, A. M., C. Layton, T. A. McHugh, M. Gleason, and N. Eddy. 2022. Kelp Restoration Guidebook: Lessons Learned from Kelp Projects Around the World.

Eger, A. M., E. M. Marzinelli, R. Beas-Luna, C. O. Blain, L. K. Blamey, J. E. K. Byrnes, P. E. Carnell, C. G. Choi, M. Hessing-Lewis, K. Y. Kim, N. H. Kumagai, J. Lorda, P. Moore, Y. Nakamura, A. Pérez-Matus, O. Pontier, D. Smale, P. D. Steinberg, and A. Vergés. 2023. The value of ecosystem services in global marine kelp forests. Nature communications 14:1894.

Ellison, A. M., M. S. Bank, B. D. Clinton, E. A. Colburn, K. Elliott, C. R. Ford, D. R. Foster, B. D. Kloeppel, J. D. Knoepp, G. M. Lovett, J. Mohan, D. A. Orwig, N. L. Rodenhouse, W. V. Sobczak, K. A. Stinson, J. K. Stone, C. M. Swan, J. Thompson, B. Von Holle, and J. R. Webster. 2005. Loss of foundation species: consequences for the structure and dynamics of forested ecosystems. Frontiers in ecology and the environment 3:479–486.

Filbee-Dexter, K., and R. E. Scheibling. 2014. Sea urchin barrens as alternative stable states of collapsed kelp ecosystems. Marine Ecology Progress Series 495:1–25.

Fisher, R. 2022. FSSgam: Full subsets multiple regresssion in R using gam(m4).

García-Reyes, M., J. L. Largier, and W. J. Sydeman. 2014. Synoptic-scale upwelling indices and predictions of phyto- and zooplankton populations. Progress in oceanography 120:177–188.

García-Reyes, M., S. A. Thompson, L. Rogers-Bennett, and W. J. Sydeman. 2022. Winter oceanographic conditions predict summer bull kelp canopy cover in northern California. PloS one 17:e0267737.

Gaylord, B., D. C. Reed, P. T. Raimondi, and L. Washburn. 2006. Macroalgal spore dispersal in coastal environments: Mechanistic insights revealed by theory and experiment. Ecological monographs 76:481–502.

Gerard, V. A. 1982. In situ rates of nitrate uptake by giant kelp, Macrocystis Pyrifera (L.) C. Agardh: Tissue differences, environmental effects, and predictions of nitrogen-limited growth. Journal of experimental marine biology and ecology 62:211–224.

Gerard, V. A. 1984. The light environment in a giant kelp forest: influence of Macrocystis pyrifera on spatial and temporal variability. Marine biology 84:189–195.

Giraldo-Ospina, A., T. Bell, M. H. Carr, D. Malone, and J. E. Caselle. 2023a. Where, when and how? A guide to kelp restoration in California using spatio-temporal models of kelp dynamics. A report to California Seagrant and California Ocean Protection Council.

Giraldo-Ospina, A., J. Caselle, M. Carr, T. Bell, and D. Malone. 2023b. Biological and environmental predictors of kelp density (2004-2021).

Goldberg, N. A., and G. A. Kendrick. 2004. Effects of island groups, depth, and exposure to ocean waves on subtidal macroalgal assemblages in the Recherche Archipelago, Western Australia. Journal of phycology 40:631–641.

Gosnell, J. S., R. J. A. Macfarlan, N. T. Shears, and J. E. Caselle. 2014. A dynamic oceanographic front drives biogeographical structure in invertebrate settlement along Santa Cruz Island, California, USA. Marine ecology progress series 507:181–196.

Graham, M. H. 1997. Factors determining the upper limit of giant kelp, Macrocystis pyrifera Agardh, along the Monterey Peninsula, central California, USA. Journal of experimental marine biology and ecology 218:127–149.

Graham, M. H., J. A. Vásquez, and A. H. Buschmann. 2007. Global ecology of the giant kelp Macrocystis: From ecotypes to ecosystems. Oceanography and Marine Biology: An Annual Review 45:39–88.

Hamilton, S. L., J. E. Caselle, D. P. Malone, and M. H. Carr. 2010. Incorporating biogeography into evaluations of the Channel Islands marine reserve network. Proceedings of the National Academy of Sciences of the United States of America 107:18272–18277.

Hanski, I. 1998. Metapopulation dynamics. Nature 396:41–49.

Hijmans, R. 2022. terra: Spatial Data Analysis.

Hoegh-Guldberg, O., P. J. Mumby, A. J. Hooten, R. S. Steneck, P. Greenfield, E. Gomez, C. D. Harvell, P. F. Sale, A. J. Edwards, K. Caldeira, N. Knowlton, C. M. Eakin, R. Iglesias-Prieto, N. Muthiga, R. H. Bradbury, A. Dubi, and M. E. Hatziolos. 2007. Coral reefs under rapid climate change and ocean acidification. Science 318:1737–1742.

Horn, M. H., L. G. Allen, and R. N. Lea. 2006. Chapter 1: Biogeography. Pages 3–25 in L. G. Allen, D. J. Pondella, and M. H. Horn, editors. The Ecology of Marine Fishes: California and Adjacent Waters. Universtiy of California Press, Berkeley.

Howells, E. J., V. H. Beltran, N. W. Larsen, L. K. Bay, B. L. Willis, and M. J. H. van Oppen. 2011. Coral thermal tolerance shaped by local adaptation of photosymbionts. Nature climate change 2:116–120.

Hughes, B. B., R. Beas-Luna, A. K. Barner, K. Brewitt, D. R. Brumbaugh, E. B. Cerny-Chipman, S. L. Close, K. E. Coblentz, K. L. De Nesnera, S. T. Drobnitch, J. D. Figurski, B. Focht, M. Friedman, J. Freiwald, K. K. Heady, W. N. Heady, A. Hettinger, A. Johnson, K. A. Karr, B. Mahoney, M. M. Moritsch, A. M. K. Osterback, J. Reimer, J. Robinson, T. Rohrer, J. M. Rose, M. Sabal, L. M. Segui, C. Shen, J. Sullivan, R. Zuercher, P. T. Raimondi, B. A. Menge, K. Grorud-Colvert, M. Novak, and M. H. Carr. 2017. Long-Term studies contribute disproportionately to ecology and policy. Bioscience 67:271–278.

Hughes, T. P., A. H. Baird, D. R. Bellwood, M. Card, S. R. Connolly, C. Folke, R. Grosberg, O. Hoegh-Guldberg, J. B. C. Jackson, J. Kleypas, J. M. Lough, P. Marshall, M. Nyström, S. R. Palumbi, J. M. Pandolfi, B. Rosen, and J. Roughgarden. 2003. Climate change, human impacts, and the resilience of coral reefs. Science 301:929–933.

Hurd, C. L. 2000. Water motion, marine macroalgal physiology, and production. Journal of phycology 36:453–472.

Huyer, A. 1983. Coastal upwelling in the California current system. Progress in oceanography 12:259–284.

Hynes, S., W. Chen, K. Vondolia, C. Armstrong, and E. O’Connor. 2021. Valuing the ecosystem service benefits from kelp forest restoration: A choice experiment from Norway. Ecological economics: the journal of the International Society for Ecological Economics 179:106833.

Jackson, G. A. 1977. Nutrients and production of giant kelp, Macrocystis pyrifera, off southern California1. Limnology and oceanography 22:979–995.

Johnson, A., and M. Koehl. 1994. Maintenance of Dynamic Strain Similarity and Environmental Stress Factor in Different Flow Habitats: Thallus Allometry and Material Properties of a Giant Kelp. The Journal of experimental biology 195:381–410.

Johnson, C. R., S. C. Banks, N. S. Barrett, F. Cazassus, P. K. Dunstan, G. J. Edgar, S. D. Frusher, C. Gardner, M. Haddon, F. Helidoniotis, K. L. Hill, N. J. Holbrook, G. W. Hosie, P. R. Last, S. D. Ling, J. Melbourne-Thomas, K. Miller, G. T. Pecl, A. J. Richardson, K. R. Ridgway, S. R. Rintoul, D. A. Ritz, D. J. Ross, J. C. Sanderson, S. A. Shepherd, A. Slotwinski, K. M. Swadling, and N. Taw. 2011. Climate change cascades: Shifts in oceanography, species’ ranges and subtidal marine community dynamics in eastern Tasmania. Journal of experimental marine biology and ecology 400:17–32.

Jones, C. G., J. H. Lawton, and M. Shachak. 1994. Organisms as ecosystem engineers. Oikos 69:373–386.

Karatayev, V. A., M. L. Baskett, D. J. Kushner, N. T. Shears, J. E. Caselle, and C. Boettiger. 2021. Grazer behaviour can regulate large-scale patterning of community states. Ecology letters 24:1917–1929.

Kavanaugh, M. T., K. J. Nielsen, F. T. Chan, B. A. Menge, R. M. Letelier, and L. M. Goodrich. 2009. Experimental assessment of the effects of shade on an intertidal kelp: Do phytoplankton blooms inhibit growth of open coast macroalgae? Limnology and oceanography 54:276–288.

Koch, M., G. Bowes, C. Ross, and X.-H. Zhang. 2013. Climate change and ocean acidification effects on seagrasses and marine macroalgae. Global change biology 19:103–132.

Koehl, M. A. R., and R. S. Alberte. 1988. Flow, flapping, and photosynthesis ofNereocystis leutkeana: a functional comparison of undulate and flat blade morphologies. Marine biology 99:435–444.

Kopczak, C. D., R. C. Zimmerman, and J. N. Kremer. 1991. Variation in nitrogen physiology and growth among geographically isolated populations of the giant kelp, Macrocystis pyrifera (phaeophyta). Journal of phycology 27:149–158.

Krumhansl, K. A., D. K. Okamoto, A. Rassweiler, M. Novak, J. J. Bolton, K. C. Cavanaugh, S. D. Connell, C. R. Johnson, B. Konar, S. D. Ling, F. Micheli, K. M. Norderhaug, A. Pérez-Matus, I. Sousa-Pinto, D. C. Reed, A. K. Salomon, N. T. Shears, T. Wernberg, R. J. Anderson, N. S. Barrett, A. H. Buschmann, M. H. Carr, J. E. Caselle, S. Derrien-Courtel, G. J. Edgar, M. Edwards, J. A. Estes, C. Goodwin, M. C. Kenner, D. J. Kushner, F. E. Moy, J. Nunn, R. S. Steneck, J. Vásquez, J. Watson, J. D. Witman, and J. E. K. Byrnes. 2016. Global patterns of kelp forest change over the past half-century. Proceedings of the National Academy of Sciences of the United States of America 113:13785–13790.

Ladah, L. B., and J. A. Zertuche-González. 2007. Survival of microscopic stages of a perennial kelp (Macrocystis pyrifera) from the center and the southern extreme of its range in the Northern Hemisphere after exposure to simulated El Niño stress. Marine biology 152:677–686.

Ling, S. D., and J. P. Keane. 2024. Climate-driven invasion and incipient warnings of kelp ecosystem collapse. Nature communications 15:400.

Ling, S. D., N. Kriegisch, B. Woolley, and S. E. Reeves. 2019. Density-dependent feedbacks, hysteresis, and demography of overgrazing sea urchins. Ecology 100:e02577.

Ling, S. D., R. E. Scheibling, A. Rassweiler, C. R. Johnson, N. Shears, S. D. Connell, A. K. Salomon, K. M. Norderhaug, A. Pérez-Matus, J. C. Hernández, S. Clemente, L. K. Blamey, B. Hereu, E. Ballesteros, E. Sala, J. Garrabou, E. Cebrian, M. Zabala, D. Fujita, and L. E. Johnson. 2015. Global regime shift dynamics of catastrophic sea urchin overgrazing. Philosophical transactions of the Royal Society of London. Series B, Biological sciences 370:20130269.

Magurran, A. E., S. R. Baillie, S. T. Buckland, J. M. Dick, D. A. Elston, E. M. Scott, R. I. Smith, P. J. Somerfield, and A. D. Watt. 2010. Long-term datasets in biodiversity research and monitoring: assessing change in ecological communities through time. Trends in ecology & evolution 25:574–582.

Malone, D. P., K. Davis, S. I. Lonhart, A. Parsons-Field, J. E. Caselle, and M. H. Carr. 2022. Large-scale, multidecade monitoring data from kelp forest ecosystems in California and Oregon (USA). Ecology 103:e3630.

McPherson, M. L., D. J. I. Finger, H. F. Houskeeper, T. W. Bell, M. H. Carr, L. Rogers-Bennett, and R. M. Kudela. 2021. Large-scale shift in the structure of a kelp forest ecosystem co-occurs with an epizootic and marine heatwave. Communications biology 4:298.

Michaud, K. M., D. C. Reed, and R. J. Miller. 2022. The Blob marine heatwave transforms California kelp forest ecosystems. Communications biology 5:1143.

Miller, K. I., C. A. Balemi, D. R. Bell, C. O. Blain, P. E. Caiger, B. J. Hanns, S. E. Kulins, O. Peleg, A. J. P. Spyksma, and N. T. Shears. 2023. Large-scale one-off sea urchin removal promotes rapid kelp recovery in urchin barrens. Restoration Ecology.

Morris, R. L., R. Hale, E. M. A. Strain, S. E. Reeves, A. Vergés, E. M. Marzinelli, C. Layton, V. Shelamoff, T. D. J. Graham, M. Chevalier, and S. E. Swearer. (n.d.). Key Principles for Managing Recovery of Kelp Forests through Restoration.

North, W. J., and R. C. Zimmerman. 1984. Influences of macronutrients and water temperatures on summertime survival of Macrocystis canopies. Pages 419–424 Eleventh International Seaweed Symposium. Springer Netherlands.

Okamoto, D. K., S. C. Schroeter, and D. C. Reed. 2020. Effects of ocean climate on spatiotemporal variation in sea urchin settlement and recruitment. Limnology and oceanography 65:2076–2091.

Randell, Z., M. Kenner, J. Tomoleoni, J. Yee, and M. Novak. 2022. Kelp-forest dynamics controlled by substrate complexity. Proceedings of the National Academy of Sciences of the United States of America 119:e2103483119.

Reed, D. C., and M. S. Foster. 1984. The Effects of Canopy Shadings on Algal Recruitment and Growth in a Giant Kelp Forest. Ecology 65:937–948.

Reed, D. C., A. Rassweiler, and K. K. Arkema. 2008. Biomass rather than growth rate determines variation in net primary production by giant kelp. Ecology 89:2493–2505.

Reed, D. C., A. Rassweiler, M. H. Carr, K. C. Cavanaugh, D. P. Malone, and D. A. Siegel. 2011. Wave disturbance overwhelms top-down and bottom-up control of primary production in California kelp forests. Ecology 92:2108–2116.

Rennick, M., B. P. DiFiore, J. Curtis, D. C. Reed, and A. C. Stier. 2022. Detrital supply suppresses deforestation to maintain healthy kelp forest ecosystems. Ecology 103:e3673.

Rodriguez, G. E., A. Rassweiler, D. C. Reed, and S. J. Holbrook. 2013. The importance of progressive senescence in the biomass dynam of giant kelp (Macrocystis pyrifera). Ecology 94:1848–1858.

Rogers-Bennett, L., and C. A. Catton. 2019. Marine heat wave and multiple stressors tip bull kelp forest to sea urchin barrens. Scientific reports 9:15050.

Sanford, E., and M. W. Kelly. 2011. Local adaptation in marine invertebrates. Annual review of marine science 3:509–535.

Schiel, D. R., and M. S. Foster. 2015. The biology and ecology of giant kelp forests. University of California Press.

Serrano, O., A. Arias-Ortiz, C. M. Duarte, G. A. Kendrick, and P. S. Lavery. 2021. Impact of Marine Heatwaves on Seagrass Ecosystems. Pages 345–364 in J. G. Canadell and R. B. Jackson, editors. Ecosystem Collapse and Climate Change. Springer International Publishing, Cham.

Seymour, R. J., M. J. Tegner, P. K. Dayton, and P. E. Parnell. 1989. Storm wave induced mortality of giant kelp, Macrocystis pyrifera, in Southern California. Estuarine, coastal and shelf science 28:277–292.

Short, F. T., and H. A. Neckles. 1999. The effects of global climate change on seagrasses. Aquatic botany 63:169–196.

Smith, J. G., D. Malone, and M. H. Carr. 2024. Consequences of kelp forest ecosystem shifts and predictors of persistence through multiple stressors. Proceedings. Biological sciences / The Royal Society 291:20232749.

Smith, J. G., J. Tomoleoni, M. Staedler, S. Lyon, J. Fujii, and M. T. Tinker. 2021. Behavioral responses across a mosaic of ecosystem states restructure a sea otter-urchin trophic cascade. Proceedings of the National Academy of Sciences of the United States of America 118:e2012493118.

Snyder, J. N., T. W. Bell, D. A. Siegel, N. J. Nidzieko, and K. C. Cavanaugh. 2020. Sea Surface Temperature Imagery Elucidates Spatiotemporal Nutrient Patterns for Offshore Kelp Aquaculture Siting in the Southern California Bight. Frontiers in Marine Science 7:22.

Springer, Y. P., C. G. Hays, M. H. Carr, and M. R. Mackey. 2010. Toward ecosystem-based management of marine macroalgae—the bull kelp, nereocystis luetkeana. Pages 1–42 Oceanography and Marine Biology. Chapman and Hall/CRC.

Steneck, R. S., M. H. Graham, B. J. Bourque, D. Corbett, J. M. Erlandson, J. A. Estes, and M. J. Tegner. 2002. Kelp forest ecosystems: Biodiversity, stability, resilience and future. Environmental conservation 29:436–459.

Stewart, H. L., J. P. Fram, D. C. Reed, S. L. Williams, M. A. Brzezinski, S. MacIntyre, and B. Gaylord. 2009. Differences in growth, morphology and tissue carbon and nitrogen of Macrocystis pyrifera within and at the outer edge of a giant kelp forest in California, USA. Marine ecology progress series 375:101–112.

Strub, P. T., and T. M. Powell. 1987. Surface temperature and transport in Lake Tahoe: inferences from satellite (AVHRR) imagery. Continental shelf research 7:1001–1013.

Supratya, V. P., L. J. M. Coleman, and P. T. Martone. 2020. Elevated Temperature Affects Phenotypic Plasticity in the Bull Kelp (Nereocystis luetkeana, Phaeophyceae). Journal of phycology 56:1534–1541.

Vadas, R. L. 1972. Ecological implications of culture studies on nereocystis luetkeana. Journal of phycology 8:196–203.

Vásquez, J. A., S. Zuñiga, F. Tala, N. Piaget, D. C. Rodríguez, and J. M. A. Vega. 2014. Economic valuation of kelp forests in northern Chile: values of goods and services of the ecosystem. Journal of applied phycology 26:1081–1088.

Ward, R. D., D. A. Friess, R. H. Day, and R. A. Mackenzie. 2016. Impacts of climate change on mangrove ecosystems: a region by region overview. Ecosystem health and sustainability 2:e01211.

Wei, Y., and V. Simko. 2021. R package “corrplot”: Visualization of a Correlation Matrix.

Wernberg, T., S. Bennett, R. C. Babocock, T. de Bettignies, K. Cure, M. Depczynski, F. Dufois, J. Fromont, C. J. Fulton, R. K. Hovey, E. S. Harvey, T. H. Holmes, G. A. Kendrick, B. Radford, J. Santana-Garcon, B. J. Saunders, D. A. Smale, M. S. Thomsen, C. A. Tuckett, F. Tuya, M. A. Vanderklift, and S. Wilson. 2016. Climate-driven regime shift of a temperate marine ecosystem. Science 353:169–172.

Wood, S. 2006. Generalized additive models: An introduction with R. CRC Press, Boca Raron, Florida.

Wood, S. N. 2011. Fast stable restricted maximum likelihood and marginal likelihood estimation of semiparametric generalized linear models. Journal of the Royal Statistical Society 73:3–36.

Young, M. A., K. Critchell, A. D. Miller, E. A. Treml, M. Sams, R. Carvalho, and D. Ierodiaconou. 2023. Mapping the impacts of multiple stressors on the decline in kelps along the coast of Victoria, Australia. Diversity & distributions 29:199–220.

Young, M., K. Cavanaugh, T. Bell, P. Raimondi, C. A. Edwards, P. T. Drake, L. Erikson, and C. Storlazzi. 2016. Environmental controls on spatial patterns in the long-term persistence of giant kelp in central California. Ecological monographs 86:45–60.

Zimmerman, R. C., and J. N. Kremer. 1984. Episodic Nutrient Supply to a Kelp Forest Ecosystem in Southern California. Journal of Marine Research 42:591–604.

Zimmerman, R. C., and D. L. Robertson. 1985. Effects of El Niño on local hydrography and growth of the giant kelp,Macrocystis pyrifera, at Santa Catalina Island, California1. Limnology and oceanography 30:1298–1302.

