## Appendix for "Drivers of spatio-temporal variability in a marine foundation species"

**Appendix S1**

**Authors:** Anita Giraldo-Ospina, Tom Bell, Mark H. Carr, Jennifer E. Caselle

**Title:** Drivers of spatio-temporal variability in a marine foundation species

### List of the environmental variables tested as potential predictors

**Table S1.** List of the environmental variables tested as potential predictors in the models of kelp and all the different metrics for each variable for a total of 96 environmental variables tested for the final models of kelp density.

| Variable type | Variable metric | Units | Original spatial resolution |
|---|---|---|---|
| Temperature | Minimum | Degrees Celsius | 5 km |
| Temperature | Maximum | Degrees Celsius | 5 km |
| Temperature | Mean | Degrees Celsius | 5 km |
| Temperature | Minimum - summer season | Degrees Celsius | 5 km |
| Temperature | Maximum - summer season | Degrees Celsius | 5 km |
| Temperature | Mean - summer season | Degrees Celsius | 5 km |
| Temperature | Minimum - upwelling season | Degrees Celsius | 5 km |
| Temperature | Maximum - upwelling season | Degrees Celsius | 5 km |
| Temperature | Mean - upwelling season | Degrees Celsius | 5 km |
| Temperature | Minimum anomaly | Degrees Celsius | 5 km |
| Temperature | Maximum anomaly | Degrees Celsius | 5 km |
| Temperature | Mean anomaly | Degrees Celsius | 5 km |
| Temperature | Minimum anomaly - summer season | Degrees Celsius | 5 km |
| Temperature | Maximum anomaly - summer season | Degrees Celsius | 5 km |
| Temperature | Mean anomaly - summer season | Degrees Celsius | 5 km |

| | | | |
|---|---|---|---|
| Temperature | Minimum anomaly - upwelling season | Degrees Celsius | 5 km |
| Temperature | Maximum anomaly - upwelling season | Degrees Celsius | 5 km |
| Temperature | Mean anomaly - upwelling season | Degrees Celsius | 5 km |
| Temperature | Marine heatwave days | Degrees Celsius | 5 km |
| Temperature | Marine heatwave intensity | Degrees Celsius | 5 km |
| Temperature | Marine heatwave days - summer | Degrees Celsius | 5 km |
| Temperature | Marine heatwave intensity - summer | Degrees Celsius | 5 km |
| Temperature | Marine heatwave days - upwelling | Degrees Celsius | 5 km |
| Temperature | Marine heatwave intensity - upwelling | Degrees Celsius | 5 km |
| Temperature | Days above 15 degrees | Degrees Celsius | 5 km |
| Temperature | Days above 16 degrees | Degrees Celsius | 5 km |
| Temperature | Days above 17 degrees | Degrees Celsius | 5 km |
| Temperature | Days above 18 degrees | Degrees Celsius | 5 km |
| Temperature | Days above 19 degrees | Degrees Celsius | 5 km |
| Temperature | Days above 20 degrees | Degrees Celsius | 5 km |
| Temperature | Days above 21 degrees | Degrees Celsius | 5 km |
| Temperature | Days above 21 degrees | Degrees Celsius | 5 km |
| Temperature | Days above 22 degrees | Degrees Celsius | 5 km |
| Temperature | Days above 23 degrees | Degrees Celsius | 5 km |
| Temperature | Degree days 15 | Degrees Celsius | 5 km |
| Temperature | Degree days 16 | Degrees Celsius | 5 km |
| Temperature | Degree days 17 | Degrees Celsius | 5 km |
| Temperature | Degree days 18 | Degrees Celsius | 5 km |
| Temperature | Degree days 19 | Degrees Celsius | 5 km |
| Temperature | Degree days 20 | Degrees Celsius | 5 km |
| Temperature | Degree days 21 | Degrees Celsius | 5 km |
| Temperature | Degree days 21 | Degrees Celsius | 5 km |
| Temperature | Degree days 22 | Degrees Celsius | 5 km |

| | | | |
|---|---|---|---|
| Temperature | Degree days 23 | Degrees Celsius | 5 km |
| Nitrate | Minimum | µmol L-1 | 5 km |
| Nitrate | Maximum | µmol L-1 | 5 km |
| Nitrate | Mean | µmol L-1 | 5 km |
| Nitrate | Mean - upwelling season | µmol L-1 | 5 km |
| Nitrate | Mean - summer season | µmol L-1 | 5 km |
| Nitrate | Minimum anomaly | µmol L-1 | 5 km |
| Nitrate | Maximum anomaly | µmol L-1 | 5 km |
| Nitrate | Minimum anomaly - summer season | µmol L-1 | 5 km |
| Nitrate | Maximum anomaly - summer season | µmol L-1 | 5 km |
| Nitrate | Minimum anomaly - upwelling season | µmol L-1 | 5 km |
| Nitrate | Maximum anomaly - upwelling season | µmol L-1 | 5 km |
| Nitrate | Days above 1 | µmol L-1 | 5 km |
| Nitrate | Days above 2 | µmol L-1 | 5 km |
| Nitrate | Days above 3 | µmol L-1 | 5 km |
| Nitrate | Days above 4 | µmol L-1 | 5 km |
| Nitrate | Days above 5 | µmol L-1 | 5 km |
| Nitrate | Days above 6 | µmol L-1 | 5 km |
| Nitrate | Days above 7 | µmol L-1 | 5 km |
| Nitrate | Days above 8 | µmol L-1 | 5 km |
| Nitrate | Degree days 1 | µmol L-1 | 5 km |
| Nitrate | Degree days 2 | µmol L-1 | 5 km |
| Nitrate | Degree days 3 | µmol L-1 | 5 km |
| Nitrate | Degree days 4 | µmol L-1 | 5 km |
| Nitrate | Degree days 5 | µmol L-1 | 5 km |
| Nitrate | Degree days 6 | µmol L-1 | 5 km |
| Nitrate | Degree days 7 | µmol L-1 | 5 km |
| Nitrate | Degree days 8 | µmol L-1 | 5 km |

| | | | |
|---|---|---|---|
| NPP | Minimum | mg C m-2 d-1 | 4 km |
| NPP | Maximum | mg C m-2 d-1 | 4 km |
| NPP | Mean | mg C m-2 d-1 | 4 km |
| NPP | Minimum - summer season | mg C m-2 d-1 | 4 km |
| NPP | Maximum - summer season | mg C m-2 d-1 | 4 km |
| NPP | Mean - summer season | mg C m-2 d-1 | 4 km |
| NPP | Minimum - upwelling season | mg C m-2 d-1 | 4 km |
| NPP | Maximum - upwelling season | mg C m-2 d-1 | 4 km |
| NPP | Mean - upwelling season | mg C m-2 d-1 | 4 km |
| Wave height | Mean | m | 1 km |
| Wave height | Max | m | 1 km |
| Wave height | 95 percentile | m | 1 km |
| Wave height | 99 percentile | m | 1 km |
| Orbital velocity | Mean | m/s | 30 m |
| Orbital velocity | Max | m/s | 30 m |
| Depth | Mean | m | 30 m |
| Depth | Max | m | 30 m |
| Depth | Min | m | 30 m |
| Slope | Mean | radians | 30 m |
| Slope | Max | radians | 30 m |
| Slope | Min | radians | 30 m |
| Vector Ruggedness Measure | Mean | n/a | 30 m |
| Vector Ruggedness Measure | Max | n/a | 30 m |
| Vector Ruggedness Measure | Min | n/a | 30 m |

**Appendix S2**

**Authors:** Anita Giraldo-Ospina, Tom Bell, Mark H. Carr, Jennifer E. Caselle

**Title:** Drivers of spatio-temporal variability in a marine foundation species

**White-zone interpolation**

We obtained the fine-scale bathymetry for the northern Channel Islands directly from NOAA. The mapped fine-scale bathymetry of California is missing or limited in shallow waters (referred to as the 'white zone'). This zone extends between 50 m to 500 m offshore. It includes areas where navigation hazards impede the access of seafloor mapping vessels and where multibeam sonar is generally infeasible due to underwater hazards and/or dense kelp canopies. However, understanding seafloor terrain in the 'white zone' is necessary since this zone supports rich kelp forests and provides critical habitat for a myriad of recreational and commercially targeted species. To extend the bathymetry files into the 'white zone' we interpolated the area between the mapped bathymetry and the shoreline using a natural neighbor algorithm in ArcGIS 10.2 (ArcGIS 10.2, Esri Industries, Redlands CA). We first aggregated the fine-scale bathymetry (4 $m^2$ grid) onto a 900 $m^2$ grid using the mean depth. We then used shoreline data available as polyline data attributed with detailed shoreline habitat characteristics from the National Oceanic and Atmospheric Administration's (NOAA) Environmental Sensitivity Index (ESI) maps, which were simplified to a binary (rock/sediment) classification. These binary files were aggregated into 900 $m^2$ grid and then interpolated across the 'white zone' using a natural neighbor algorithm in ArcGIS 10.2 (ArcGIS 10.2, Esri Industries, Redlands CA). Substrate type information (hard vs. soft) was extended into the 'white zone' using binary shoreline files of rock vs. sand and interpolating from the edge of the mapped bathymetry into the shoreline data. To do this, we first aggregated the binary fine scale substrate data (4 $m^2$ grid) onto a 900 $m^2$ grid, by calculating the proportion of rock within each 900 $m^2$ cell of the grid (each 900 $m^2$ cell contains a total of 225 4 $m^2$ cells) and transforming the shoreline type vector files into binary 900 $m^2$ raster cells. Finally, an inverse distance weighting interpolation (IDW) with a weight of 0.5 was conducted between the aggregated substrate type rasters to the shoreline type raster using ArcGIS 10.2. The IDW interpolation was selected as it has been shown to provide accurate substrate type interpolations across a range of substrate configurations and white zone widths (Saarman et al. n.d.). This resulted in continuous depth and substrate type files that included a combination of mapped and interpolated bathymetry.

**Appendix S3**

**Authors:** Anita Giraldo-Ospina, Tom Bell, Mark H. Carr, Jennifer E. Caselle

**Title:** Drivers of spatio-temporal variability in a marine foundation species

**Zoospore availability and density calculations**

We created an annual layer that estimated surface canopy-forming kelp zoospore availability at any given pixel as a function of the maximum kelp biomass each year and an empirical zoospore dispersal function estimated by (Gaylord et al. 2006). Spatial layers of estimated annual zoospore availability were only estimated for giant kelp since an equivalent dispersal function for bull kelp did not exist at the time we produced this work. The framework to obtain annual spatial files of maximum zoospore density per year required 4 steps: 1) Estimate giant kelp fecundity ($cm^2$ of sorus area per $m^2$ of kelp biomass) from Landsat-derived kelp biomass using a non-linear relationship identified in previous work by (Castorani et al. 2017); 2) estimate number of spores released per fertile unit ($cm^2$ of sorus area per $m^2$ of kelp biomass); 3) estimate the dispersal distances of spores for each pixel using a dispersal curve for giant kelp spores by (Gaylord et al. 2006), and construct a map of zoospore dispersal densities; and 4) filtering the maps of zoospore density to areas with adequate substrate (rock).

First we obtained spatial data of giant kelp biomass (central and south coasts of California) for semester 1 and semester 2 from the year 2003 to 2020 derived from Landsat imagery. Kelp biomass from Landsat is updated quarterly, so to obtain one value per semester, we extracted the maximum kelp biomass between quarter 1 and quarter 2 (semester 1) and quarter 3 and quarter 4 (semester 2). Landsat data is obtained at 30 m resolution which results in a value of kelp biomass per pixel of kg/900 $m^2$. We then obtained a biomass value per $m^2$ by dividing the kelp biomass of each pixel by 900 and used this biomass value to estimate giant kelp fecundity ($cm^2$ of sorus area per $m^2$ of kelp biomass) using a non-linear relationship identified in previous work by (Castorani et al. 2017). The relationship between pixel fecundity ($cm^2$ sorus per $m^2$) and the square root of pixel canopy biomass density (kg wet per $m^2$) in the first semester of the year (January to June, Fecundity density = 1463 × √ canopy biomass density) was found to be different from the second semester of the year (July – December, Fecundity density = 609.6 × √ canopy biomass density) (Castorani et al. 2017), so for each semester of the year the adequate equation was applied to the spatial data of kelp biomass, to transform it into fecundity. We then had an estimate of sorus area per $m^2$, and we transformed it into zoospores per $m^2$ by using estimates from (Anderson and North 1966) for giant kelp where they found that there were up to 35 million of zoospores per $cm^2$ of sorus. We then scaled up this to 900 $m^2$ to obtain an estimate of zoospores number per pixel (30 x 30 m), per semester of the year.

With the number of spores per pixel, we then estimate their potential dispersal by using a lognormal plus Gaussian dispersal curve (equation S4.1), where p is the probability of dispersal to a particular radial distance r, and a, b and c are fitted parameters (a = 2.03, b = 27.8, c = 1060). The dispersal curve is the sum of a lognormal and a normal probability density function, each scaled to ensure that their sum integrates to 1, and which was found to be the most appropriate to describe the dispersal of giant kelp zoospores at different current speeds and directions by (Gaylord et al. 2006). To achieve this, we used a dispersal kernel analysis, using the 'kernel2dsmooth' function of the R package 'smoothie' to apply the dispersal curve to each spatial file of zoospores per pixel resulting in a series of maps of zoospore dispersal densities for both semesters of the years 2003 to 2020.

$$p(r) = \frac{1}{2a\sqrt{2\Pi r}}\, exp\left[-\frac{(lnr - lnb)^2}{2a^2}\right] + \frac{1}{c\sqrt{2\Pi}}\, exp\left(-\frac{r^2}{2c^2}\right)$$

(S4.1)

The dispersal densities of both semesters for every year were summed to obtain a single value of dispersal density resulting in a series of maps of zoospore number per 900 $m^2$ for every year from 2003 to 2020. Then, we filtered the annual maps of zoospore density and removed areas where the proportion of rock was less than 0.1, as zoospores need to settle in adequate substrate to survive, metamorphose into gametophytes, fertilize, and grow into sporophytes. Finally, we aggregated the maps from 30 x 30 m resolution to 300 x 300 m resolution by summing the zoospore densities.

The zoospore density data was extracted for every site and year used to build our models of giant kelp density and was included in the model as a predictor with a lag effect variable, so that the zoospore density calculated for year 1 (2003), was used as a predictor for kelp density in year 2 (2004).

Anderson, E. K., and W. J. North. 1966. “In Situ Studies of Spore Production and Dispersal in the Giant Kelp, Macrocystis.” *Proceedings of the Internation Seaweed Symposium*, no. 5: 73–86.

Castorani, Max C. N., Daniel C. Reed, Peter T. Raimondi, Filipe Alberto, Tom W. Bell, Kyle C. Cavanaugh, David A. Siegel, and Rachel D. Simons. 2017. “Fluctuations in Population Fecundity Drive Variation in Demographic Connectivity and Metapopulation Dynamics.” *Proceedings of the Royal Society B: Biological Sciences* 284 (1847). https://doi.org/10.1098/rspb.2016.2086.

Gaylord, Brian, Daniel C. Reed, Peter T. Raimondi, and Libe Washburn. 2006. “Macroalgal Spore Dispersal in Coastal Environments: Mechanistic Insights Revealed by Theory and Experiment.” *Ecological Monographs* 76 (4): 481–502.

**Appendix S4**

**Authors:** Anita Giraldo-Ospina, Tom Bell, Mark H. Carr, Jennifer E. Caselle

**Title:** Drivers of spatio-temporal variability in a marine foundation species

**Purple sea urchin spatio-temporal model results and reconstructed maps**

**Table S1.** Summary statistics of predictive performance of the GAMs developed to describe the drivers of purple urchin density dynamics in California.

| **Species** | **Region** | **Model** | **$r^2$** | **Deviance explained** |
|---|---|---|---|---|
| *S. purpuratus* | North | Days 10N +<br>maximum temperature anomaly upwelling +<br>max orbital velocity (log) +<br>mean NPP (log) +<br>minimum NPP (log) +<br>mean depth | 0.79 | 63.29% |
| *S. purpuratus* | Central | Maximum nitrate anomaly summer +<br>minimum temperature +<br>days 16C (log)<br>mean wave height +<br>max NPP +<br>mean slope | 0.88 | 75.45% |
| *S. purpuratus* | South | Minimum nitrate +<br>days 8N (log) +<br>maximum temperature anomaly upwelling<br>+ days 20C (log) +<br>mean wave height +<br>min NPP upwelling (log) +<br>mean depth | 0.63 | 52.80% |

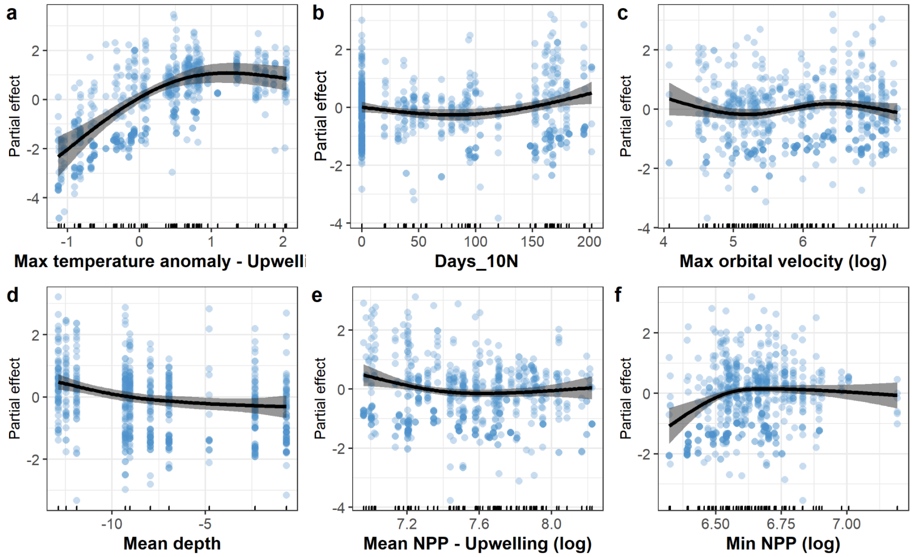

**FIGURE S1.** Partial response of *S. purpuraturs* density to each of the explanatory variables in the selected model for the north coast. Solid lines indicate the mean fitted values from cubic regression splines with other predictors held constant. Gray bands indicate 95% confidence intervals.

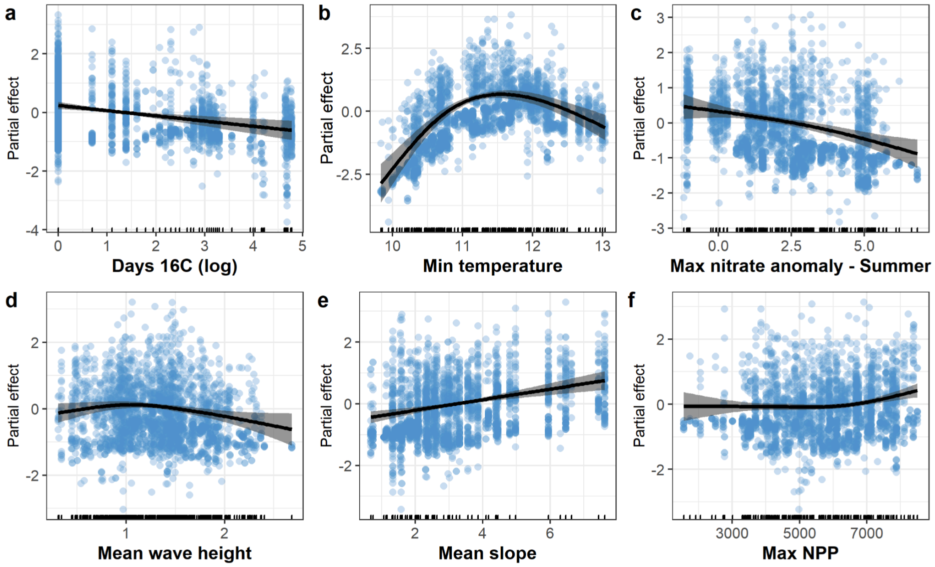

**FIGURE S2.** Partial response of *S. purpuraturs* density to each of the explanatory variables in the selected model for the central coast. Solid lines indicate the mean fitted values from cubic regression splines with other predictors held constant. Gray bands indicate 95% confidence intervals.

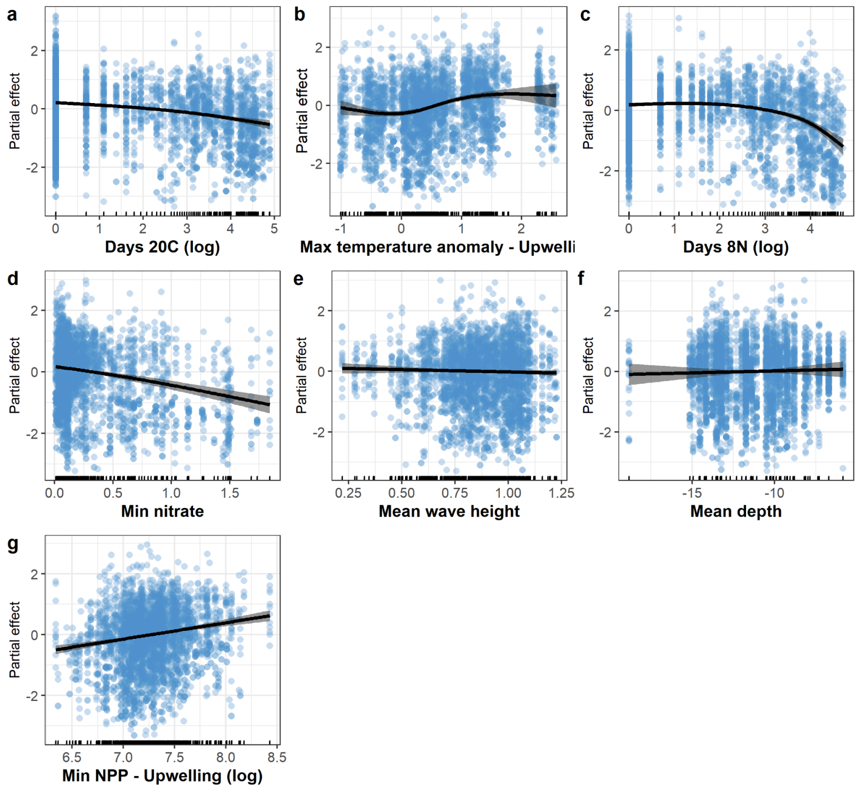

**FIGURE S3.** Partial response of *S. purpuraturs* density to each of the explanatory variables in the selected model for the south coast. Solid lines indicate the mean fitted values from cubic regression splines with other predictors held constant. Gray bands indicate 95% confidence intervals.

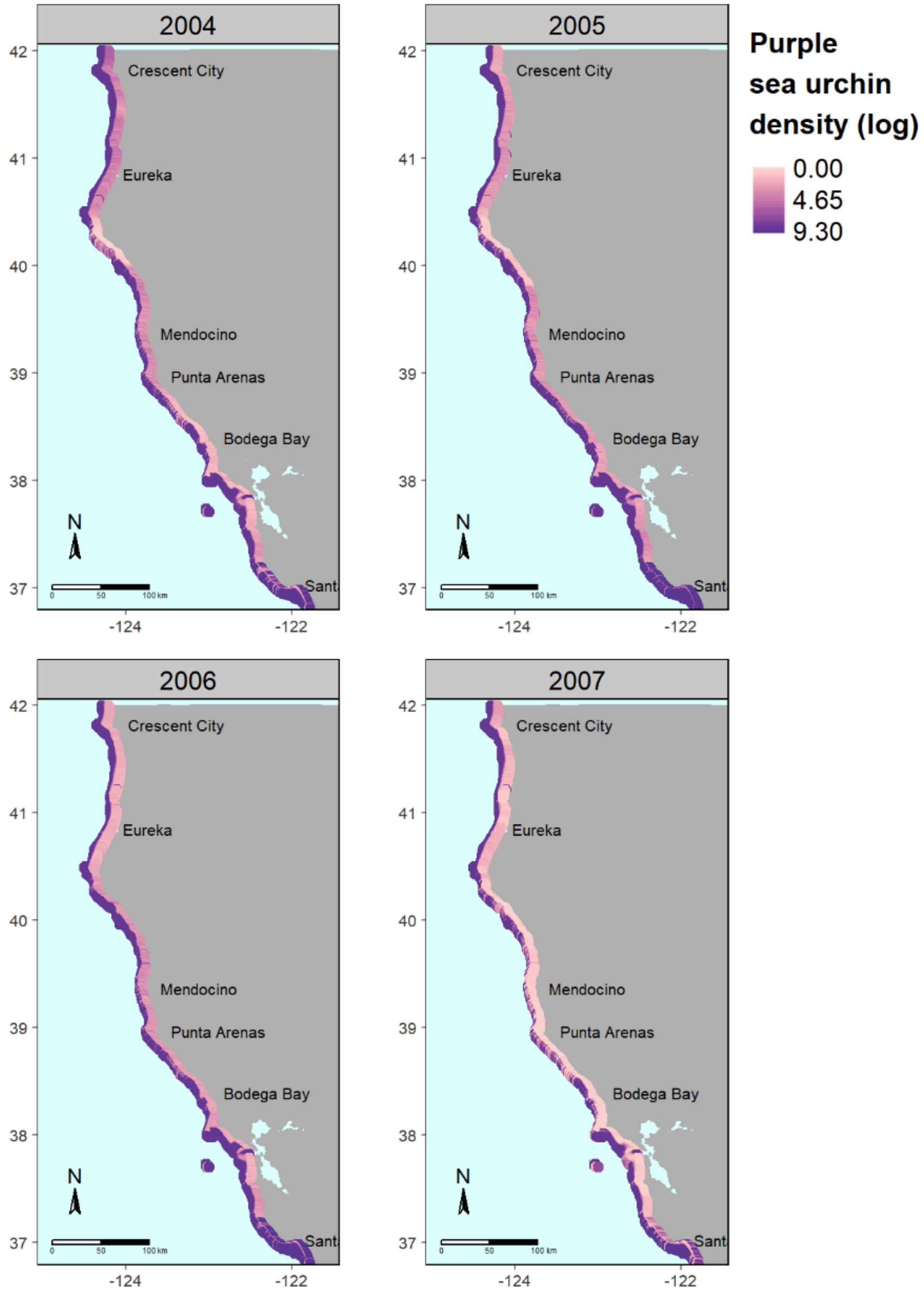

**FIGURE S4.** Predicted density of purple sea urchins (log urchins/60 $m^2$) in northern California from 2004 to 2007.

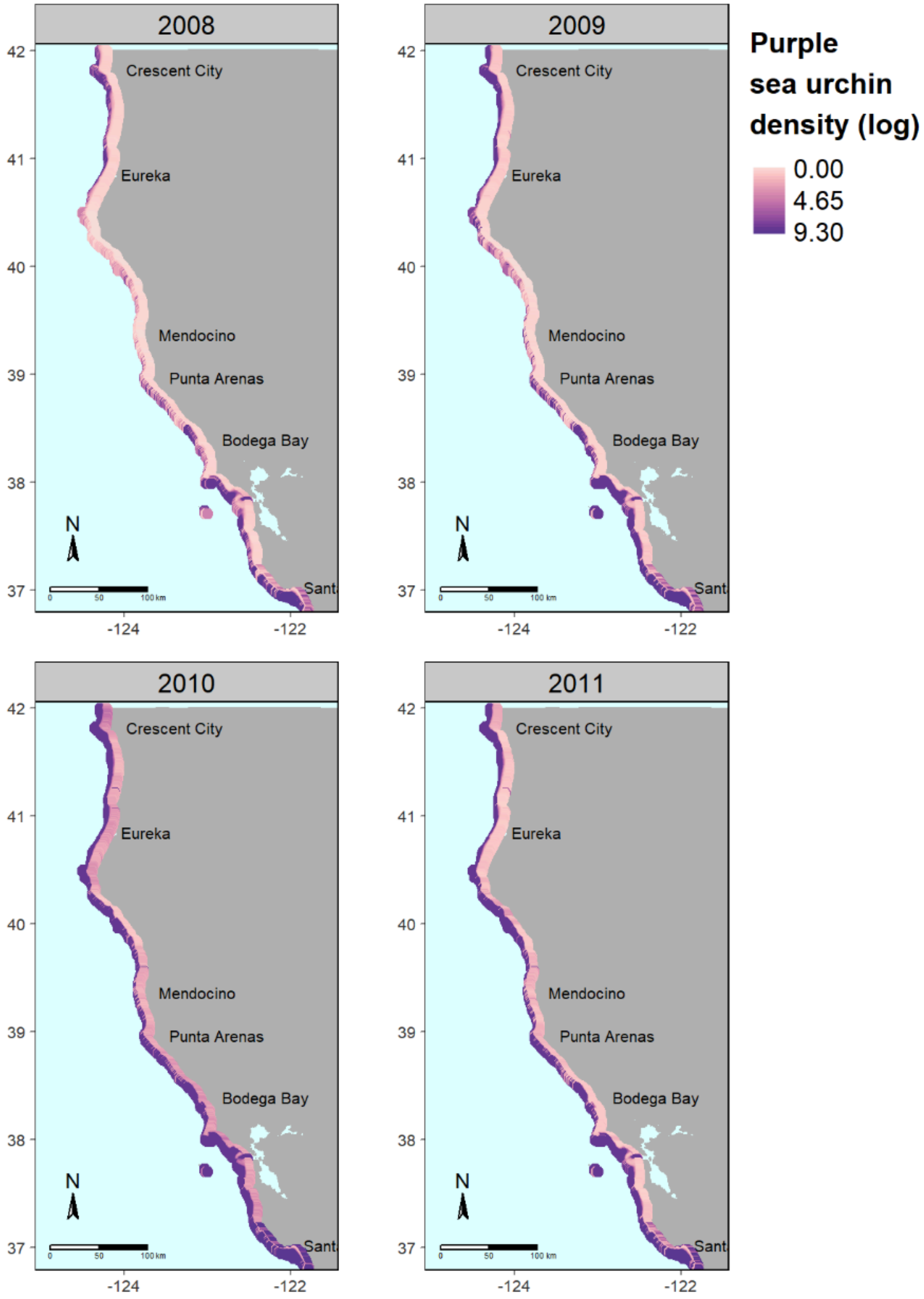

**FIGURE S5.** Predicted density of purple sea urchins (log urchins/60 $m^2$) in northern California from 2008 to 2011.

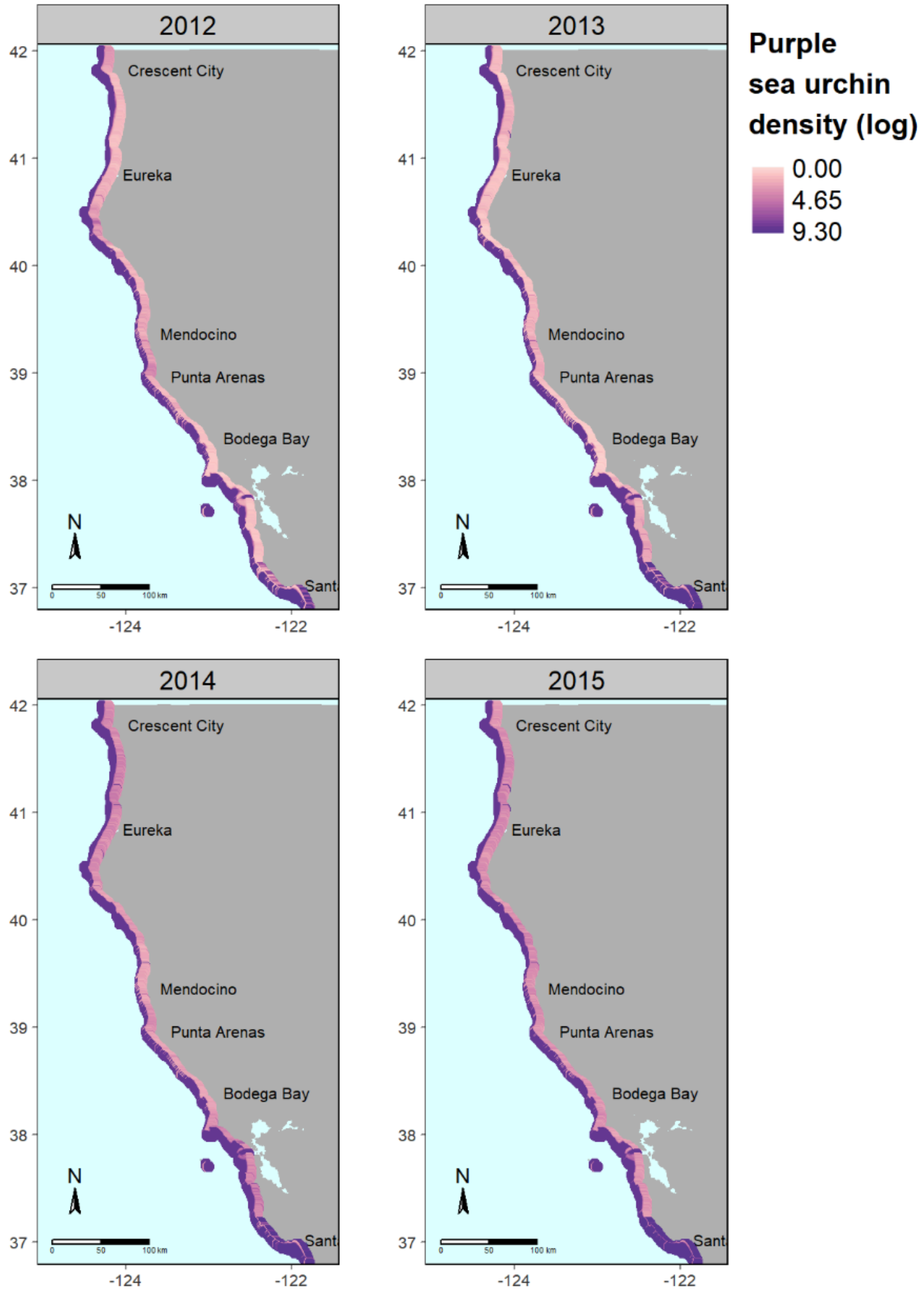

**FIGURE S6.** Predicted density of purple sea urchins (log urchins/60 $m^2$) in northern California from 2012 to 2015.

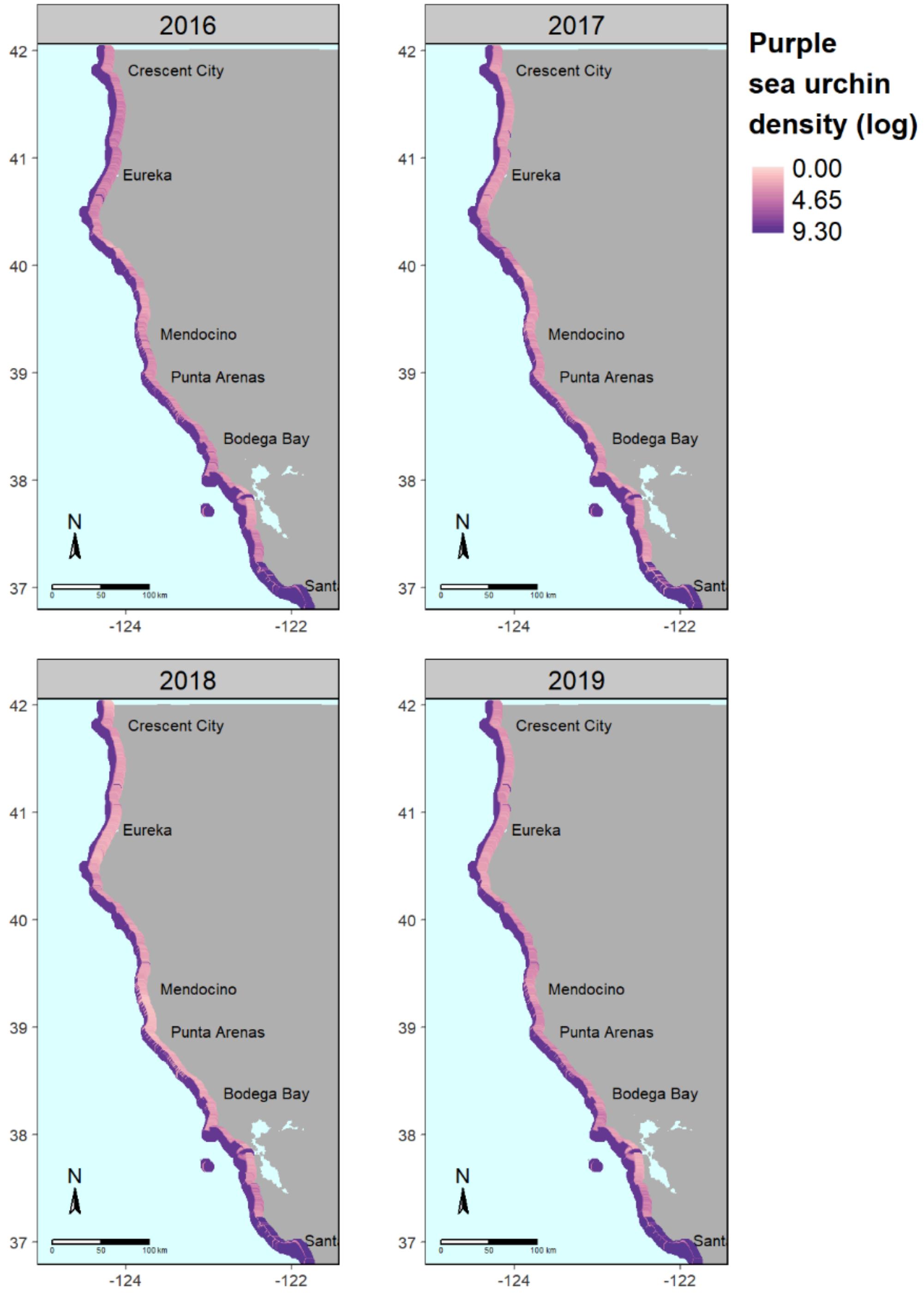

**FIGURE S7.** Predicted density of purple sea urchins (log urchins/60 $m^2$) in northern California from 2016 to 2019.

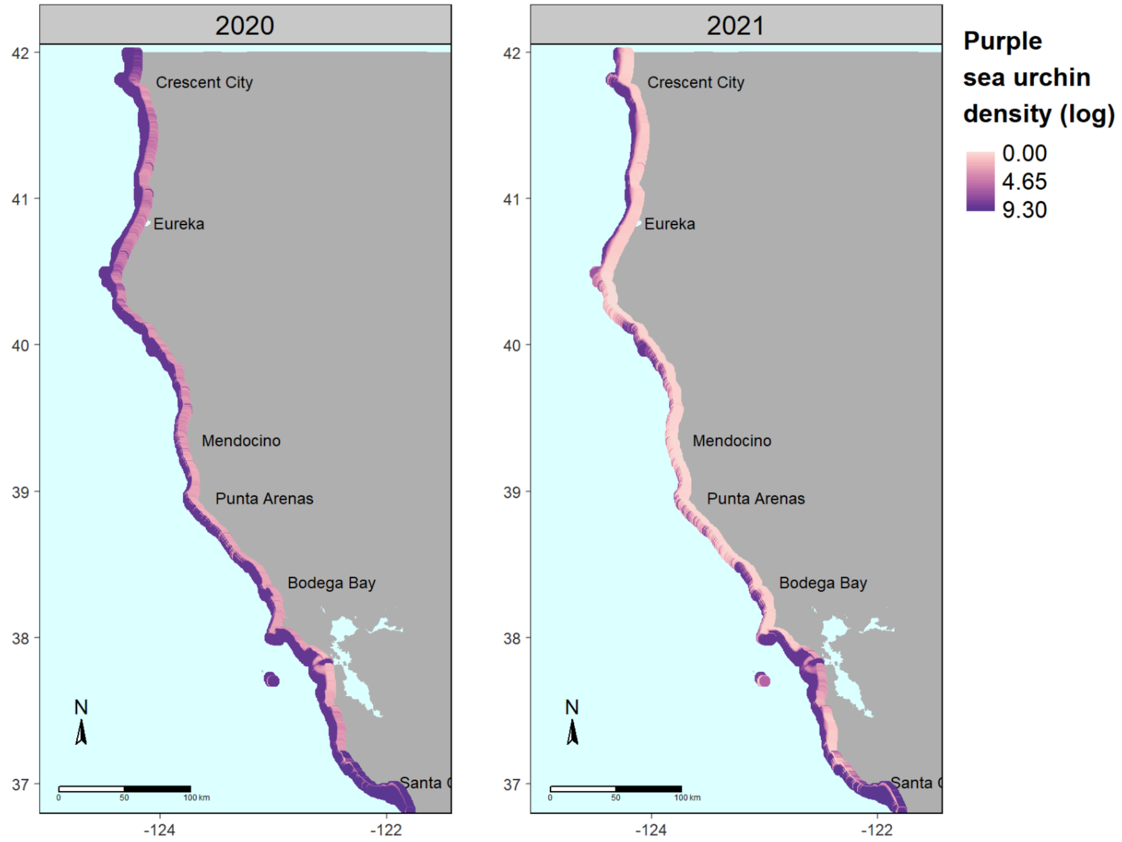

**FIGURE S8.** Predicted density of purple sea urchins (log urchins/60 $m^2$) in northern California from 2020 to 2021.

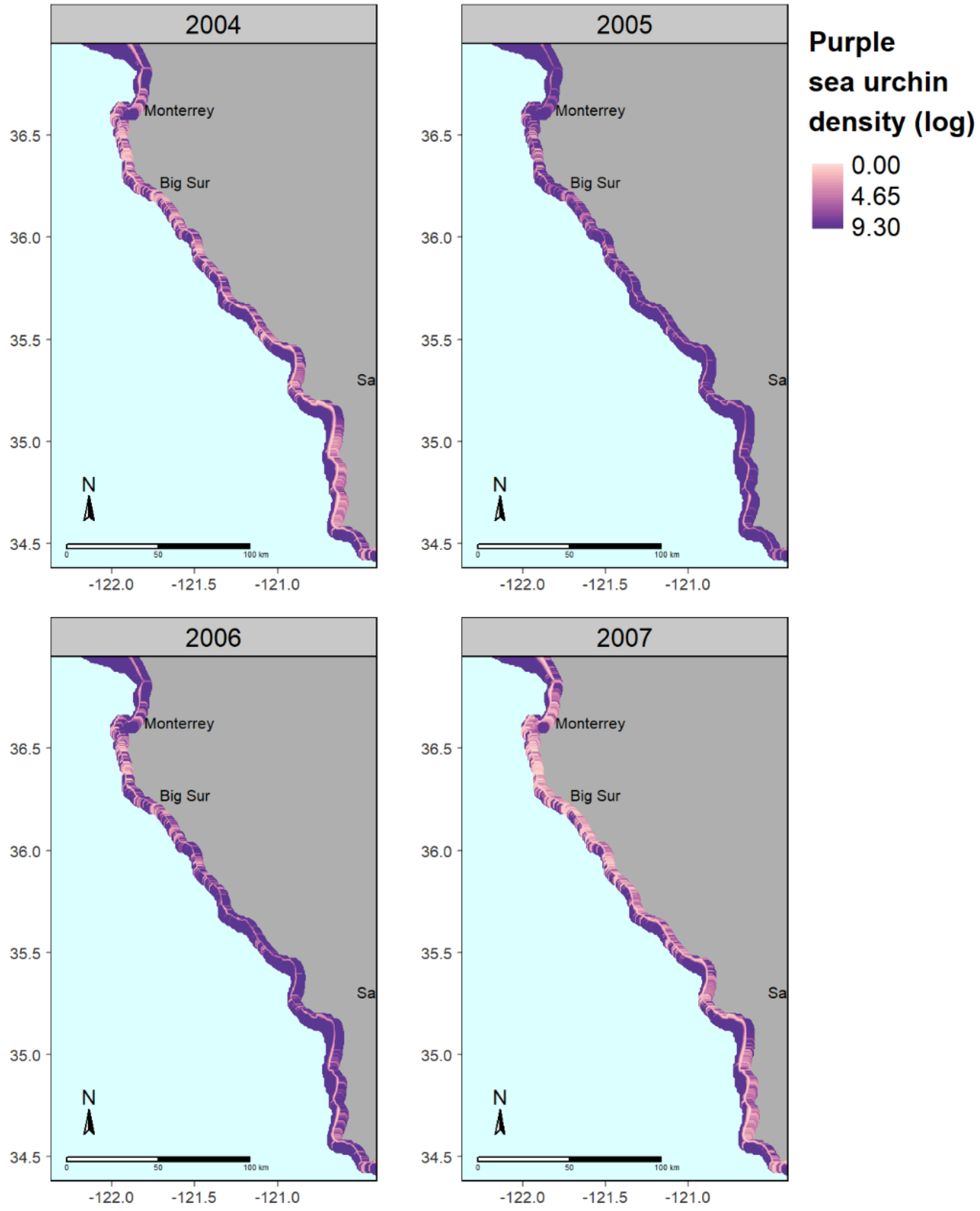

**FIGURE S9.** Predicted density of purple sea urchins (log urchins/60 m$^2$) in central California from 2004 to 2007.

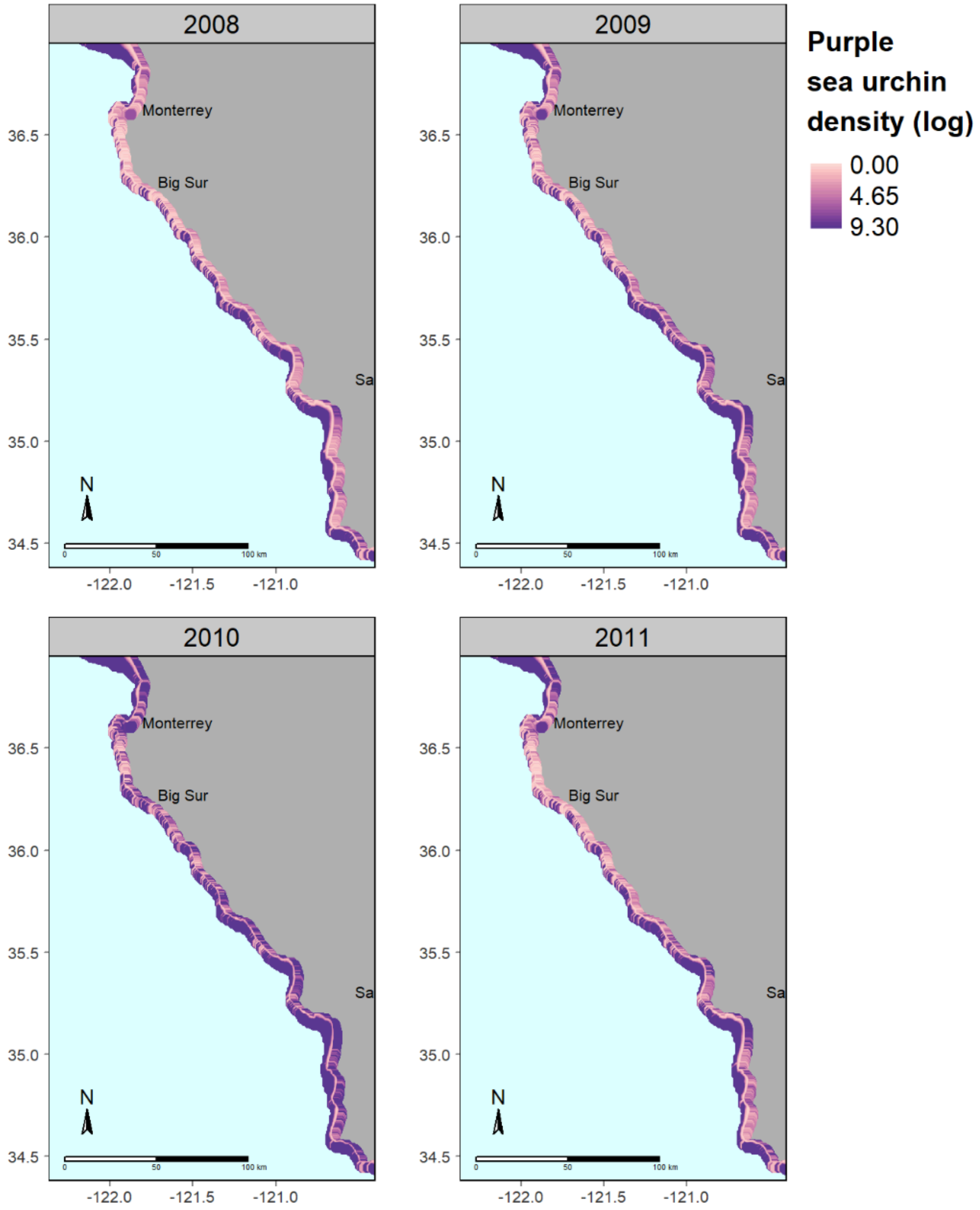

**FIGURE S10.** Predicted density of purple sea urchins (log urchins/60 m$^2$) in central California from 2008 to 2011.

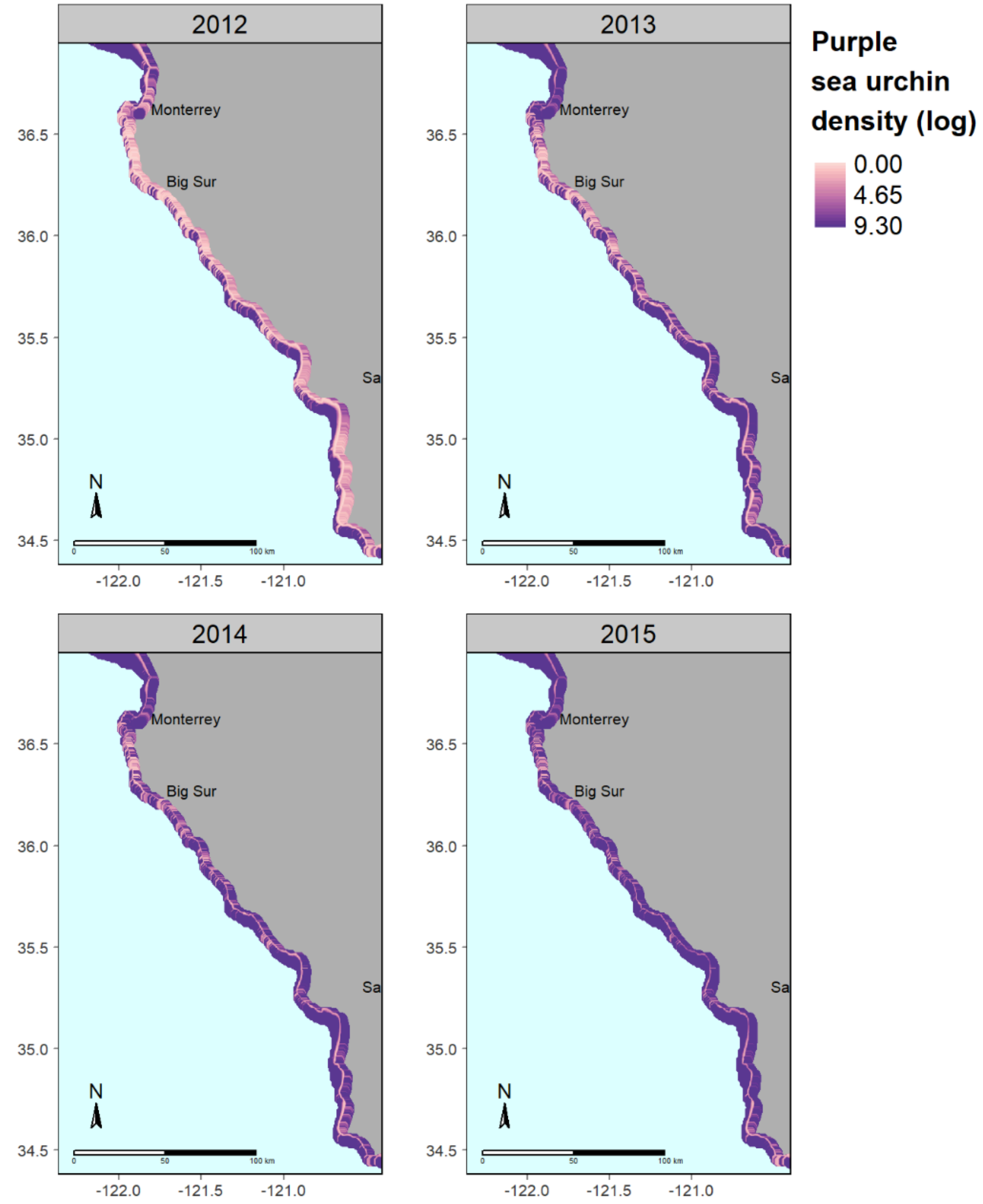

**FIGURE S11.** Predicted density of purple sea urchins (log urchins/60 m$^2$) in central California from 2012 to 2015.

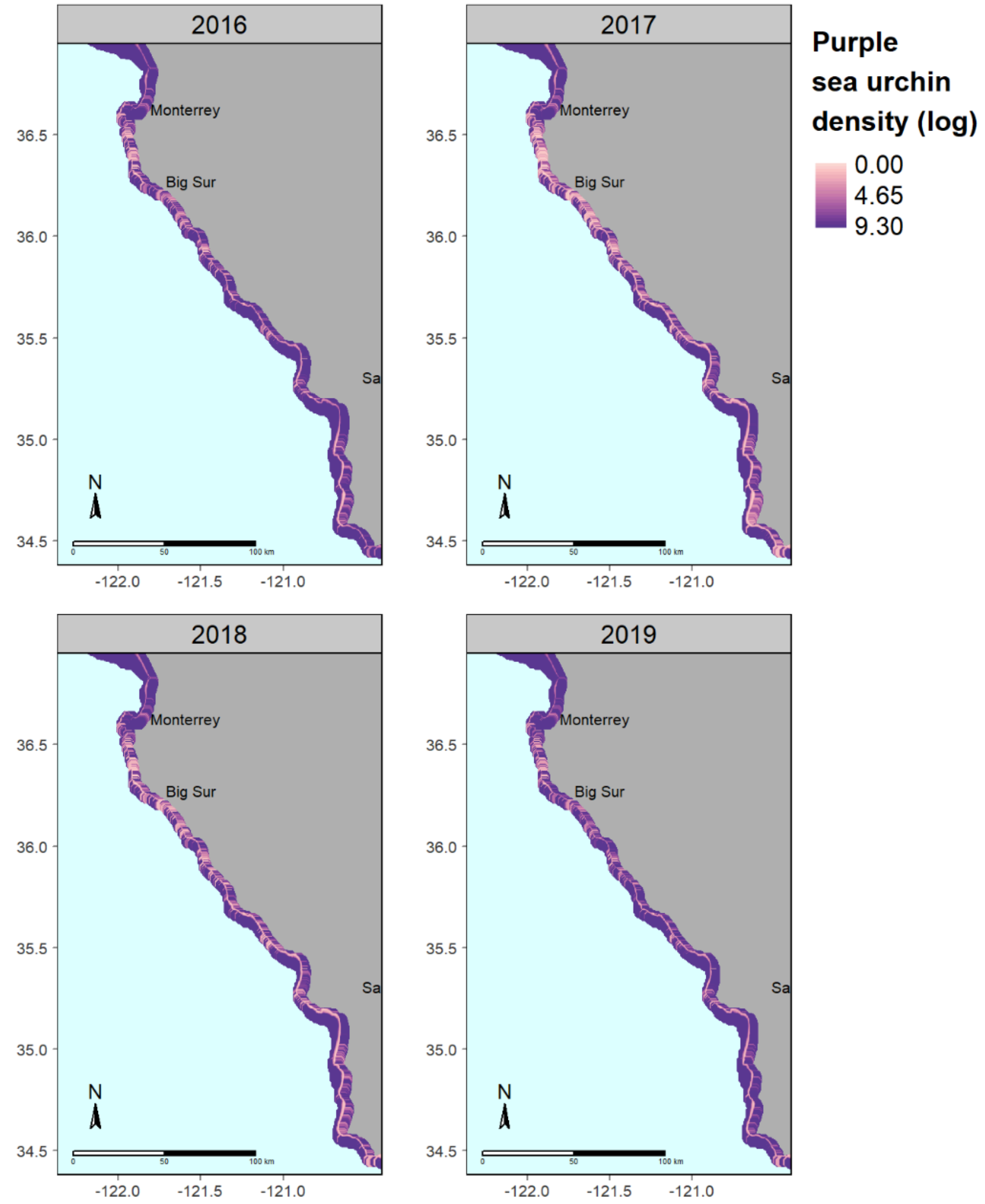

**FIGURE S12.** Predicted density of purple sea urchins (log urchins/60 m$^2$) in central California from 2016 to 2019.

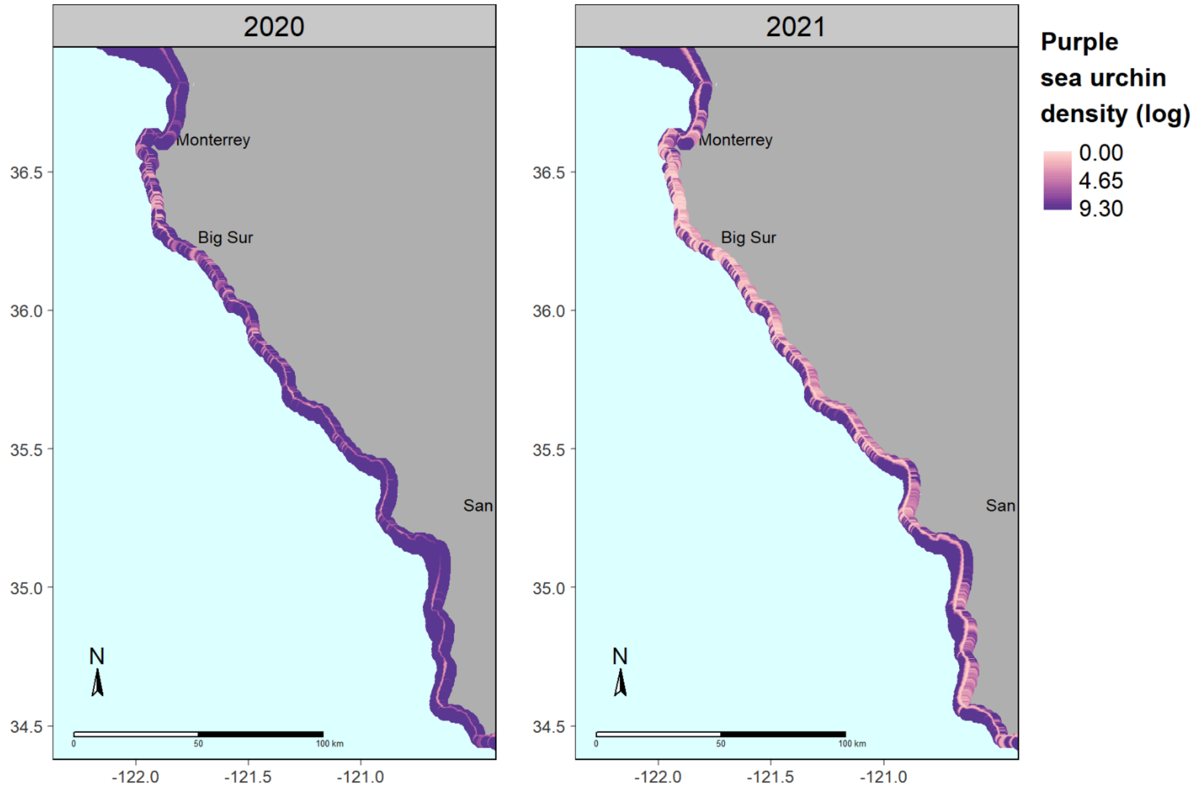

**FIGURE S13.** Predicted density of purple sea urchins (log urchins/60 m$^2$) in central California from 2020 to 2021.

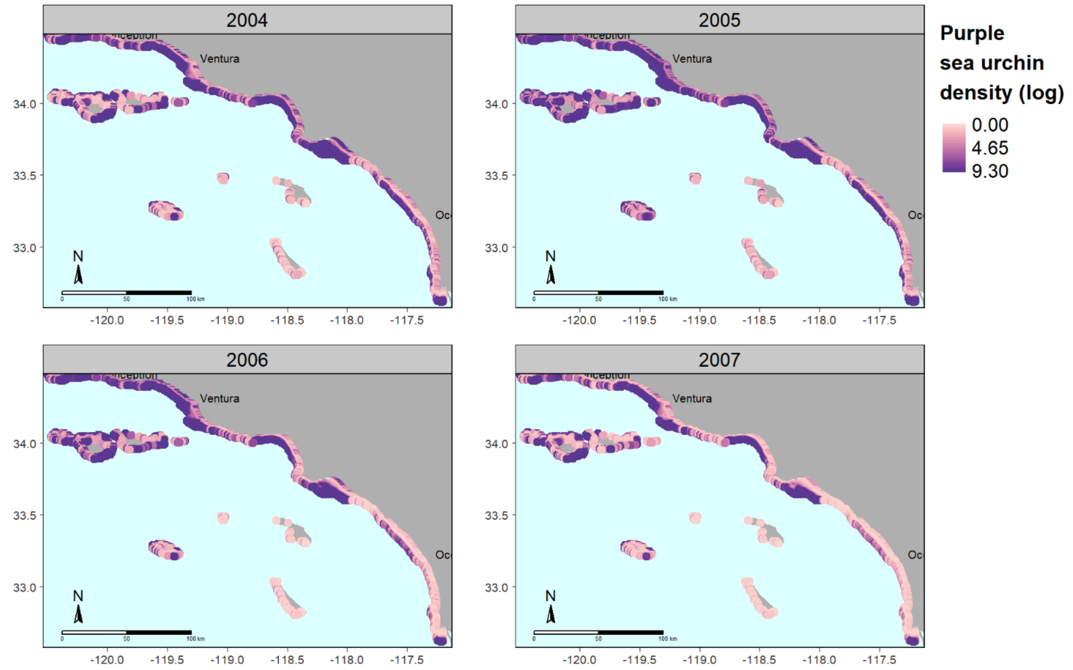

**FIGURE S14.** Predicted density of purple sea urchins (log urchins/60 $m^2$) in southern California from 2004 to 2007.

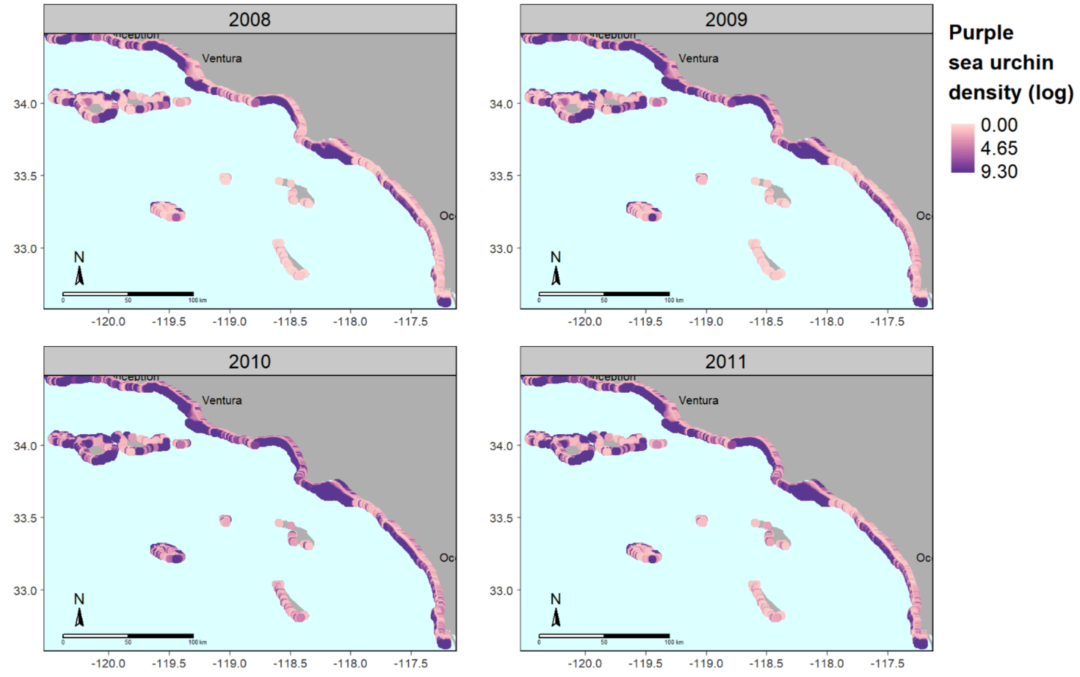

**FIGURE S15.** Predicted density of purple sea urchins (log urchins/60 $m^2$) in southern California from 2008 to 2011.

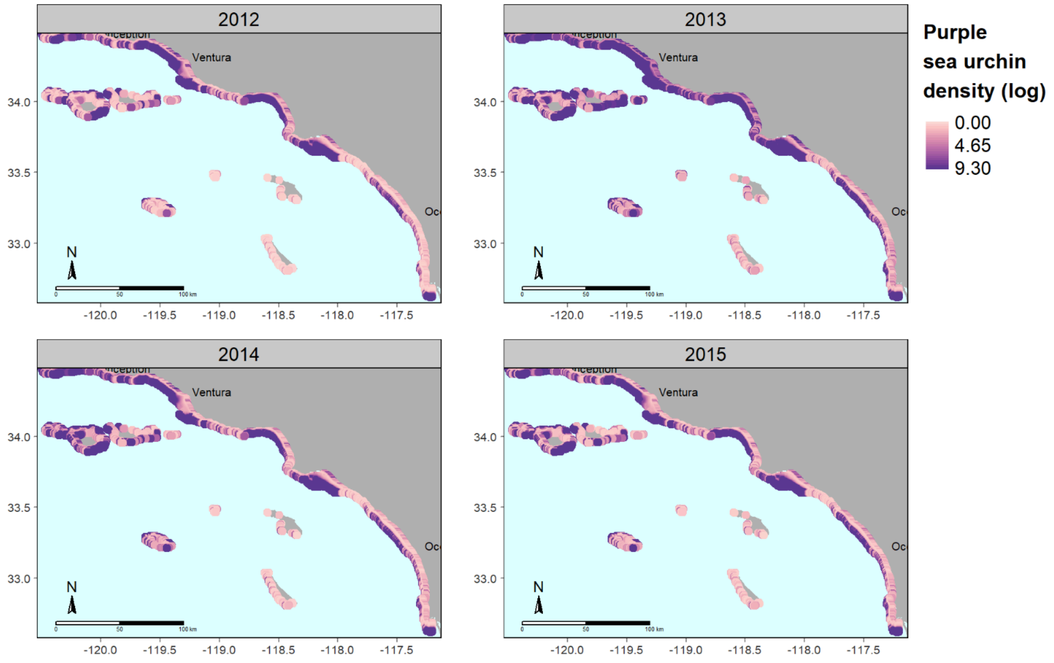

**FIGURE S16.** Predicted density of purple sea urchins (log urchins/60 $m^2$) in southern California from 2012 to 2015.

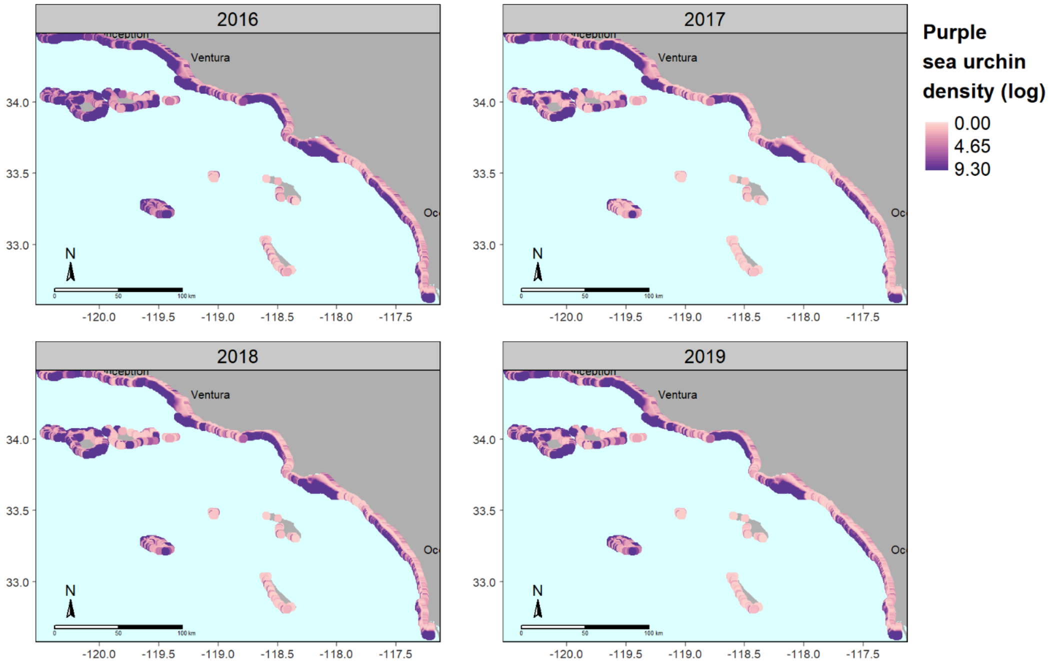

**FIGURE S17.** Predicted density of purple sea urchins (log urchins/60 $m^2$) in southern California from 2016 to 2019.

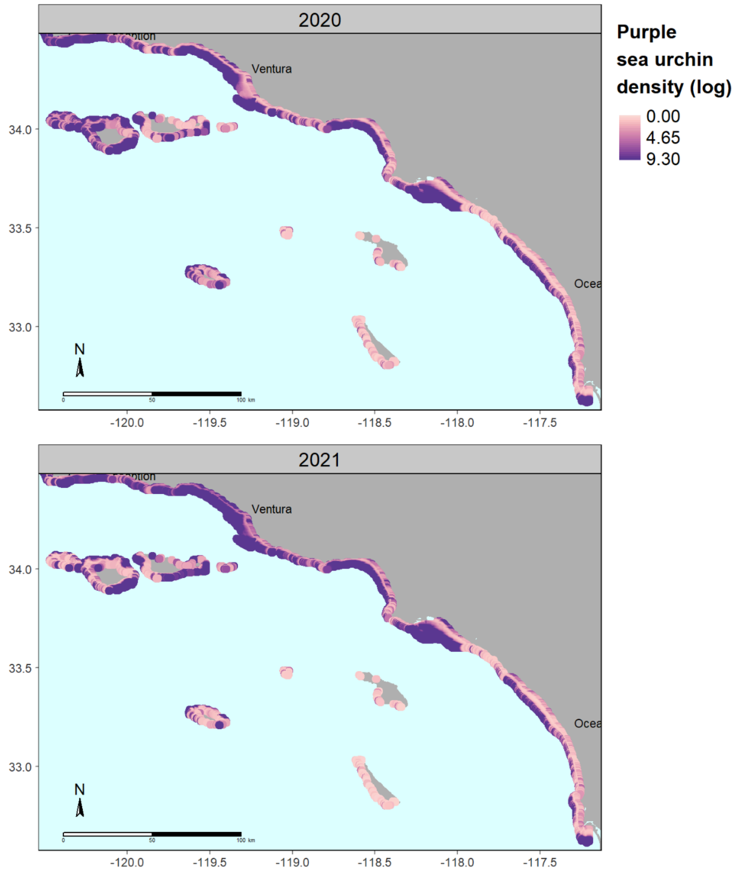

**FIGURE S18.** Predicted density of purple sea urchins (log urchins/60 $m^2$) in southern California from 2020 to 2021.

**Appendix S5**

**Authors:** Anita Giraldo-Ospina, Tom Bell, Mark H. Carr, Jennifer E. Caselle

**Title:** Drivers of spatio-temporal variability in a marine foundation species

**Extended results of bull and giant kelp spatio-temporal models and reconstructed maps of annual bull and giant kelp density from spatio-temporal models (2004-2021)**

**Table S1.** Summary statistics of predictive performance of the GAMs developed to describe the drivers of *N. luetkeana* density dynamics in northern and central California. Several models were tested for each kelp species to identify if drivers varied with region or described the density across their entire distribution in California. Higher $r^2$ and deviance explained indicate improved model fit. Asterix (*) denotes the models selected to project kelp density maps based on their higher $r^2$ and deviance explained.

| **Species** | **Region** | **Model** | **$r^2$** | **Deviance explained** | **AIC** |
|---|---|---|---|---|---|
| *N. luetkeana* * | North and Central | Minimum nitrate + mean temperature upwelling + mean wave height + max orbital velocity (log) + mean NPP + depth + density of purple sea urchins (log) | 0.58 | 63.3% | 5098.27 |

| | | | | | |
|---|---|---|---|---|---|
| *N. luetkeana* | North and Central<br><br>(Best model from North Coast) | Maximum nitrate +<br><br>mean temperature upwelling +<br><br>mean wave height +<br><br>max orbital velocity (log) +<br><br>mean NPP +<br><br>depth +<br><br>density of purple sea urchins (log) | 0.58 | 63.28% | 6883.98 |
| *N. luetkeana* | North and Central<br><br>(Best model from Central Coast) | Minimum nitrate +<br><br>minimum temperature anomaly +<br><br>mean wave height +<br><br>max orbital velocity (log) +<br><br>mean NPP +<br><br>depth +<br><br>density of purple sea urchins (log) | 0.57 | 62.99% | 6904.92 |
| *N. luetkeana* | North | Maximum nitrate +<br><br>mean temperature upwelling +<br><br>mean wave height +<br><br>max orbital velocity (log) +<br><br>mean NPP +<br><br>depth +<br><br>density of purple sea urchins (log) | 0.69 | 66.66% | |

| | | | | |
|---|---|---|---|---|
| *N. luetkeana* | Central | Minimum nitrate +<br>minimum temperature anomaly +<br>mean wave height +<br>max orbital velocity (log) +<br>mean NPP +<br>depth +<br>density of purple sea urchins (log) | 0.59 | 67.31% |

**Table S2.** Summary statistics of predictive performance of the GAMs developed to describe the drivers of *M. pyrifera* density dynamics in California. Several models were tested for giant kelp to identify if drivers varied with region or described the density across their entire distribution in

California. Higher $r^2$ and deviance explained indicate improved model fit. Asterix (*) denotes the models selected to project kelp density maps based on their higher $r^2$ and deviance explained.

| **Species** | **Region** | **Model** | **$r^2$** | **Deviance explained** |
|---|---|---|---|---|
| *M. pyrifera* * | Central and South-west | Days 4N (log) +<br>days 21C (sqrt) +<br>mean wave height +<br>max orbital velocity (log) +<br>mean NPP +<br>depth +<br>density of purple sea urchins (log) +<br>Previous year spores (log) | 0.59 | 45.03% |
| *M. pyrifera* * | South-east | Maximum nitrate summer anomaly (log) +<br>days 21C (log) +<br>mean wave height (log) +<br>mean NPP +<br>depth +<br>density of purple sea urchins (log) +<br>Previous year spores (log) | 0.62 | 51.44% |
| *M. pyrifera* | Central | Depth +<br>density of purple sea urchins (log) + | 0.57 | 47.66% |

| | | | | |
|---|---|---|---|---|
| | | Previous year spores (log) | | |
| *M. pyrifera* | South-west | Mean nitrate upwelling +<br>days 18C (log) +<br>mean temperature upwelling +<br>max orbital velocity +<br>minimum NPP upwelling +<br>mean depth +<br>density of purple sea urchins (log) +<br>Previous year spores (log) | 0.54 | 40.76% |
| *M. pyrifera* | Central and South | Days 4N (log) +<br>days 21C (sqrt) +<br>mean wave height +<br>depth +<br>density of purple sea urchins (log) +<br>Previous year spores (log) | 0.57 | 45.06% |
| *M. pyrifera* | Central, South-west and South-east | Days 4N (log) +<br>days 21C (sqrt) +<br>mean wave height +<br>depth +<br>density of purple sea urchins (log) +<br>Previous year spores (log) | 0.57 | 45.06% |

| | | | | |
|---|---|---|---|---|
| *M. pyrifera* | South-west and South-east | Days 3N (log) + <br>days 16C (sqrt) + <br>mean wave height + <br>maximum orbital velocity (log) + <br>depth + <br>density of purple sea urchins (log) + <br>Previous year spores (log) | 0.56 | 44.13% |

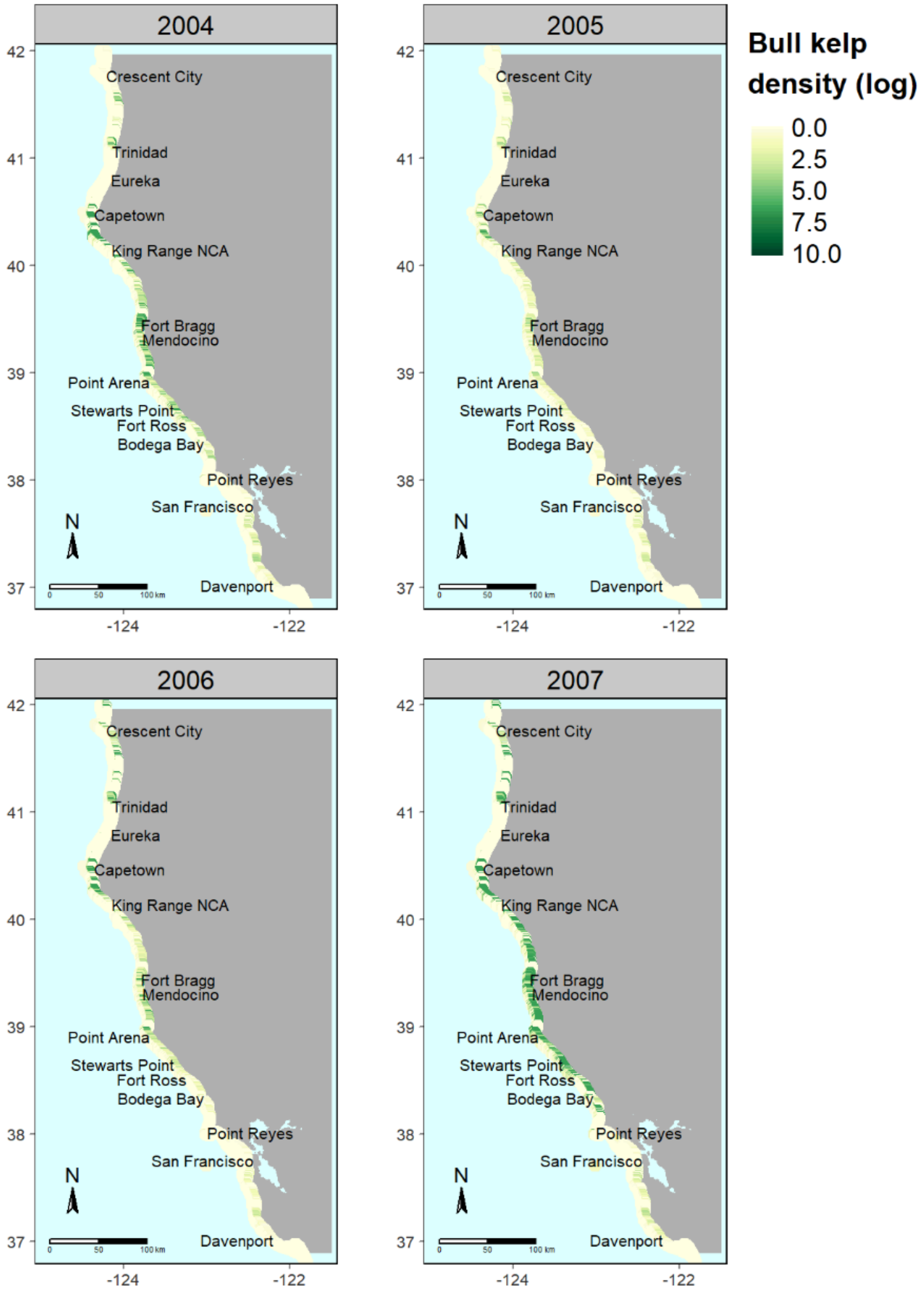

**FIGURE S1.** Predicted density of bull kelp (log individuals/60 $m^2$) in northern California from 2004 to 2007.

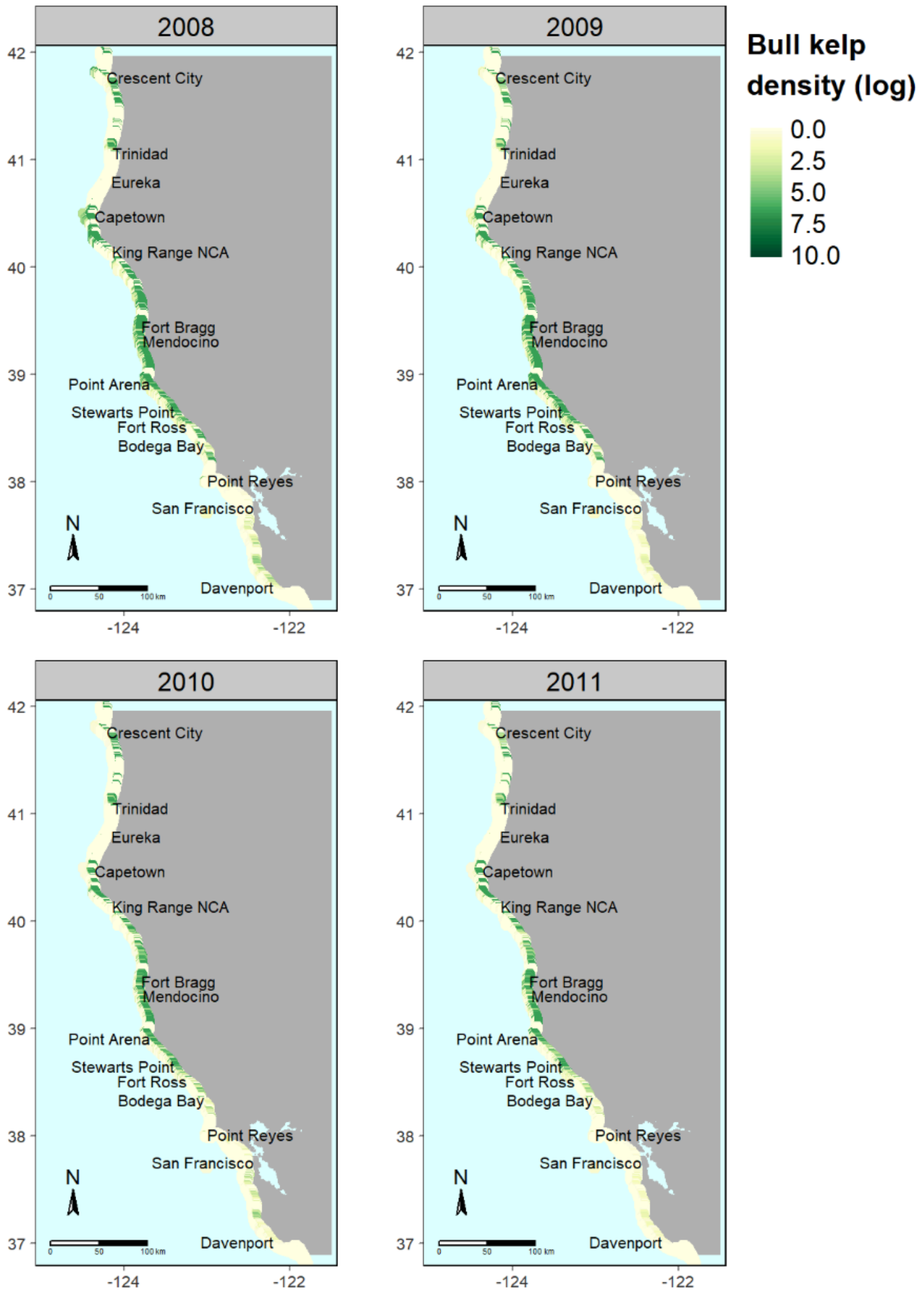

**FIGURE S2.** Predicted density of bull kelp (log individuals/60 $m^2$) in northern California from 2008 to 2011.

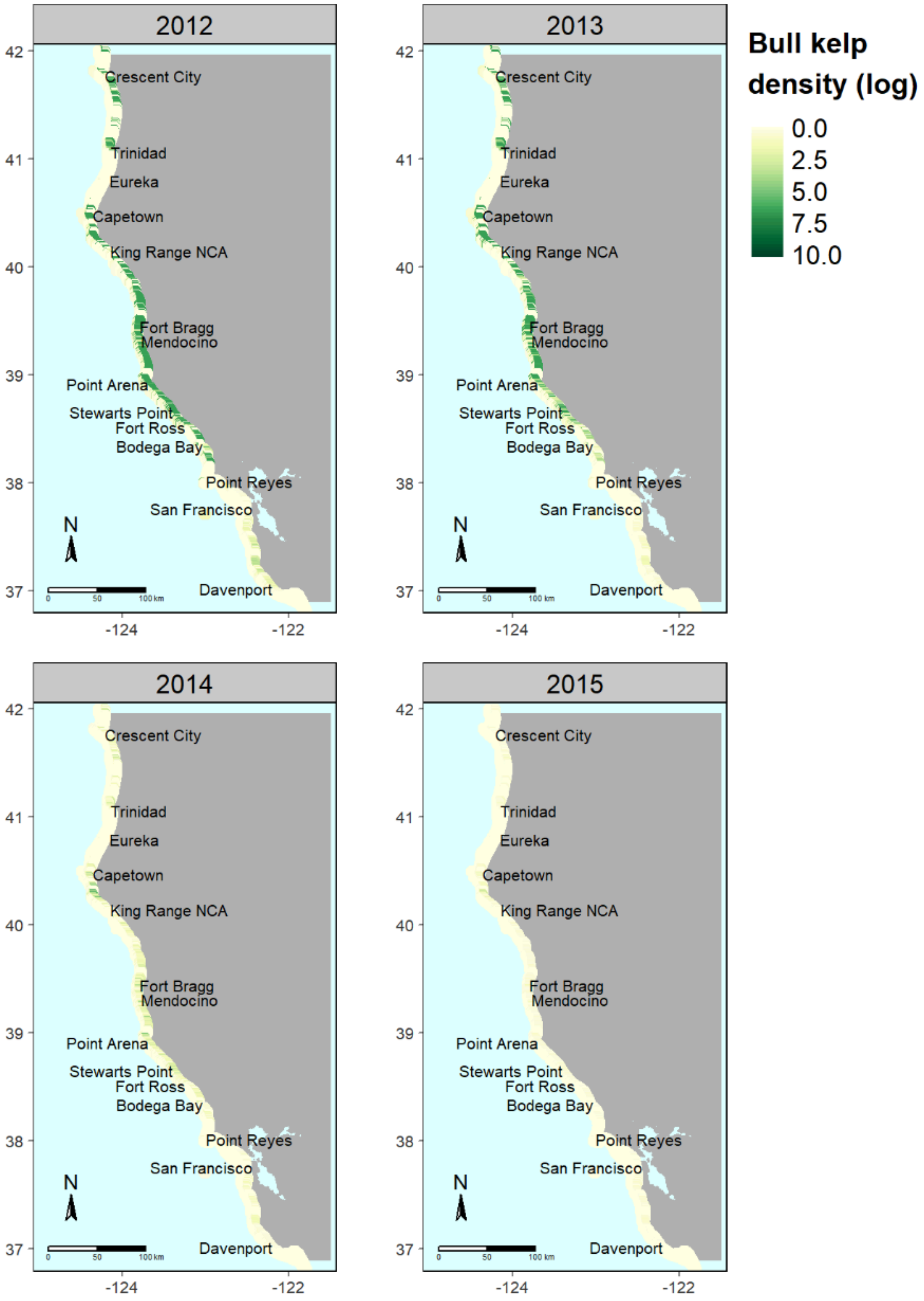

**FIGURE S3.** Predicted density of bull kelp (log individuals/60 $m^2$) in northern California from 2012 to 2015.

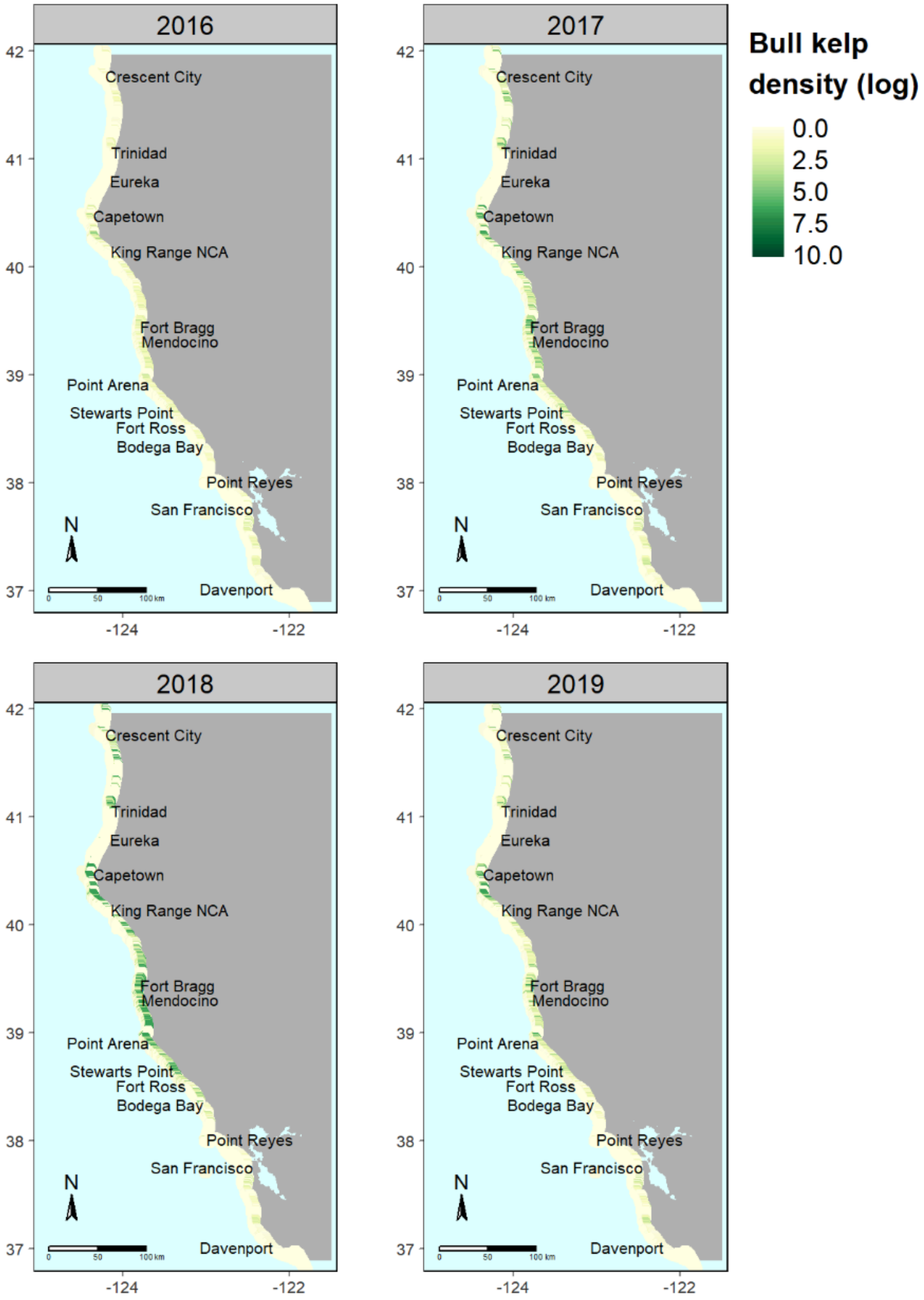

**FIGURE S4.** Predicted density of bull kelp (log individuals/60 $m^2$) in northern California from 2016 to 2019.

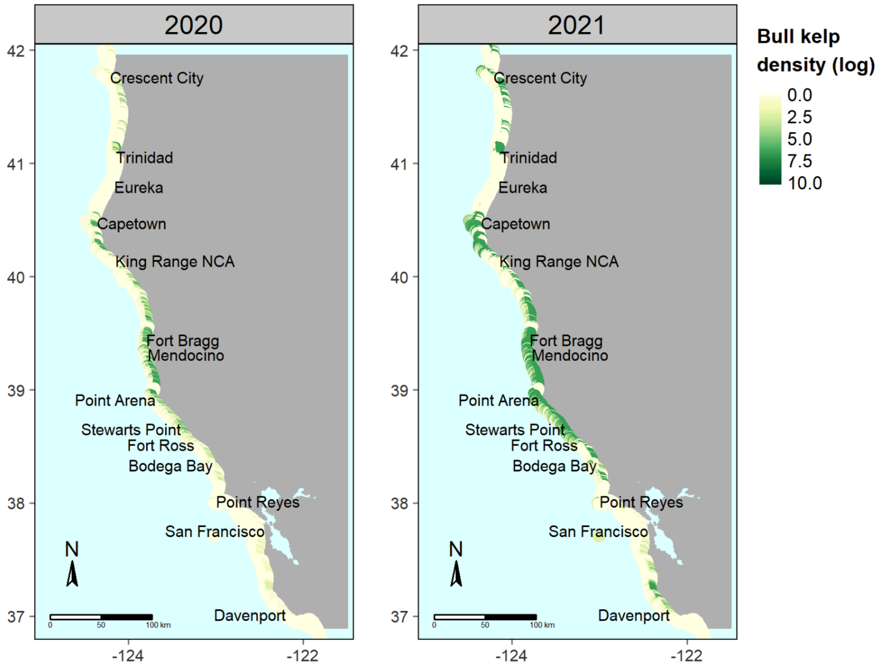

**FIGURE S5.** Predicted density of bull kelp (log individuals/60 $m^2$) in northern California from 2020 to 2021.

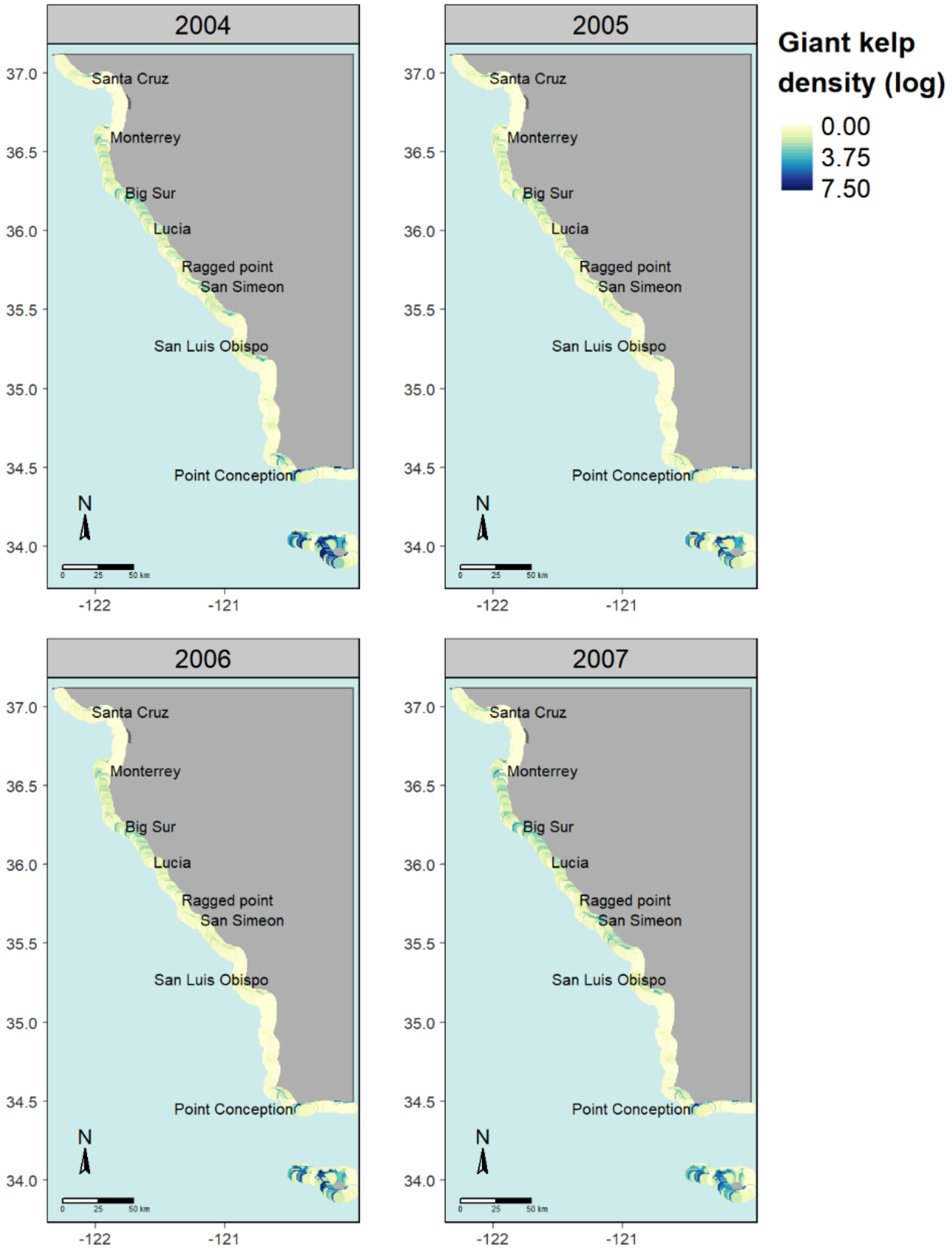

**FIGURE S6.** Predicted density of giant kelp (log stipes/60 $m^2$) in central California and outer Channel Islands (San Miguel and Santa Rosa) from 2004 to 2007.

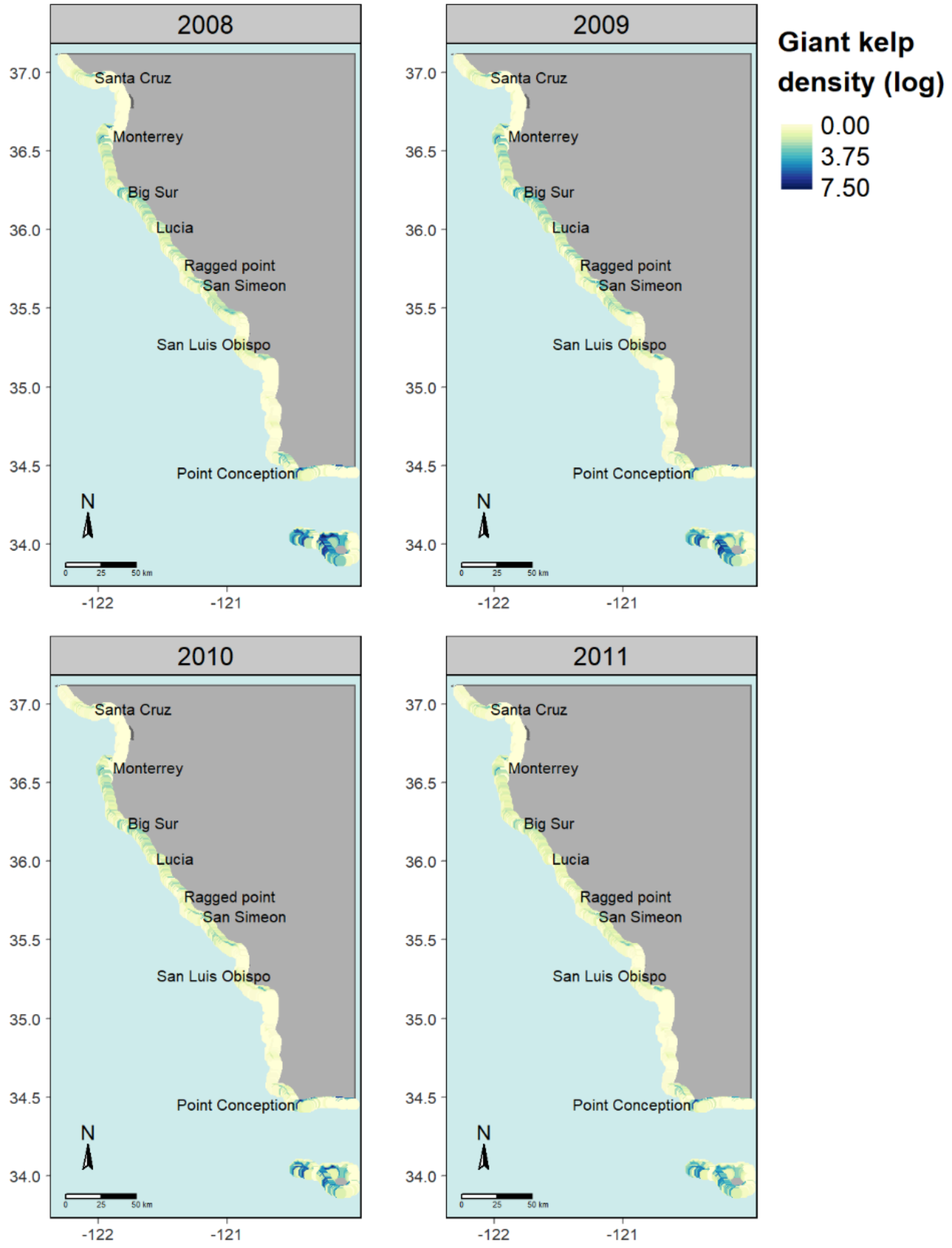

**FIGURE S7.** Predicted density of giant kelp (log stipes/60 $m^2$) in central California and outer Channel Islands (San Miguel and Santa Rosa) from 2008 to 2011.

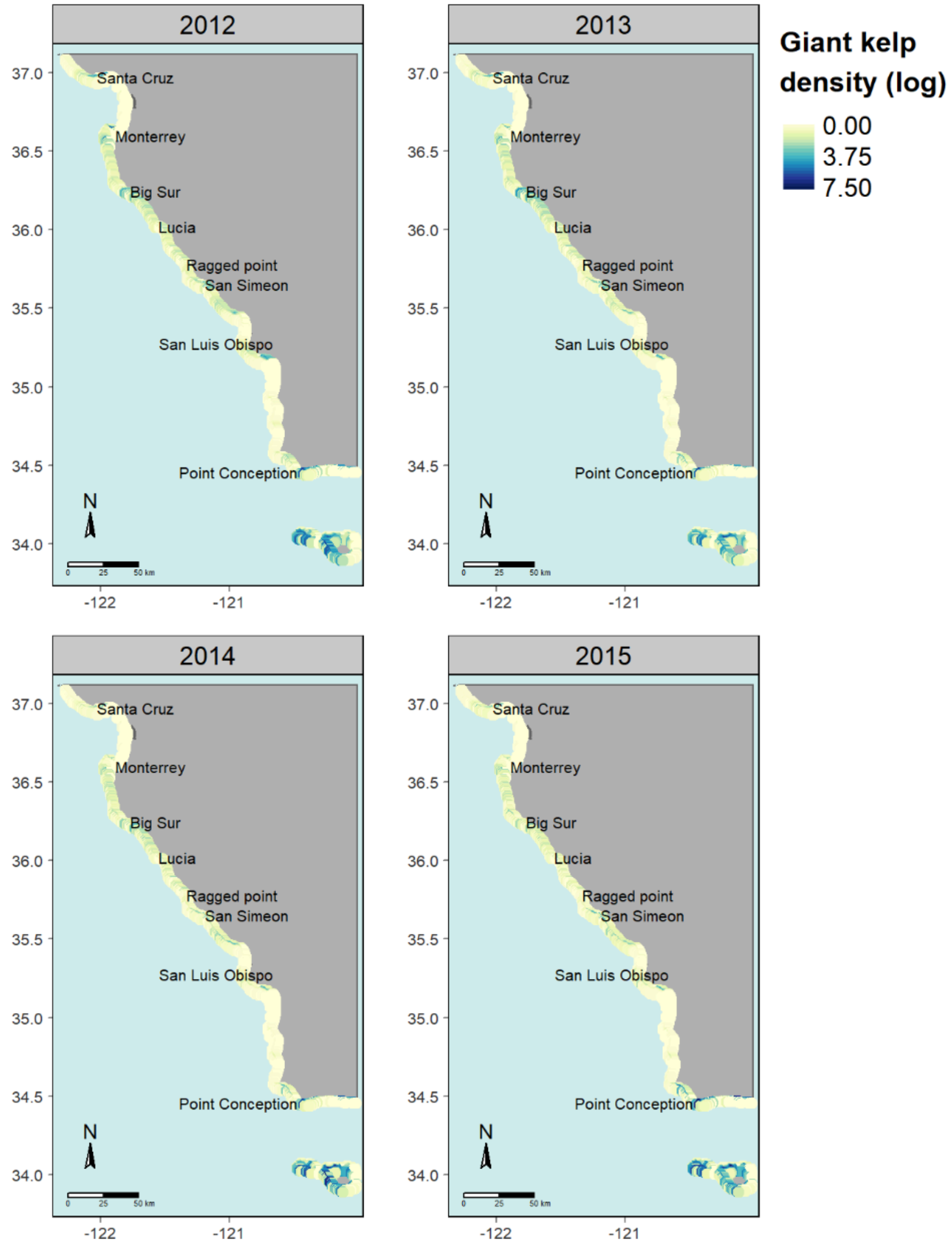

**FIGURE S8.** Predicted density of giant kelp (log stipes/60 $m^2$) in central California and outer Channel Islands (San Miguel and Santa Rosa) from 2012 to 2015.

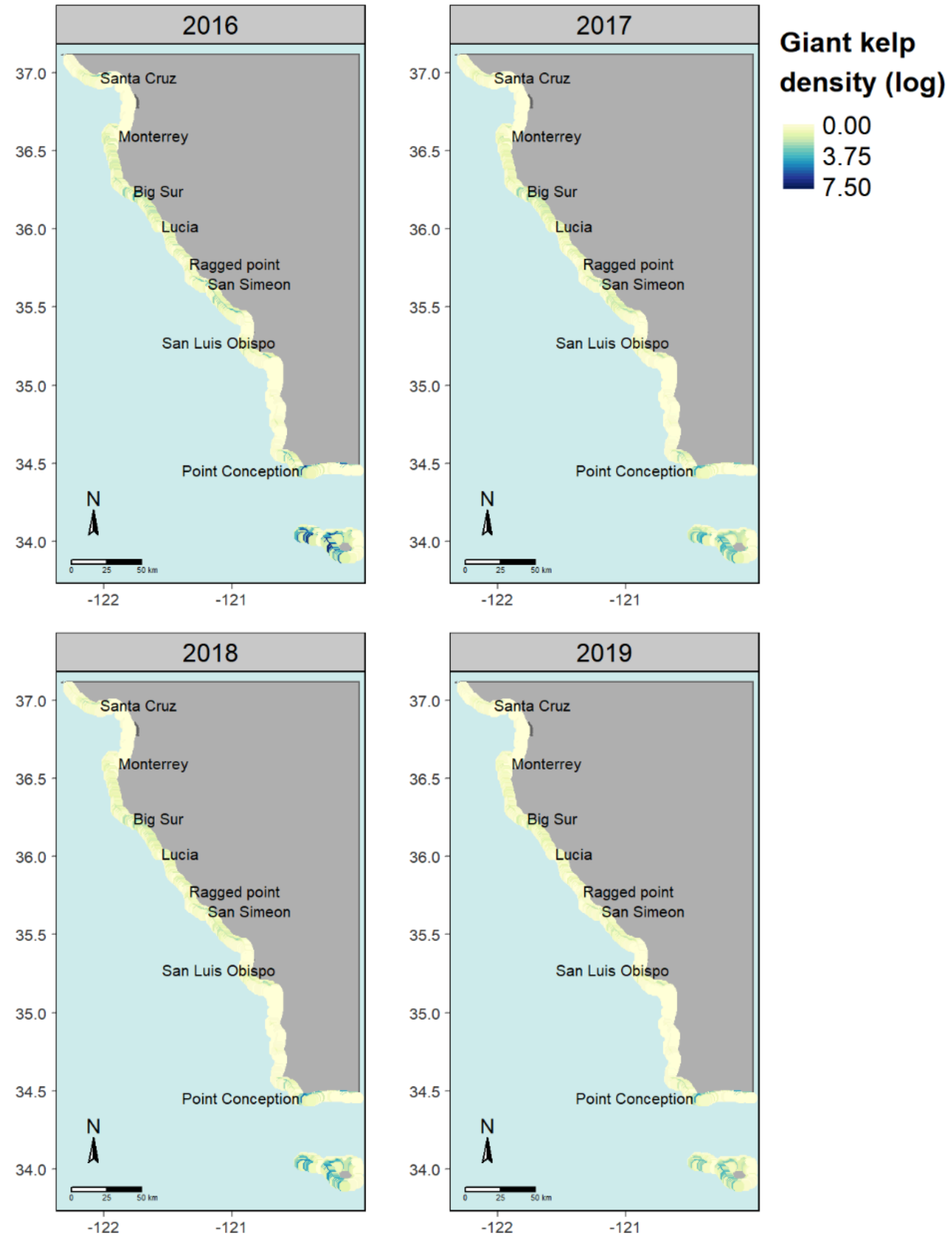

**FIGURE S9.** Predicted density of giant kelp (log stipes/60 $m^2$) in central California and outer Channel Islands (San Miguel and Santa Rosa) from 2016 to 2019.

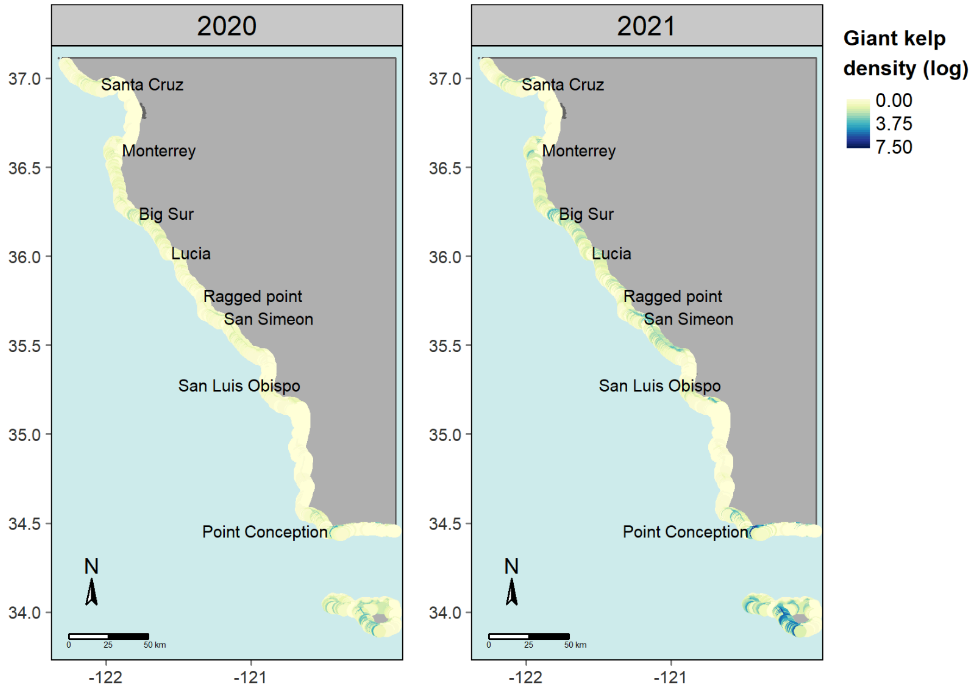

**FIGURE S10.** Predicted density of giant kelp (log stipes/60 $m^2$) in central California and outer Channel Islands (San Miguel and Santa Rosa) from 2020 to 2021.

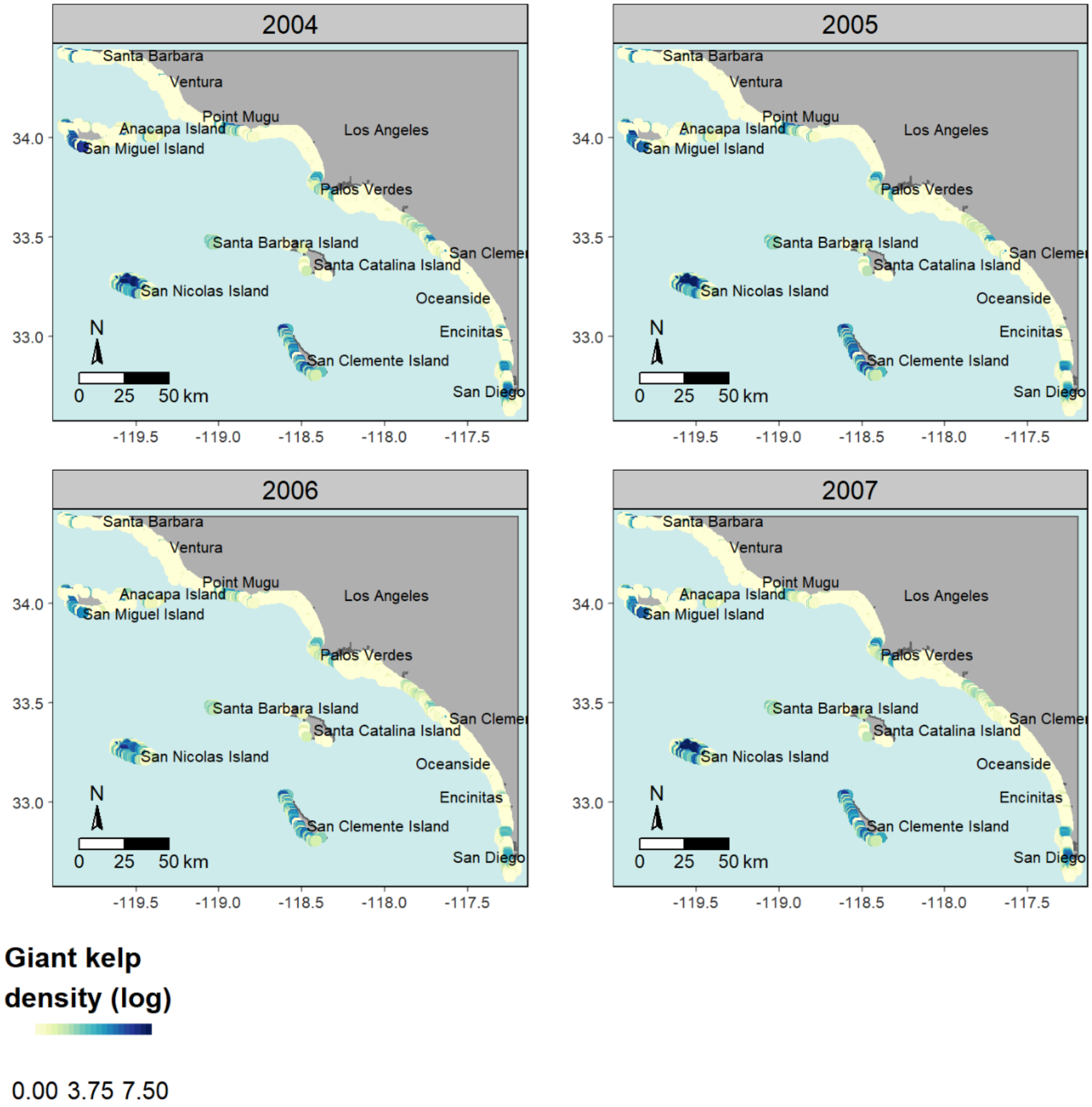

**FIGURE S11.** Predicted density of giant kelp (log stipes/60 $m^2$) in southern California from 2004 to 2007.

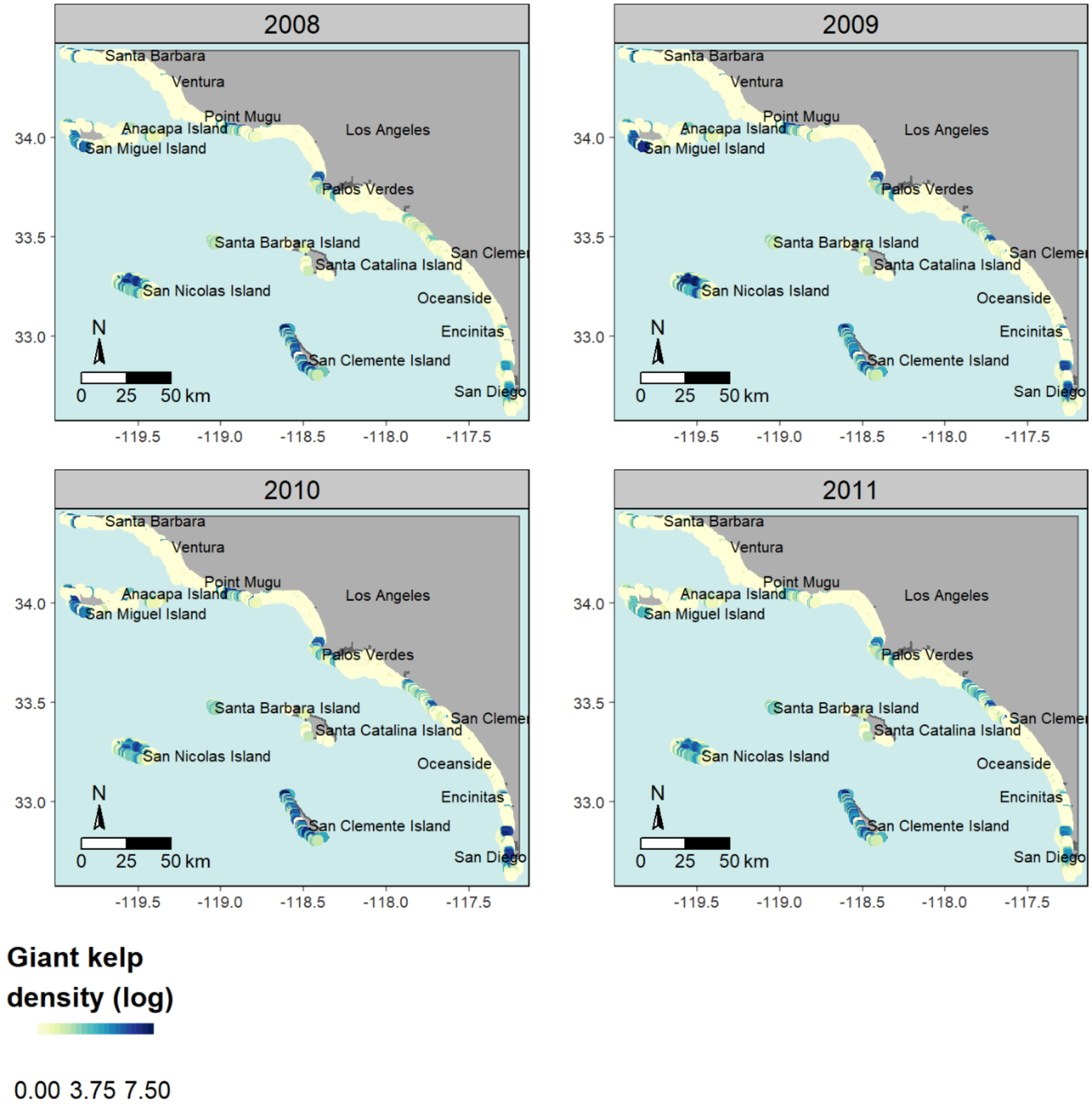

**FIGURE S12.** Predicted density of giant kelp (log stipes/60 $m^2$) in southern California from 2008 to 2011

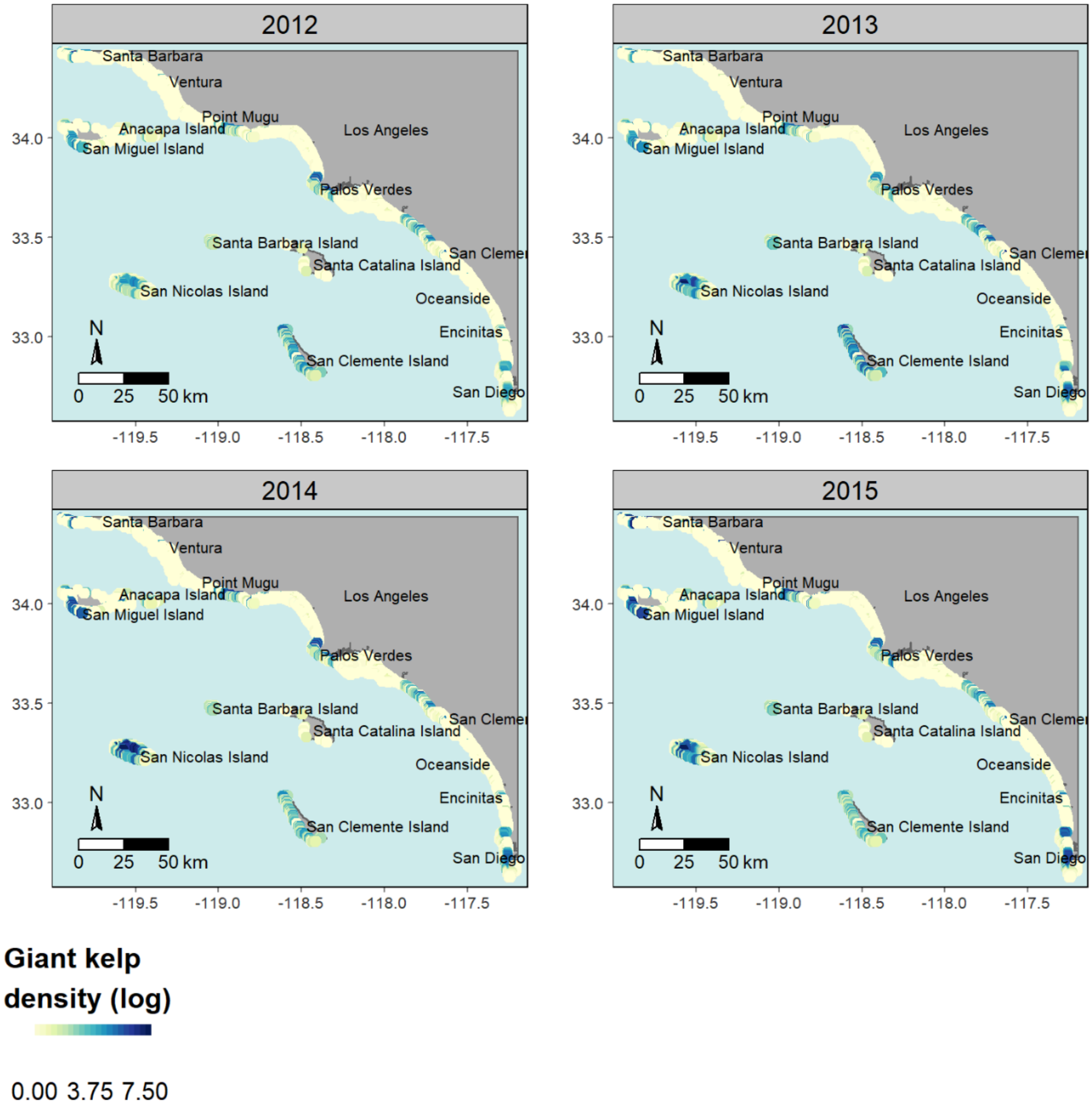

**FIGURE S13.** Predicted density of giant kelp (log stipes/60 $m^2$) in southern California from 2012 to 2015.

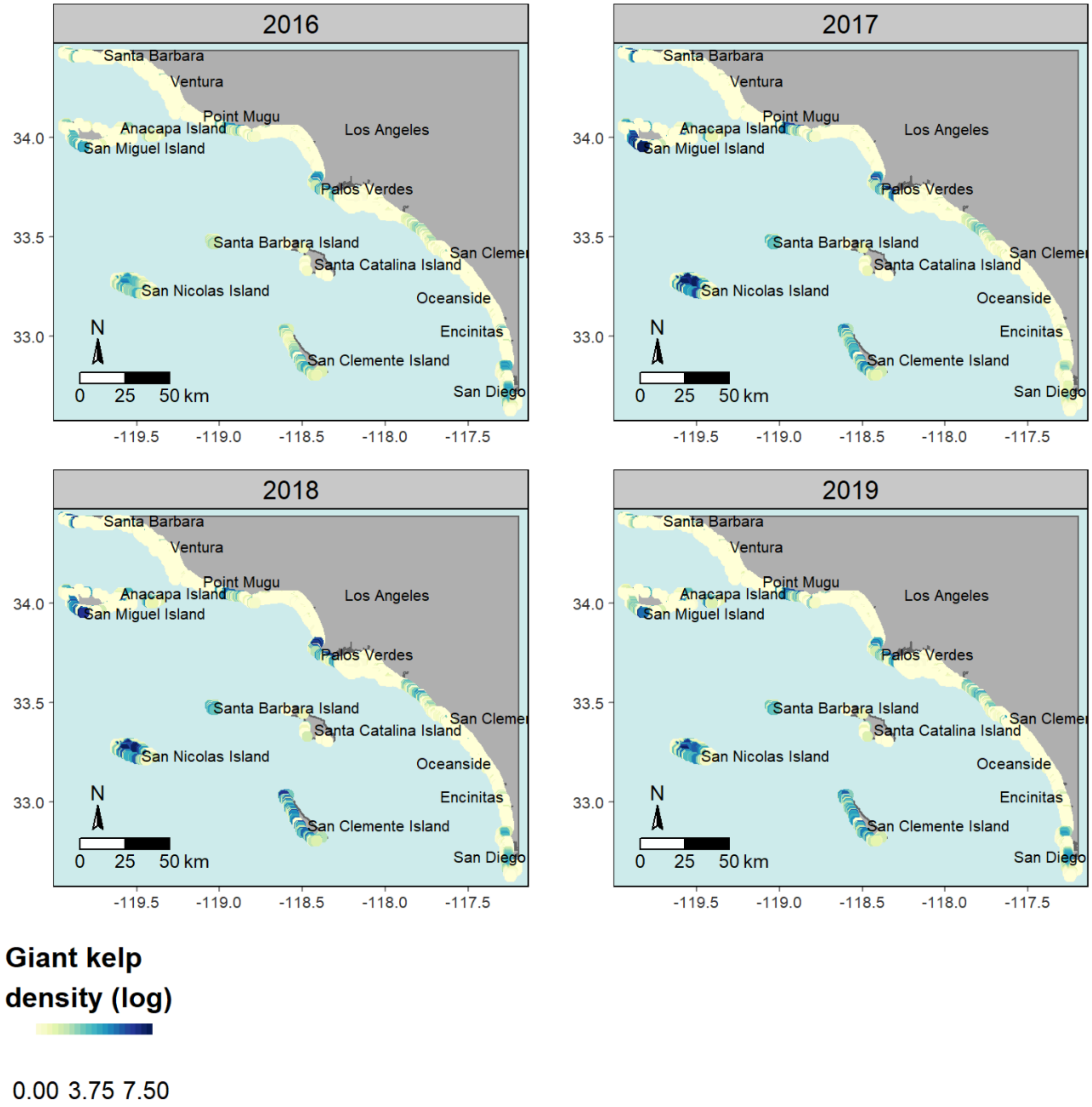

**FIGURE S14.** Predicted density of giant kelp (log stipes/60 m$^2$) in southern California from 2016 to 2019.

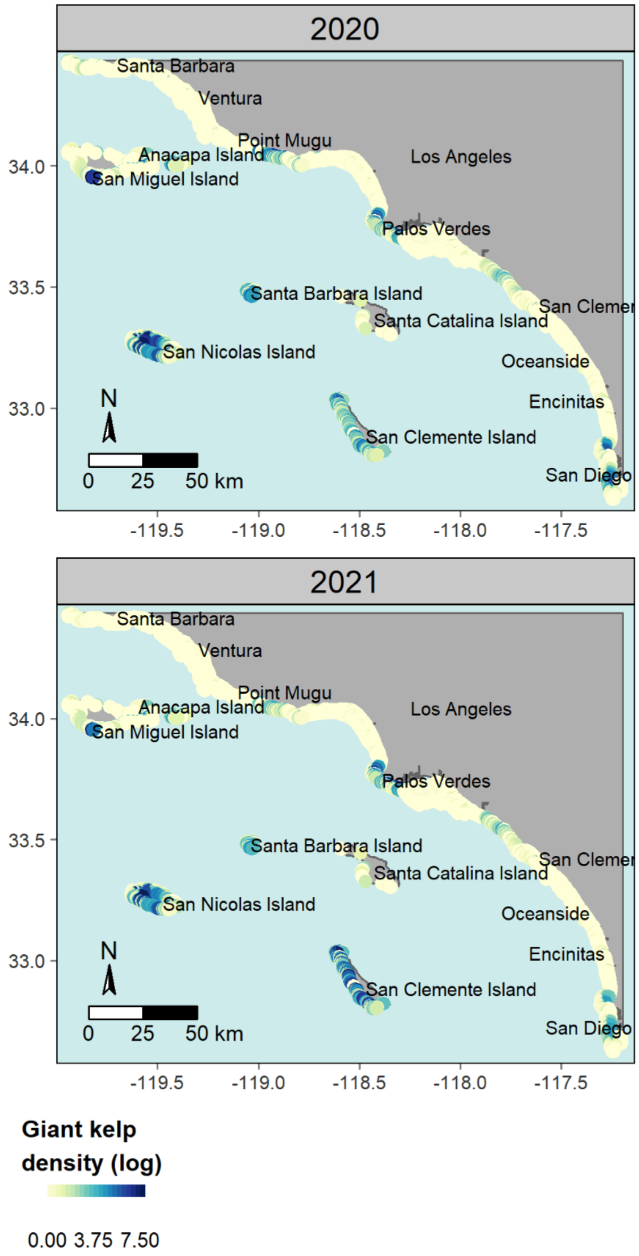

**FIGURE S15.** Predicted density of giant kelp (log stipes/60 $m^2$) in southern California from 2020 to 2021.
